# An ecological, phenotypic and genomic survey of duckweeds with their associated aquatic environments in the United Kingdom

**DOI:** 10.1101/2024.08.14.607898

**Authors:** Kellie E. Smith, Laura Cowan, Paulina Flis, Chris Moore, Matthew Heatley, Carlos A. Robles-Zazueta, Adam Lee, Levi Yant

## Abstract

The duckweeds feature global distributions and diverse applications in phytoremediation and nutrition, as well as use in fundamental studies of development. Existing collections have minimal environmental data linked to natural habitats. Thus, there is a lack of understanding of natural variation in the context of native habitats. Here, a novel collection of 124 duckweed accessions from 115 sites across the United Kingdom were characterised by genome sequencing and ionomics. In nutrient-replete conditions all accessions hyperaccumulated P, K, Mg and Ca. Local but not large-scale associations were revealed between elemental composition of duckweed in common, replete conditions and native water profiles. *Lemna minor* was the most prevalent species in the UK, with a closely related hybrid *L. japonica* frequently found in waters with higher micronutrient concentrations. Invasive *L. minuta* was common in the southern and midland regions, but restricted in Scotland. *Lemna* accessions accumulated heavy metal contaminants typically together with macronutrients, suggesting phytoremediation potential, but some limitations as food. Furthermore, monitoring the ecological interactions between native, hybrid and invasive *Lemna* species should be ongoing in the interest of biodiversity.

## Introduction

Duckweeds (Lemnaceae) represent some of the fastest growing flowering plants in the world (Sree et al., 2015; Ziegler et al., 2015) and are powerful models for studies in development (Ware *et al*.; 2023), bioremediation and ecotoxicology (Laird and Barks, 2018). There are at least thirty-six species of duckweed, consisting of simple stem-leaf structures called fronds in the rootless *Wolffia* and *Wolffiella* genera, to early diverged root-bearing genera *Spirodela*, *Landoltia* and *Lemna* (Lam and Michael, 2022; Ware et al., 2023). Duckweed have increasing roles in wastewater purification through uptake of excessive nutrients, metals and toxic elements leached from industrial and agricultural activities (Landesman et al., 2010; Ekperusi et al., 2019). Moreover, other varieties are emerging as food sources when grown hydroponically and axenically in vertical farms, providing comparable protein and nutritional contents to wheat (Appenroth et al., 2017; Xu et al., 2023). However, applications development requires novel duckweed clones with improved understanding of environmental interactions (Barton, 2024). For context, traditional crops like wheat and maize have been optimized for abiotic stress resilience and quality traits by assessing wild relatives in their natural habitats (Zhang et al., 2017; Reynolds and Braun, 2022). Adopting similar approaches with new duckweed germplasms may unlock development of future food and phytoremediation applications.

Presently duckweed collections are generally domesticated to axenic artificial conditions (Sree and Appenroth, 2020). Collection dates, original environmental data and genome sequencing for existing clones are largely unavailable, limiting studies of accession optimisation to bioremediation or food production. Ecological characterisation at regional scales have been performed with some collections: from European (Kirjakov and Velichkova, 2016), Middle Eastern (Friedjung Yosef et al., 2022; Taghipour et al., 2022) and Asian accessions (Xu et al., 2015; Chen et al., 2022; Kadono and Iida, 2022; Tran et al., 2022). These works have uncovered various invasive and hybrid *Lemna (L.)* species. Within Europe, four invasive alien species have been identified, with *L. minuta* dominating (Lansdown, 2008; Fedoniuk et al., 2022; GBIF.org, 2022). However, characterisation is limited in the UK due to paucity of clones with only four classified as *L. minuta* species (Lam, 2018), a species that opportunistically outcompetes native *L. minor* under high nutrient and light conditions with potentially highly destructive ecological consequences (Njambuya et al., 2011; Ceschin et al., 2016; Paolacci et al., 2016; Paolacci et al., 2018a; Paolacci et al., 2018b).

Clear temporal and spatial patterns in *L. minuta* dispersal have been determined which drive species invasion fronts (Ceschin *et al*., 2018a). For example, avian species are an important vector of dispersal (Coughlan *et al*., 2015; Silva *et al*., 2018) whereas, increasing ambient temperature and nutrient availability promotes invasive *L. minuta* growth (Njambuya *et al*. 2011, Peeters *et al*., 2013). From initial invasion fronts upon European Atlantic coasts in the 1960s, *L. minuta* is now firmly established in the UK (Ceschin *et al*., 2018a). Negative impacts of *L. minuta* dense infestation include thick mat formations which decrease light penetration, pH and oxygenation into water bodies, thereby reducing native biodiversity and causing problems for aquatic fauna (Janes et al., 1996; Ceschin et al., 2019), although *L. minor* can also form dense mats and dominate aquatic environments in optimal conditions. Wetland habitats in the UK of high conservation status are now threatened by hyper-eutrophication, ecosystem imbalance and duckweed invasion (Feller *et al*., 2024). It is therefore timely to conduct regional surveys of both native and invasive duckweed species in wild wetlands with a view to assessing species specific adaptations in these environments.

Additionally, particularly ‘adaptable’ or ‘extremophile’ duckweed have great promise for the development of phytoremediation and food applications. Consideration of the plant ‘ionome’ refers to its whole-tissue or organismal levels of macro-, micro-nutrients and trace minerals (Salt et al., 2008). The applications of ionomics ranges from assessments of nutrient uptake and soil/water relations to understanding the nutritional composition of food and biofortification of crops. In duckweed, clones of *L. minor* and *Wolffia globosa* hyperaccumulate over 1 g/kg dry weight of heavy metals such as Cd, Cu and As and may have phytoremediation potential (Zayed et al., 1998; Zhang et al., 2009). From a worldwide collection, *L. yungensis* clones displayed local-scale variation in macronutrients Mg, S and Mn (Smith et al., 2024b). This suggests that nutrient uptake is linked to highly specific adaptations to micro-habitats. However, there is still a lack of understanding of the scale of variation in either water environments or the attendant accumulation potential of native duckweed accessions.

This paper presents a genomic, ecological and environmental assessment of novel UK duckweed accessions, detailing 115 environments and 124 accessions. We discover elemental variation using ionomics at local scales and document the spread of invasive *L. minuta*, as well as new reports of hybrid species. A common garden experiment with replete nutrient media was used to measure differences in duckweed whole plant tissue ionomes and native environmental water using inductively coupled mass spectrometry (ICP-MS). Overall this work provides a local-scale and UK-wide assessment of duckweed variation and water habitats, providing accessions with promising elemental accumulation profiles with potential for food and phytoremediation applications.

## Results

### Cohort construction and environmental assessment

To assess distributions across fine-to moderate geographic scales, duckweeds were collected from across England, Wales and Scotland. Initially, morphology was used for species determination, with later confirmation by genomic sequencing. Environmental assessment was performed concurrent with plant collections, focusing on water body analysis for elemental composition. Names and descriptions of sites are given in Table S1 and a map for sampling regions presented in Fig. S1. Site locations were chosen as described in methods. The primary latitudinal axis was between 41 sites in southern England and Wales (regions HAS, COR, BRI, NEW) and 37 sites across Scotland (regions ABE, ELG, GLA), giving a total of 103 accessions. The central UK consisted of five sampling regions LAN, BFD, YOR, HUL and MID, yielding a total of 44 duckweed accessions. All accession names, sampling coordinates, dates and characteristics are provided in Table S2A.

### Phenotype-based species identification

Morphological factors were used to determine species membership of accessions, including frond and root characteristics and turion production. Phenotypes were quantified first upon collection and then confirmed during laboratory growth for three years cultivated in controlled growth environments. Principal component analysis (PCA) was used with a subset of UK accessions suspected to be different species to discriminate species clusters based on morphological characteristics (Fig. 1B). Morphological assessment confirmed that UK duckweed consisted of species in the *Lemna* and *Spirodela* genera.

**Figure 1.**
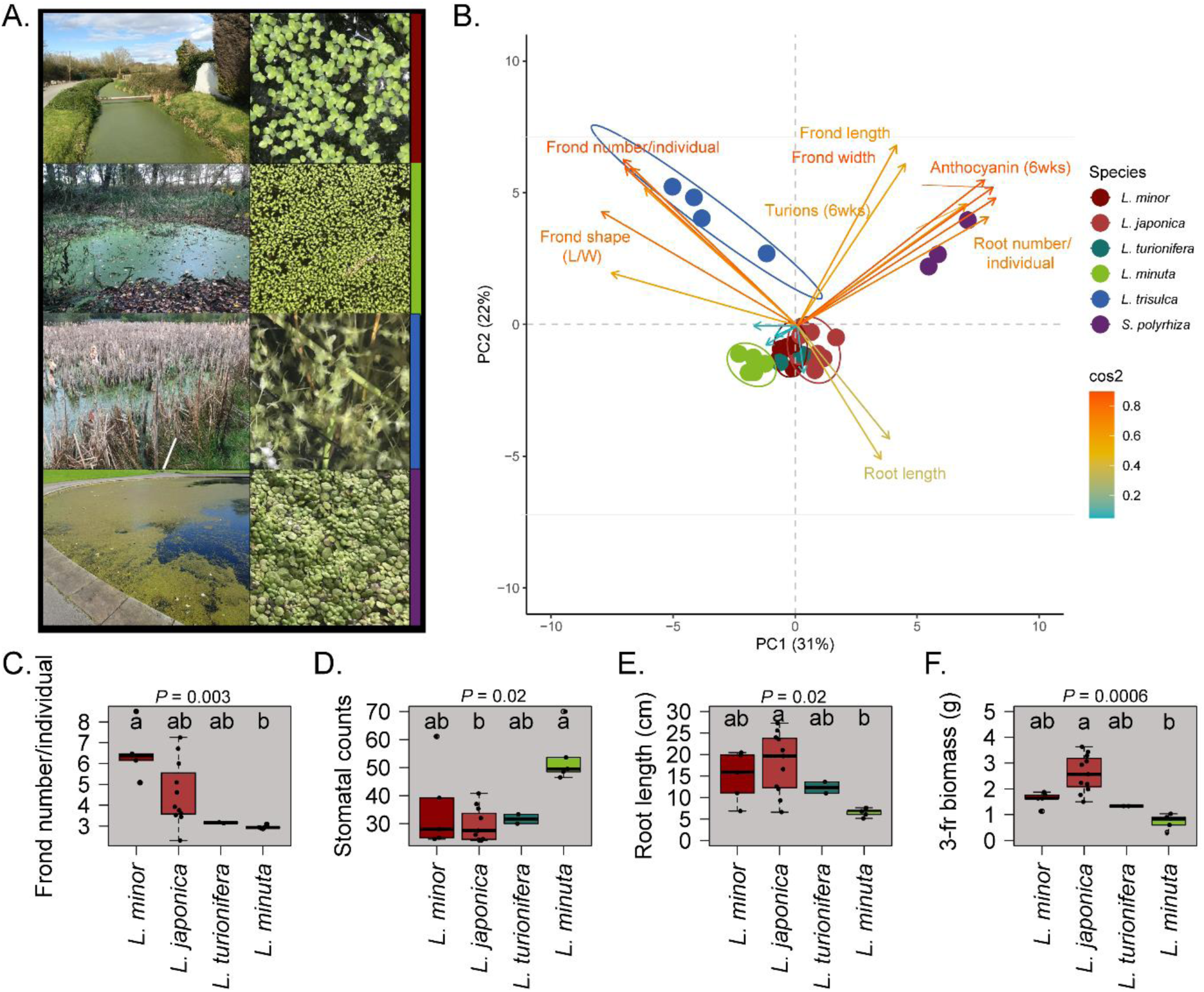
Twenty-two phenotypic traits used to classify five UK duckweed species. *A.* Photographs of four duckweed species growing in native sites with colour codes: genus *Lemna* (*L*.) *L. minor* (red), *L. minuta* (green), and *L. trisulca* (blue) and *Spirodela* (*S*.) *S. polyrhiza* (purple). *B.* A subset of the UK cohort consisting of 30 accessions used for species confirmation by phenotyping. Morphological traits are represented on a PCA using principal components PC1 and PC2 to explain ∼50% data variation. Ellipses display 90% confidence intervals for species groups and overlaps indicate reduced morphological criteria to differentiate between four *Lemna* species: *L. minor*, *L. minuta*, *L. turionifera* and *L. japonica*. Within species groups the number of accessions were: *L. japonica* = 11, *L. minor =* 5, *L. minuta =* 5, *L. trisulca* = 4, *L. turionifera* = 2 and *S. polyrhiza* = 3. Arrows are coloured by squared cosine (Cos2) with values > 0.5 showing phenotypic traits contributing most to dataset variation on PC1 and PC2. *C-F.* Differences in morphological traits between four *Lemna* species *C*. Frond number per individual *D*. Root length *E*. Stomatal counts and *F*. Three frond biomass per individual (3-fr biomass). Boxes display median and 25% and 75% percentiles for each species. Kruskal-Wallis *P =* < 0.05 was used to derive phenotypes significantly different between species and are indicated on the top of each plot. Different letters within the box plots indicate significant differences between species using a Dunn’s post-hoc test with Bonferroni adjustment using *P =* < 0.05. For *L. minor* and *L. minuta* difference in frond number (*P* = 0.001), and for *L. japonica and L. minuta* difference in root length (*P* = 0.004), stomatal counts (*P* = 0.003) and biomass of a three-frond individual (*P* = 0.0002).

*Spirodela* and *Lemna* species can be differentiated by frond and root characteristics (Fig. 1). Criteria for membership in the *Spirodela* genus included larger fronds and multiple roots per individual (Landolt, 1986), anthocyanin accumulation, shorter roots and lower length-to-width frond ratios (L:W) than *Lemna* (Fig. 1A, B). Within *Lemna*, *L. trisulca* had thin, pointed fronds, giving the highest L:W ratios and higher fronds per individual connected by long stipes (Fig. 1A, B). *Lemna turionifera* were deduced from other *Lemna* species by observations of turions (overwintering bodies) produced in nutrient-depleted conditions. *Lemna minuta* produced fewer fronds (three per individual), compared to other species including *L. minor,* producing a maximum of eight (Fig. 1C). Roots were shorter in *L. minuta* but frond adaxial stomatal counts were almost two-fold higher than other *Lemna* species (Fig. 1D:E). In contrast, *Lemna japonica* and *L. minor* could not be differentiated by morphological criteria (Fig. 1B, 1C:E).

### Genomics-based species identification

To extend the criteria for distinction of these two *Lemna* species and further confirm other species definitions, a genetic structure analysis was carried out. Individual whole genome sequencing of 122 new UK accessions was performed and mapped to a common *L. minor* 7210 reference genome. These were processed with ten additional newly sequenced from the Rutgers duckweed collection and four individuals downloaded from public repositories of known clones (see Methods, Table S2A:B) for a total of 136 accessions. All individuals were classified into clusters by a variety of genomic clustering methods, including PCA and FastStructure analysis.

This PCA shows genetic groupings by species, when using the primary and secondary principal components (Fig. 2). PC1 vastly explained 72% of the variance and clearly discriminated species groups *L. minuta* and *L. minor,* with *Spirodela* emerging on PC2. Overall UK species were clustered with known species and genomic analysis aided species discrimination, compared to just using morphology alone. Native *Lemna trisulca* species formed a cluster with (*L. trisulca* 7192), invasive *L. turionifera* (*L. turionifera* 6002) and *Spirodela* (*S. intermedia* 9394) (Fig. 2A). Genome analysis was key for distinction of native *Lemna minor* and hybrid *L. japonica* which showed similar phenotypic traits. Two clusters of *L. minor (Lmo)* clones were initially observed and differentiated as C1 and C2 (Fig. 2A, C, D). These were very closely neighbouring in the PCA but were much better discriminated by FastStructure and tree-based approaches (Fig. 2A-B:D). C1 neighboured with English *Lmo*7016 and Irish *Lmo*5500, which were inferred as *L. minor*, while C2 is located between *L. minor* (C1) and *L. turionifera* clusters on the PCA (Fig. 2A). In cluster C2, very strongly admixed European *L. japonica* 9250, Canadian *L. japonica* 7123, South African *Lmo*8389 and North African *Lmo*7295 were found along with hybrid *L. japonica* (*Ljp*) species.

**Figure 2.**
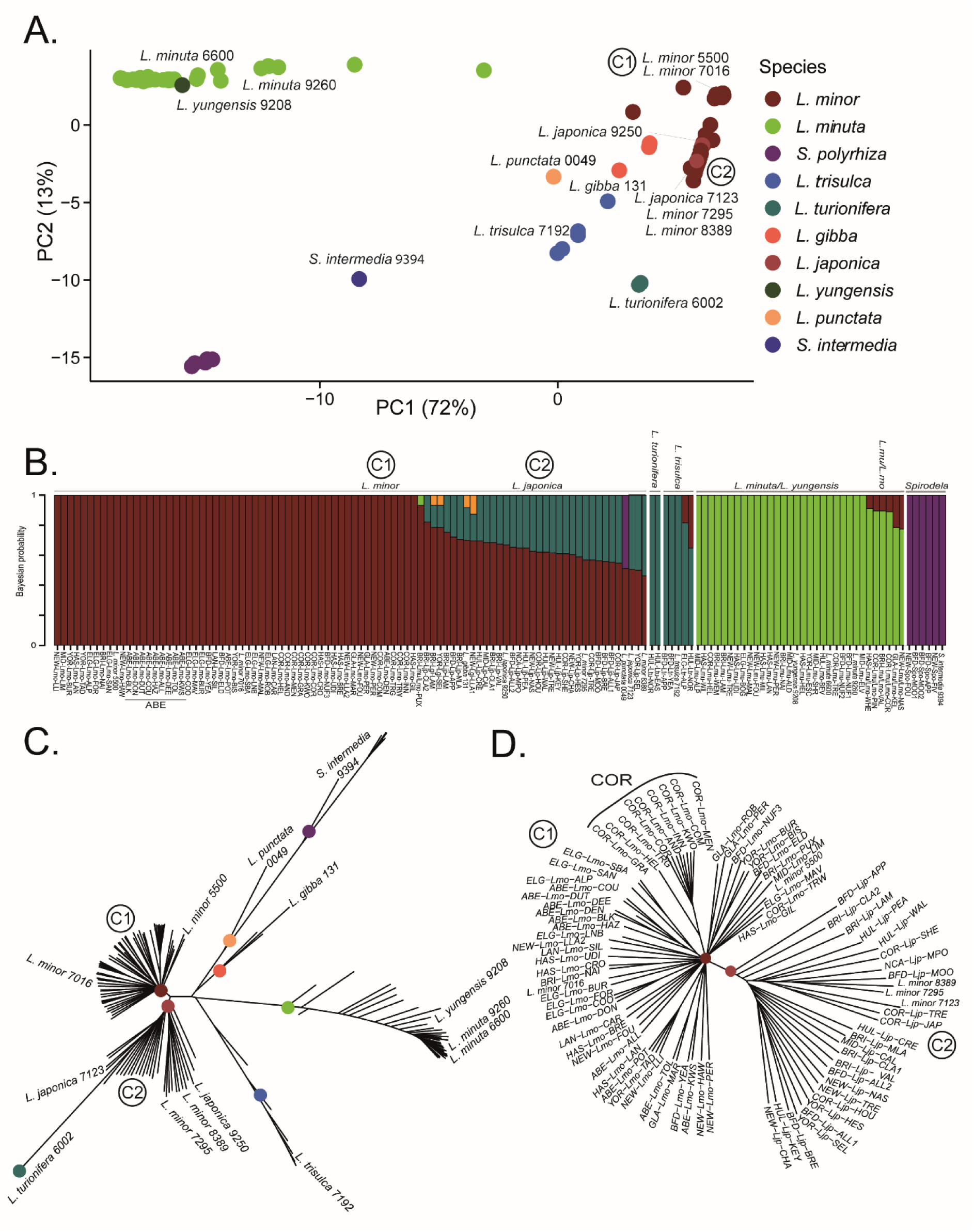
Genetic structure of 136 duckweed accessions, primarily from the novel UK cohort. One hundred and twenty-two novel UK accessions, four previously published (*L. minor* 7016, *L. minor* 5500, *L. gibba* 131, *L. turionifera* 6002), and ten newly sequenced duckweeds from the Rutgers duckweed collection. *A.* PCA of 11,088 quality-filtered four-fold degenerate (neutrally evolving) SNPs. Species are coloured by clusters determined by previously identified clones. The *L. minor* (red) clade shows two clusters conforming to membership with established, previously phenotyped clones. The *L. minor* (red) clade shows two clusters labelled as cluster one (C1) and two (C2). *B.* FastStructure analysis differentiates accessions by group membership. *Lemna minor* accessions from the ABE region are labelled. Individuals with Bayesian probability assigning them to two or more species groups show admixture and are determined as hybrid species. K=9 was the model complexity with maximum marginal likelihood. The scale represents Bayesian probability of likelihood of species membership. *C.* Neighbour-joining tree showing genetic differentiation between species. *Lemna minor* (C1) clustered with *L. minor* 7016, 5500 and *L. japonica* (C2) grouped with *L. minor* 7295 and 8389 and *L. japonica* 7123. *D.* Close-up of a neighbour-joining tree distinguishing *L. minor* from *L. japonica*. *Lemna minor* accessions from COR with a common ancestor are labelled. Clone sequences from the Sequence Read Archive (SRA) or newly sequenced in this study from the duckweed stock database are labelled in italics with their corresponding identifying number.

Structure analysis was used to estimate ancestry and to assign membership of each accession to species (Fig. 2B). This confirmed that C1 group were entirely *L. minor* species, and the C2 cluster contained accessions with substantial admixture with invasive *L. turionifera* gene pools. In the C2 cluster, the admixture between *L. minor* and *L. turionifera* species (Fig. 2B) and the intermediate cluster on the PCA is likely composed of an *L. minor* and *L. turionifera* interspecific hybrid *L. japonica (Lmo/Ltu)*. Six accessions showed admixture between *L. minuta* and *L. minor*, (*Lmu/Lmo* hybrid; Fig. 2B). Morphological criteria did not differentiate these from *L. minuta* but they likely form a presently undescribed hybrid duckweed species between native *L. minor* and invasive *L. minuta*.

Interestingly, some *Lemna* species are difficult to resolve by genetic structure alone*. Lemna yungensis* clone 9208 from Bolivia clustered with UK *L. minuta* accessions, showing a high degree of similarity between these species, both a part of the ‘Uninerves’ section of *Lemna*, consisting of one frond nerve (Fig. 2A,C, Bog *et al*; 2020). *Lemna turionifera (Ltu)* and *L. trisulca (Ltr)* are robustly separated using morphology (Fig. 1) but were undifferentiated by FastStructure (Fig. 2B), possibly due to having few representative accessions in each group (2 and 5) or low mapping efficiency to the *L. minor* reference (Table S2C).

### Highly variable species distributions by region

*Lemna minor* (*n* = 81) were the most common species in number and diversity across the UK survey (Fig. 3A). This species was found both in monocultures and co-existing with other species. *Lemna minuta* were also frequent (*n* = 30), and exhibited a marked latitudinal contrasting distribution (Fig. 3D). *Lemna minuta* prevalence in south England and Wales was greatest (20/41 sites; 49% prevalence), with presence at 11/32 sites in central England (34% prevalence) compared to negligible presence in Scotland (3/36 sites; 8% prevalence; Fig. 3D). *Lemna minor* was the only species found across all sampling sites within the ABE region in the north of Scotland, (Figs. 2B, 3). In contrast, the southwestern BRI and NEW regions had the greatest species diversity both between sites (Fig. 3) and within sites, with up to three species co-existing in several sites (Table S2A, Fig. S2I, J and S3I, J), including the less frequent *S. polyrhiza* and *L. gibba*.

**Figure 3.**
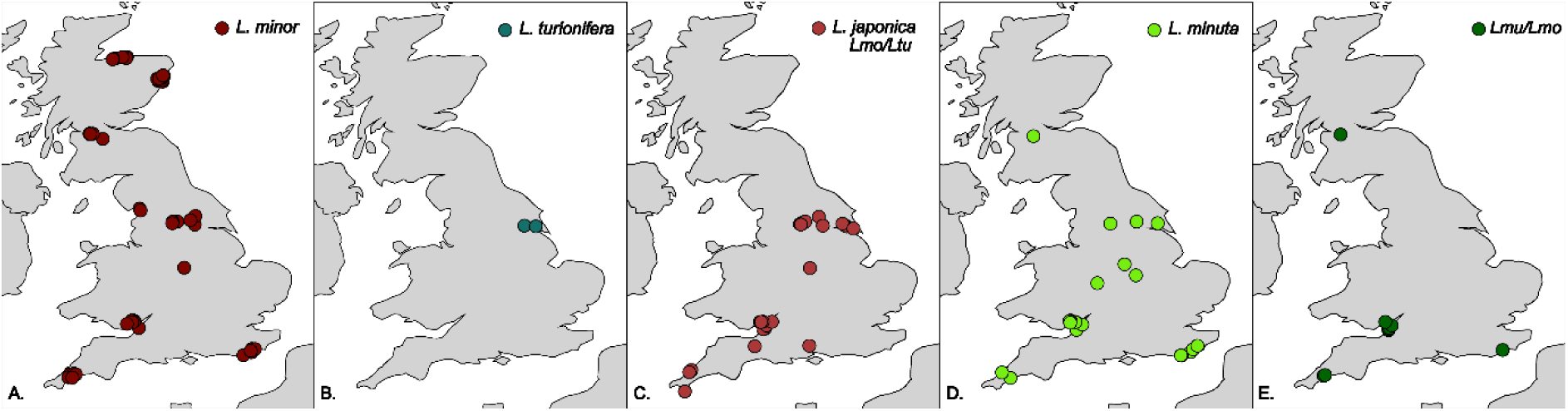
Different prevalence of *Lemna* species within UK sampling regions. Five *Lemna* species, coloured by species. *A. L. minor, B. L. turionifera, C. L. japonica. D. L. minuta. E. Lmu/Lmo*. The five species include common native duckweed (*L. minor*), invasive species (*L. minuta*, *L. turionifera*) and two hybrid species (*L. japonica (Lmo/Ltu)* and *Lmu/Lmo*). The total regions *n* = 12, and sample sites within regions *n* = <10.

*Lemna turonifera* was sparse throughout the UK; searches yielded only two accessions isolated in the northeast of England (Fig. 3B). The *L. japonica* (*Lmo/Ltu)* hybrids were abundant and overlapped with the two *L. turionifera* accessions, from which interspecies hybridisation may have occurred (Fig. 3B, C). *Lemna japonica* were more prevalent than the *L. turionifera* parental species but not as cosmopolitan as the *L. minor* parental species, as they were not found in Scotland (Fig. 3A, C). Conversely, *Lmu/Lmo* hybrids were found in southern regions, mirroring the pattern of parental species *L. minuta* (Fig. 3D, E). Single accessions of *Lemna minuta*, *Lemna japonica* and *Lmu/Lmo* were found in GLA. There were no hybrid or invasive species found in regions in the north of Scotland ABE and ELG during this survey (Fig. 3C, E).

Duckweed species broadly classified as native, invasive and hybrid types following morphological and genomic assessments. Native UK species included *L. minor*, *L. trisulca* and *S. polyrhiza*. We aimed to characterise presence *of L. minuta* invasive species, but we found that and an additional invasive species, *L. turionifera*. Furthermore, two hybrids formed between native and invasive species *L. japonica* and *Lmu/Lmo* were detected. Species types exhibited different regional distributions (Fig. 3) and further showed contrasting whole-plant ionomes in common, replete conditions, along with native water elemental differences between derived habitats (Fig. 4).

**Figure 4.**
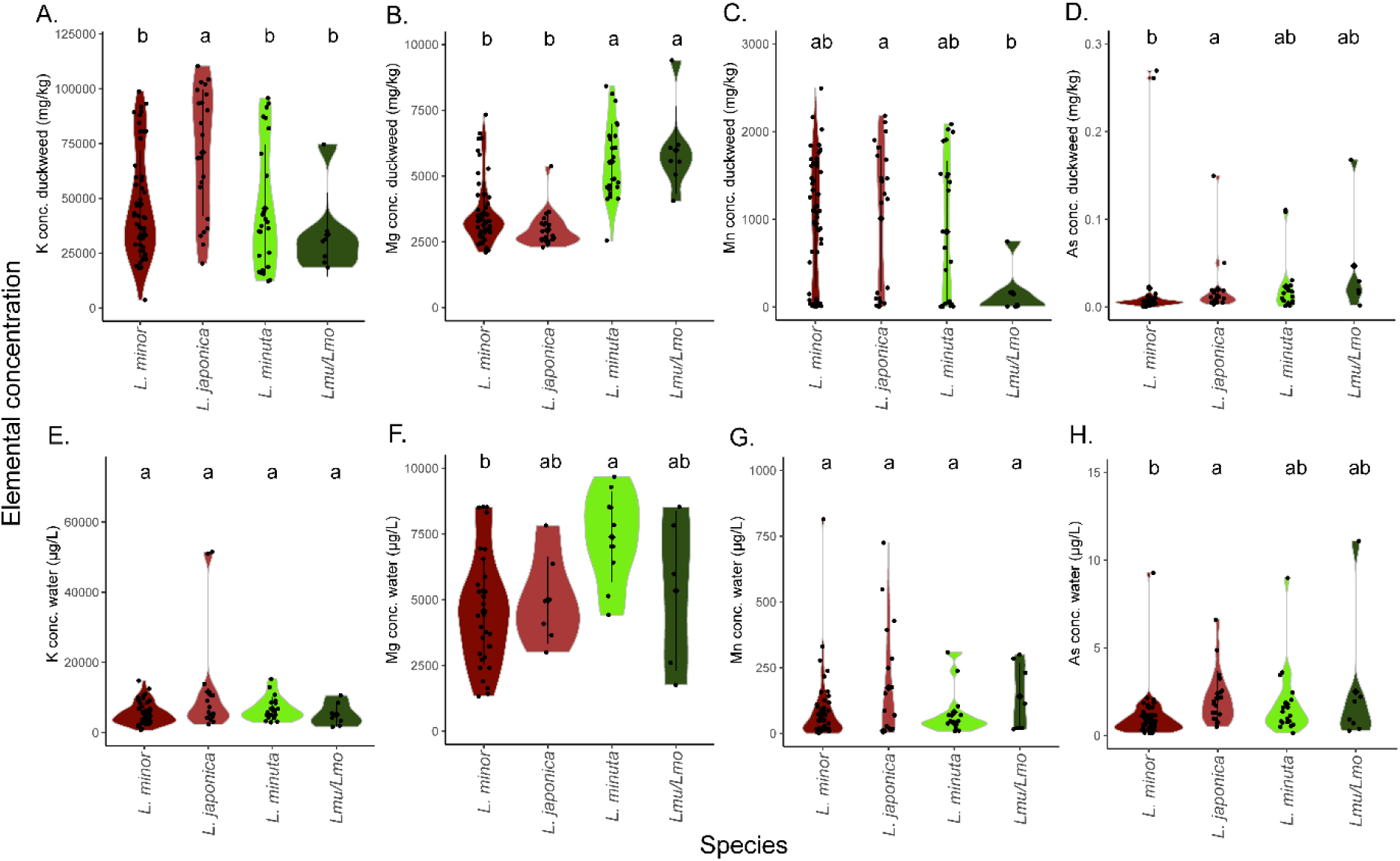
Macronutrients (K, Mg, Mn) and As composition varies between species, with additional variation of Mg and As between water environments. *(A-D)*. Elemental composition of K, Mg, Mn and As, for whole duckweed tissue. *(E-H)*. Environmental water concentrations of K, Mg, Mn and As, grouped by species. Whole tissue ionome element content averaged per individual (mg/kg). Site water average elemental composition in µg/L from *n* = 3 water replicates per site. Significance was assessed by a Kruskal-Wallis test and Dunn’s post-hoc test with Bonferroni adjustment using *P* = < 0.05 to indicate species significant differences using letters above plots.

### Variable ionomic profiles are species-specific

To infer relationships of genetic local adaptation, native water chemistry was compared with ionomes of plants grown in common nutrient replete conditions. In total, twenty-six elements were measured in 116 accessions from 100 water sampling sites. After classification into species using phenotyping and genomic clustering, species differences between plant ionomes detected in a common garden and their home water chemistries were compared (Fig. 4). Overall, tissue levels of Mg, K and Mn contents varied significantly between species grown in common conditions (Fig. 4A, C, D). Interestingly, the two hybrid species *L. japonica* and *Lmu/Lmo* showed both higher and lower levels of Mg, K and Mn in both upper and lower directions, which may point to transgressive segregation (Fig. 4A-D), which can provide an evolutionary advantage for their presence in stressful environments. The highest Mg content overall was found in *L. minuta*, followed by hybrid *Lmu/Lmo* (*P* = < 0.0001, Fig. 4A, Table S3). Hybrid *L. japonica* had higher K levels than other *Lemna* species (*P* = 0.0015, Fig. 4C, Table S3) and hybrid *Lmu/Lmo* also had reduced Mn compared to other species, whilst *L. japonica* had the most (*P* = 0.044, Fig. 4D, Table S3). In some cases, hybrids mirrored one of their parental phenotypes, as found for internal Mg content in *L. minuta* and *Lmu/Lmo* hybrids (Fig. 4B).

Species showed contrasting accumulation profiles in replete nutrient conditions and also showed differing originating water elemental profiles. *Lemna minuta* accumulated more Mg and was found in higher Mg environments (Fig. 4B, F, Tables S3, S4). *Lmu/Lmo* accumulated comparable higher accumulation of Mg mirroring its parent, but the environmental Mg was not above average. *Lemna japonica* showed higher As accumulation and higher As contamination in water environments (Fig. 4D, H, Tables S3, S4), demonstrating one example of higher elemental compositions in both native water and in ionomes of originating species. Additionally, *L. japonica* was found on waters higher in other macro- and trace minerals Ca, B, Mo, Sr than *L. minor* (*P* < 0.05, Table S4). *Lemna japonica* accumulated higher K levels than other *Lemna* (Fig. 4A). However, originating water levels of K did not vary between species (Fig. 4E, Table S4). Similarly, there was a disconnect between higher Mn accumulation in *L. japonica* and reduced Mn in *Lmu/Lmo* respectively but no difference in species originating water levels of Mn (Fig. 4C, G, Tables S3, S4).

### Widespread within-species ionomic variation in common conditions

In common conditions all accessions accumulated macronutrients P, K, Mg, Ca above the hyperaccumulation threshold of 1 g/kg. Overall, the largest variation of tissue concentrations between duckweed accessions were found for Mn and Pb, followed by S (Table S5). Almost all accessions hyperaccumulated S, with the exception of four Scottish accessions (ELG-*Lmo*-BUR, GLA-*Lmo*-PER, GLA-*Lmo*-CHA, GLA-*Lmu/Lmo*-KEL) and two from NEW (NEW-*Lmo*-LLI, NEW-*Lmo*-CHA), mostly *L. minor* species. The most variable element in the duckweed ionome was Mn, with 59/116 accessions hyperaccumulating it. In contrast, accumulation of Na was relatively rare with 17/116 accessions hyperaccumulating, but others maintaining very low levels. For other independent elements, rare, single accessions accumulated those: for example, B was hyperaccumulated by accession HAW, Si by accession BOG and Fe by accession LAN (Table 1). All other trace elements and metals were below the hyperaccumulation threshold, possibly due to limited presence of these in N-medium.

**Table 1.** Accessions showing accumulation of elements as measured by ICP-MS. Green triangles 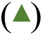 indicate higher accumulation and red triangles 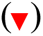 indicate reduced accumulation compared to cohort average. Accumulation differences are considered significant when z-scores exceed +/- 2 SD for *n* = 116 accessions. *Lmo* - *L. minor*, *Lmu* - *L. minuta*, *Ljp* - *L. japonica*.

| Accession | Region | Species | Duckweed ionome elemental variation |
| --- | --- | --- | --- |
| BOG | GLA | *Lmu* | ▲ B, Na, Si, S, Ba, Pb, Al, Ti, Zn, Cd ▼ P |
| LAN | HAS | *Lmu* | ▲ Fe, S, Si, Mg, Pb, Al, Ti, Zn |
| HAW | NEW | *Lmo* | ▲ B, Ca, Fe, Cd, Ba, Pb ▼ K, Rb |
| ALL2 | BFD | *Lmo/Ltu hybrid (Ljp)* | ▲ B, Na, Sr, Ba, P ▼ Fe |
| MAV | ELG | *Lmo* | ▲ B, Fe, Cu, Ba, Cd |
| APP | BFD | *Lmo/Ltu hybrid (Ljp)* | ▲ Mo, Na, P |
| CRO | HAS | *Lmo* | ▲ B, Ca, Cu |

Often hyperaccumulation of one element is accompanied by changes in suites of others (Table 1). The hyperaccumulating accessions (HAW, BOG, LAN) all had higher concentrations of multiple other elements, including heavy metals, compared to the cohort average (Table 1, Fig. S2C,J,L S3C,J,L), indicating some interdependence. BOG showed the most differential ionome, accumulating ten different elements. Overall, B was the most accumulated element in five out of seven higher accumulator accessions. Higher accumulation of elements co-occurred with reduced levels of macronutrients P, K and Fe in some instances (Table 1). High accumulators consist of several *Lemna* species and originated from a range of collection regions (Table 1, Fig. S2 and S3).

### Regional and local-scale site water elemental variation

Simultaneous with duckweed collection, water samples were collected for elemental composition in order to relate environmental chemistry with those ionomes of specific accessions. High nutrient water bodies included the BRI southwestern region, which exhibited higher concentrations of S, Mg, Ca and alkali metals Li and Sr and from the BFD region, higher concentrations of K, B and Mo (Table 2, Fig. S6). Different regions showed different water hardness, with ranges of over 2,000-fold in Ca, 56-fold in Mg and nearly 7,000-fold differences in Mn concentrations (Table S6). Additionally, within 19 sites in the BFD, YOR and HUL regions, the levels of Mg, Ca, Mn and Fe were highly variable due to the effect of seasonality, with the site ALL showing the largest variation overall (Fig. S7, Table S7),

**Table 2.**
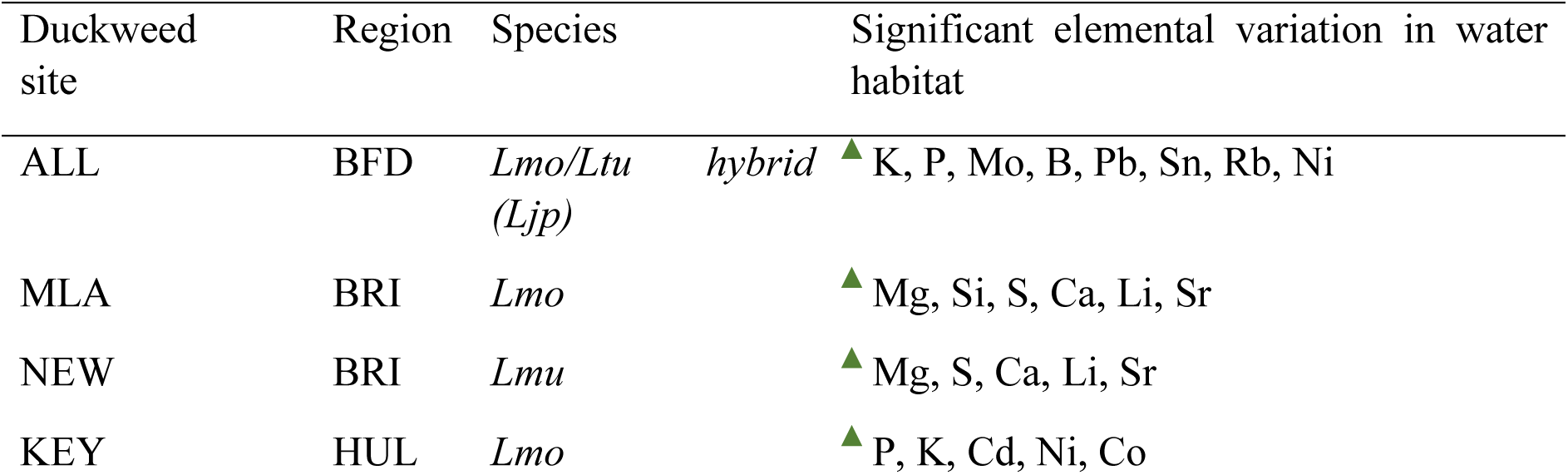
Native duckweed environmental water sites with significant levels of five or more elements as measured by ICP-MS. The table summarises water sampling sites with increased levels of macronutrients and heavy metals. Green triangles 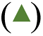 indicate higher accumulation compared to water site average and is considered significant when z-scores exceed +/- 2 SD. Species refers to the duckweed species found at water sites, *Lmo* - *L. minor*, *Lmu* - *L. minuta*, *Ljp* - *L. japonica*.

| Duckweed site | Region | Species | Significant elemental variation in water habitat |
| --- | --- | --- | --- |
| ALL | BFD | *Lmo/Ltu* *hybrid* (*Ljp*) | ▲ K, P, Mo, B, Pb, Sn, Rb, Ni |
| MLA | BRI | *Lmo* | ▲ Mg, Si, S, Ca, Li, Sr |
| NEW | BRI | *Lmu* | ▲ Mg, S, Ca, Li, Sr |
| KEY | HUL | *Lmo* | ▲ P, K, Cd, Ni, Co |

### Duckweed ionome responses in relation to native aquatic environments

To infer whether accessions exhibited signal of specific local genetic adaptation, common garden ionomes of each accession were compared with the corresponding native water chemistry using linear models with Pearson correlation. At the grossest scale, no significant relationships were observed for twenty-two elements between water and duckweed ionomes across the sampling range as a whole. However, at a finer scale, associations by region were evident (Fig. 5) showing that region- or concentration-specific levels of elements in water habitats may therefore drive specific duckweed responses.

**Figure 5.**
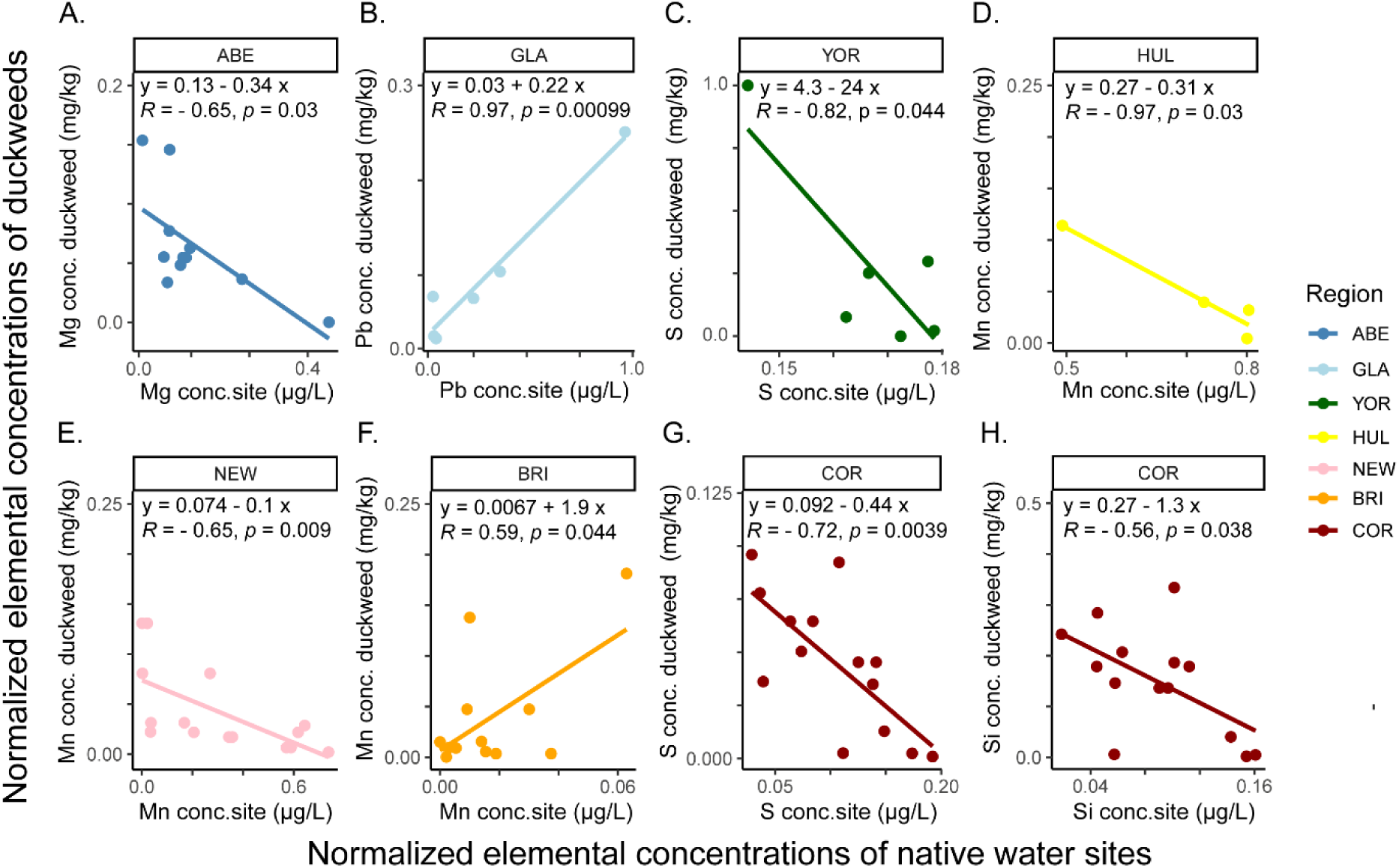
Specific regions show significant directional linear associations between site water elemental concentrations and *Lemna* duckweed tissue concentrations. *A-D.* Elements with significant correlations in northern UK regions. *E-H.* Elements with significant correlations in southern UK regions. Normalized water elemental concentration (µg/L) are recorded on the x axis with normalized duckweed whole tissue element concentration on the y axis (mg/kg) to scale values between 0 – 1. Duckweed accessions and sites within each region is *n*=>5. y and x represents line slope and intersect, *R* and *P* values depict Pearson model coefficients. *R* values <0.50 or <-0.50 are significant when *P* = <0.05. Non-*Lemna* (*Spirodela*) individuals were removed from analysis.

Together with its variability in water sites, Mn shows varied tissue-level responses in duckweeds from different regions (Fig. 5D:F). In BRI, concentrations of Mn in duckweed tissue shows an increasing trend with increasing Mn concentrations present in native sites. In contrast in NEW and HUL, duckweeds from sites associated with higher Mn concentrations show reduced Mn tissue contents in replete conditions. Higher accumulation of Pb by GLA accessions was positively associated with higher water levels of Pb in native environments (Fig. 5B). Concentrations of macro- and micro-nutrients including Mg, S and Si showed the opposite trend in specific regions, with higher concentrations in duckweed tissue associating with lower concentrations of these elements in water sites (Fig. 5A,C,G,H). Therefore, in these specific regional cases, heavy metals such as Pb tended to increase in duckweeds originating from contaminated water environments, but for some macro- and micro-nutrients, low levels in the water environment potentially stimulated higher relative accumulation in duckweeds from these native sites when grown in replete nutrient conditions.

Water body elemental levels did not appear to drive broadly deviant plant ionomes. Whereby, none of the accessions with particularly many elements outside the normal range had higher or lower elemental concentrations compared to the UK average in their source water profiles (Tables 1, 2, Fig. S2, S5). From the seven accumulating accessions presented in Table 1, (including hyperaccumulators of B, Si and Fe), six of these came from water bodies with broadly normal levels of elements, including B, Si and Fe (Fig. S5C). An excellent example is the accession BOG which exhibited the most extreme ionome overall, hyperaccumulating ten macronutrients and heavy metals with reduced P (Figs. S2C, S3C). However, the source water chemistry from which BOG was collected harboured no elements significantly differing from the cohort average (Figs. S4C, S5C). Concurrently, accession ALL1 from the most nutrient-dense and contaminated ALL site (Figs. S4D, S5D) only highly accumulated Fe in replete conditions (Tables 1, 3, Figs. S2E, S3E). It is noteworthy that Fe, K and B were highly variable at this site, with Fe showing maximum levels in autumn 2020 but in decline until 2022 (Fig. S7).

From this survey of duckweed in the UK, native *L. minor* was diverse and commonly found. Presence of invasive species *L. minuta* and *L. turionifera* were more limited as were new reports of hybrid species *L. japonica* and *Lmu/Lmo*. Species showed different potentials to uptake macronutrients (Mg, K, Mn) and heavy metals (As) and also inhabited waters with diverse elemental profiles. Hyperaccumulators and high accumulator accessions were detected, usually coincident with higher uptake of multiple elements. Relationships between duckweed ionomes and water profiles were complex, with specificity to the element in consideration with accessions, species and in a few localised regions.

## Discussion

Region-wide genomic assessments of duckweed diversity are scarce. Furthermore, existing duckweed collections lack data on source environmental parameters, and no study has assessed local-scale whole-plant ionomes in common conditions using regional sampling. Nor is there any genomic assessment of invasive duckweed impact on native accessions. In this study we fill these gaps, and further interpret this information with a view toward identification of useful accessions tailored to phytoremediation and food development applications.

### UK duckweeds are composed of native, invasive and hybrid *Lemna* species

This survey confirms that native *L. minor* is cosmopolitan and thus still more prevalent than invasive *L. minuta*, which had a more limited distribution, especially in the north. *Lemna minor* is evidently well-adapted to the UK environments here, although hybridisation with invasives is a liability. *Lemna minuta* was only prevalent in the south and midlands of England with some ingress into Scotland. *Lemna minuta* has been added to the Global Register of Introduced and invasive Species of Great Britain (GRIS); however observations have been in decline since 2019, according to the Global Biodiversity Information Facility (GBIF, 2022). Thus, presence of *L. minuta* is possibly not as damaging to native species as previously reported (Paolacci et al., 2018b) and this study found evidence of co-occurrence of both species in water bodies. Although invasive species can be opportunistic in *in-vitro* high light and Mg conditions (Paolacci et al., 2016; Paolacci et al., 2018a), dominance of invasive or non-invasive species depends on competition in particular environments (Paolacci *et al*., 2016, 2018; Ceschin *et al*., 2018b; Gérard & Triest, 2018). It appears that environments in Scotland are less suited to promote opportunism in *L. minuta* species.

Within the worldwide duckweed collection (such as the Landolt collection), many *L. minor* species have been reclassified as hybrids including *L. japonica* and *L. mediterranea* (Braglia et al., 2021; Braglia et al., 2024), indicating that the previous assignments of *L. minor*, *L. japonica* and other *Lemna* species may not be fully correct. Genetic contribution from both *L. minor* and *L. turionifera* in *L. japonica* accessions indicated admixture and therefore hybrid presence in the UK. The presence of both *L. japonica* and *Lmu/Lmo* hybrid varieties have not been previously reported in the UK. In part, because the morphology of hybrid and native species can be very similar making it difficult to differentiate between them without genomic or ionomic confirmation, as performed here.

Indeed, *L. japonica* was not easily differentiated from *L. minor* by morphology here (Fig. 1) and in Eastern Europe (Volkova et al., 2023). Among these hybrids, there is heterogeneity of parental introgression, putatively resulting in fitness performance attributable to parental gene variants in the context of different ecological backgrounds (Gompert & Buerkle, 2012). Similarly, *L. japonica* accessions exhibit variable propensities to form turions under inductive conditions, probably in line with receiving varied allelic contributions from *L. minor* (non-turionating) or *L. turionifera* (turionating) parents (Ernst *et al*, 2023).

Hybrid *L. japonica* appears more successful than its invasive parent *L. turionifera*, which showed limited geographic presence in this UK survey (Fig. 4B) and has only been reported twice previously, also localised in the east England region (Lansdown, 2008). In this instance, hybridisation has afforded wider spread than one of the parental genotypes (Volkova et al., 2023), especially in the case of *L. japonica*. Hybridisation thus may occur to access wider adaptative potential of genetic variants from native species. Supporting this, we find evidence for hybrid differences in both water elemental niches and nutrient uptake.

### Species show differences in ionomes and native water chemistry

Hybridisation is a proposed mechanism to generate ionome variation in plants and aid their adapt to the water environment (Chen et al., 2017; Wang et al., 2021). Hybridisation also increases vigour in plants by increasing allelic diversity, especially in stressful environments (Washburn and Birchler, 2014). *Lemna japonica* reportedly form both diploid and triploids, and contain more transposable elements compared to parental species, hinting at possible increased capacity for environmental adaptation (Hoang *et al*., 2022, Ernst *et al*., 2023). There is also experimental support for increased adaptation to high light irradiance in *L. japonica,* relative to parental species (Smith et al., 2024a). Hybrids and parental species showed some differences in elemental composition between their water environments too, in line with possible adaptive speciation to specialised environmental niches.

Here, *L. japonica* water sites were contaminated with more As than *L. minor* sites and included higher concentrations of Ca, Mo, B and Sr. These elements were typical of BRI-region water bodies linking species distribution with adaptation to a differential water environmental niche in the southwest. Transgressive phenotypes can arise commonly in interspecific hybrid plants to give them higher abiotic tolerance during niche establishment, to enable divergence away from competition from parental species (Rieseberg et al., 1999). Transgressive phenotypes are common in hybrids, for instance to improve NaCl and Cd tolerance from that of parental species (Xue et al., 2021; Ortega-Albero et al., 2023).

In this study, invasive *L. minuta* was found in higher Mg-containing sites than native *L. minor*, accumulating more Mg than other *Lemna* species. This finding provides affirmation that this species is a high Mg-tolerator, supporting both invasive behaviour in foreign environments, possible tolerance to hardwater areas and enhanced potential for phytoremediation of high Mg-containing wastewater (Paolacci et al., 2016; Ceschin et al., 2020; Walsh et al., 2020). *Lemna minuta* and hybrid *Lmu/Lmo* had the highest internal Mg concentrations overall, showing a mirroring phenotype between parent and hybrid species. Higher Mg in *L. minuta* is consistent with enhanced Mg accumulation occurring within species within the ‘Uninerves’ section of *Lemna* (Smith et al., 2024b) but for the first time this work links accumulation to higher Mg tolerance in native habitats.

From the UK panel, *L. japonica* shows higher K and Na tissue concentration compared to parent *L. minor*, which is consistent with findings from a worldwide duckweed ionome comparison (Smith et al., 2024b). As K is widely available, provided at the highest concentrations in water sites found here and all accessions accumulated it, it can be inferred that increased accumulation of K has a functional purpose in *L. japonica*, and may enhance tolerance to other elements, such as As to allow niche environment establishment. Some support comes from *Arabidopsis thaliana* and *Vicia faba* (Broad bean), whereby higher K uptake mitigated As and NaCl toxicity (Chao et al., 2013; Che et al., 2022; Shah et al., 2022).

### Duckweed accumulator accessions and their potential applications

Using native water data, duckweed tolerance to macronutrients in the environment could be defined at a regional-scale, further highlighting specific UK accessions with tolerance traits for phytoremediation. Here, UK native water sites showed 81%, 74% and 44% higher maximum values for Mg, Ca and K than previously reported (Linton and Goulder, 1998). Additionally the maximum concentrations of K, Mg, Ca, Mn and Fe in native water exceeded concentrations of these elements tolerated by *L. minor* grown on dairy wastewater (O’Mahoney et al., 2022) and. Thus, inhabiting accessions may have developed useful tolerance and accumulating traits for phytoremediation and enhanced supply of macronutrients for nutrition.

That said, high accumulators did not tend to come from sites with higher elemental concentrations (accession BOG), nor did high accumulators necessarily come from contaminated UK environments (accession ALL1). Therefore, there is some degree of unlinking of elemental tolerance from accumulation potential. The accession GLA-*Lmu*-BOG showed enhanced concentrations of Ba, Pb, Al, Ti, Zn and Cd (Fig. 3D, Table 1) and can be considered for remediation of contaminated water courses, provided the tendency to accumulate can be tested on real-world water conditions. For example, Zn and Pb pollution in GLA waterbodies require remediation and the accession is already inhabiting the region (Fordyce et al., 2019; Eschenfelder et al., 2023). Unfortunately, accumulation of several macronutrients including B and Fe co-occurred with heavy metal uptake, surprisingly even in low level controlled conditions. Whilst willingness to uptake heavy metals is an optimal trait for phytoremediation, this is problematic for direct applications in nutrition. Thus, mechanisms to retain high macronutrients but mitigate heavy metal levels such as inoculation with a synthetic microbiome or post-harvest washing or cooking steps may be future directed targets for duckweed consumption (Stout et al., 2010; Sattar et al., 2015; Amir et al., 2019).

In conclusion, this collection is the first of its nature, presenting a unique mixture of large-scale duckweed genomics, ionomics, and species assessment with environmental water data. Hybrid and invasive species were associated with novel water chemistry niches compared to parental species but were more restricted in their ranges. Therefore, high water hardness may be a predictor for future *L. minuta* colonisation in new regions as well as highlighting their potential in bioremediation. It is possible that invasive species are not currently establishing well in north Scotland but their introduction should be mitigated. Heavy metal accumulating accessions identified from this study (BOG, LAN and others) should be further explored for phytoremediation potential using outside transplantation experiments and *in-vitro* elemental spiking experiments to maximize hyperaccumulation. This UK collection serves as a useful resource to explore desired traits for human consumption, bioremediation or ecological population studies and promotes the further genetic understanding of hybrid and parental duckweed species.

## Methods

### Selection of site locations

An inland-coastal transect with decreasing altitude was selected locally to give seasonal time points for water collection. Original sites were chosen in May 2020, concordant with duckweed collection and further duckweed and water collections performed during autumn 2020, summer 2021, autumn 2021 and winter 2022. For spatial assessment of the UK, duckweed and water collections were conducted in spring 2021, starting at southern locations in early April and finishing mid-May 2021 in northern Scotland, to account for variation in springtime across UK. Regions were chosen to span the UK and using duckweed observations reported recently using the Global Biodiversity Information Facility (GBIF.org, 2022). Duckweed observations were particularly dense in BRI and ELG. Locations from GBIF.org were mapped onto Google maps and from that, several potential sites were searched within each region to give n=>6 local sites with duckweed presence.

### Collection of duckweeds and morphological assessment

Duckweeds were collected as described in (Smith et al., 2024a) for temporal and spatial collections. In sites with more than one suspected duckweed species, these were collected and cultured separately based on size and denoted as A, B, or C. From across 19 sites along the seasonal transect, duckweeds were phenotyped at each time point, and a handful re-sequenced and denoted as 1, 2 or 3, when they showed differences from species previously characterised there. For other sites, duckweeds were collected at a single time point. Frond characteristics (length, width, length width ratio (L:W), number of fronds per individual and anthocyanin presence) were assessed initially from images taken with a Zeiss SV6 stereo microscope (Ziess, Oberkochen, Germany) (*n* = 10 individuals per accession). Then each of these characteristics and root lengths were measured at two later timepoints after lab cultivation. All images were analysed with Fiji (Schindelin et al., 2012). Stomatal counts were performed for *n* = 3 whole fronds of 24 accessions using a Leica TCS SP5 confocal microscope (Leica, Wetzlar, Germany) using preparations as described in (Kurihara et al., 2015; Smith et al., 2024b). Biomass was measured for three frond individuals (*n* = 3 per accession) and presence or absence of turions were assessed from cultures exhausted in nutrient media over three years.

### Collection of water samples

For seasonal water collection, 100 ml samples were taken four times at each site, unless accessibility issues or water was not present at each time point. Solid Phase Microextraction Polytetrafluoroethylene (SPME PTFE) amber bottles were pre-washed with ultra-pure 10% nitric acid overnight followed by soaking in MilliQ water (Milipore, USA) and then thoroughly air dried. At each site, water bottles were washed at the top surface of the water, filled to 100 ml and 0.5% ultra-pure 1 ml nitric acid added before storage at 4 °C. Later, 18 ml water was filtered through a 1.45 µm syringe filter into 2 ml 10% Primar grade nitric acid to acidify samples and samples were stored at 4°C for elemental analysis. For UK-wide samples, water samples were collected in triplicate per site from the top water surface in Falcon™ tubes (Fisher Scientific, Loughborough, UK) and filtered through a 1.45 µm syringe filter into High density polyethylene (HDPE) Universal 25mm x 90mm 30 ml tubes (Sarstedt, Leicester, UK). HDPE tubes were pre-weighed, then 2 ml Primar grade 10% nitric acid added and then re-weighed. After addition of 18 ml water from each site into acid, tubes were stored at 4°C before ICP-MS analyses. All samples were re-weighed after water collection using a precision 5 dp balance (Mettler Toledo, Ohio, USA).

### Plant care and harvesting for DNA sequencing

Duckweeds were sterilised using 0.5% sodium hypochlorite and grown in GEN2000 SH, controlled cabinets (Conviron, Winnipeg, Canada). Among all collections, four accessions from the south and 14 accessions from Scotland could not be successfully cultured in laboratory conditions. The majority of losses occurred in *L. trisulca* and provisional *L. gibba* clones, possibly due to sensitivity to sodium hypochlorite or specific adaptation to locality so were not included for sequencing or ionomics. After sterilisation and weekly media changes of successful cultures, independent sealed flasks of UK accessions were grown for four weeks to bulk tissue for DNA harvesting. Duckweeds from the Landolt collection and available at Rutger’s stock database (www.ruduckweed.org) were also grown and harvested for DNA to provide known species controls. For each accession, 20-100 mg fresh duckweed tissue was harvested into liquid nitrogen and then stored at -80 °C.

### DNA isolation, short-read library preparation and sequencing

Accessions were ground using a Tissuelyser II (Qiagen, Hilden, Germany) and DNA extracted using DNAeasy Plant kit (Qiagen, Hilden, Germany). DNA quantification was performed using dsDNA HS assay (Thermo Fisher Scientific, Massachusetts, USA) and Qubit 2.0. DNA was diluted to < 20 ng/µl with sterile MilliQ water. Individual Illumina DNA Prep (Illumina, San Diego, USA) sequencing libraries were prepared at the Deepseq sequencing facility, University of Nottingham, UK. on a Mosquito HV (SPT Labtech, Melbourn, UK) liquid handling robot using 1/10^th^ volumes at all steps. A total of 9-48 ng of DNA was used as library input and 5 cycles of Polymerase Chain Reaction (PCR) were used for the library amplification step. Final libraries were normalised and pooled on a Fluoroskan Ascent fluorometer (Thermo Fisher, Massachusetts, USA) and the resulting pools were size selected using 0.65X Ampure XP (Beckman Coulter, California, USA) to remove library fragments < 300 bp. Short read sequencing using paired end reads with Illumina HiSeq 2500 platform sequencing was performed by Novogene, Cambridge, UK using a target of 20x coverage.

### Variant calling

The processing pipeline involved three parts: (1) preparing the raw sequencing data, (2) mapping and re-aligning the sequencing data and (3) variant discovery (GATK v.4 following GATK best practices). In addition to newly sequenced samples, previous sequencing data was downloaded from the National Centre for Biotechnology Information (NCBI) Sequence Read Archive (SRA) and are summarised in Table S2B. To prepare the raw sequencing data for mapping, the different sequencing lanes were concatenated, followed by quality trimming using Trimmomatic (Bolger et al., 2014). All genomes were then aligned to reference genome *L. minor* 7210 (SRR10958743) using BWA 0.7.17 (Li & Durbin, 2009) and processed using Samtools v1.9 (Li *et al*., 2009) and duplicate reads flagged using ‘MarkDuplicates’ from picard-tools 1.13464 followed by GATK v.4 to re-align reads around indels (McKenna et al., 2010). The variant dataset was filtered for biallelic sites and mapping quality with GATK using QD < 2.0, FS > 60.0, MQ < 40.0, MQRankSum < -12.5, ReadPosRankSum < -8.0, HaplotypeScore < 13.0 and sites remaining after depth filtering DP <141 carried forward for analysis. The code for batch processing is available at https://github.com/mattheatley/ngs_pipe.

### Genomic analysis

Degenotate (https://github.com/harvardinformatics/degenotate) was used to identify sites encoding fourfold degenerate sites (4FDS) as proxies for neutrally-evolving sites. These sites were further filtered >20% missingness to reduce the cohort from 143 individuals to 135. The dataset was pruned by linkage disequilibrium in order to obtain independent segregating markers out of linkage using a custom script (Hämälä et al., 2024). The final genomic analysis included only biallelic single nucleotide polymorphisms (SNPs) at allele frequencies >2.5%, with one SNP per 100 kb sliding windows with a step size of 50 kb and r2 of 0.1. Species allocation was confirmed using a mixture of PCA, tree, and structure-based approaches using R v3.6.3. The PCA was produced for variants using ggplot2 (Gómez-Rubio, 2017). Unrooted neighbour joining trees were compiled using ape v5.4 package (Paradis & Schliep, 2019) for *L. minor* and *L. japonica*. For structure plots, 4FDS variants with > 20% missingness were dropped, removing two individuals, and then converted into genotype call files using PLINK v1.9 (Chang *et al*., 2015) based on Hardy-Weinberg equilibrium and Fisher exact tests. Allele frequencies were used for group allocation and admixture proportions by FastStructure v1 (Raj *et al*., 2014) with K groups between 4-10. Selection of K=4 was chosen for visualisation with a Structure plot v2 using Omicsspeaks http://omicsspeaks.com/strplot2/

### Duckweed growth and harvesting for ionomics experiments

After nine months of subculturing, ionomic experiments were conducted for accessions grown in controlled environment cabinets. Two individuals of each accession were grown in 500 ml Erlenmeyer flasks containing 250 ml Nutrient medium, replenished weekly for six weeks. N-medium was used as described in (Appenroth *et al*., 1996; Appenroth & Sree, 2015) and contains KH_2_PO_4_ (0.15 mM), Ca(NO_3_)^2^ (1 mM), KNO_3_ (8 mM), MgSO_4_ (1 mM), H_3_BO_3_ (5 μM), MnCl_2_ (13 μM), Na_2_MoO_4_ (0.4 μM) and FeEDTA (25 μM). Duckweeds were grown at 25 °C day and 18 °C night, with 16 h day lengths. To harvest, duckweed were rinsed with three two minute MilliQ water washes. Three replicates were obtained for each accession from three independent flasks using 150 mg duckweed tissue per sample. Duckweed were dried in an oven at 88 °C overnight and stored in a desiccator before analysis. Weights of the dried tubes were made using a 5 dp precision balance (Mettler toledo, Ohio, USA).

### Ionomics processing using inductively coupled plasma mass spectrometry (ICP-MS)

Elemental analysis for water and duckweed samples were analysed on a NexION 2000 ICP-MS (PerkinElmer, Massachusetts, USA) in Helium collision mode. For each set of water and duckweed analyses, calibration standards were run throughout using single element standards (Inorganic 226 Ventures; Essex Scientific Laboratory Supplies Ltd, Essex, UK), to subtract against background samples. Concentrations of elements in water samples were measured in µg/L for Na, Mg, Si, S, K, Ca, Al, P, Li, B, Ti, Cr, Mn, Fe, Co, Ni, Cu, Zn, As, Rb, Sr, Mo, Cd, Pb and Sn. Ti, Cr, Sn were removed from water analysis as they were below the limit of detection (LOD). Ba was also removed as it was not measured across all sites. Duckweed samples were digested with 2 ml 63% nitric acid at 115 °C for 4 hrs (spiked with the element Indium as an internal standard) and then 0.5 ml hydrogen peroxide for a further 1.5 hrs at 115 °C, before dilution into 10 ml MilliQ water. Elements Li, B, Na, Mg, Al, Si, P, S, K, Ca, Ti, Cr, Mn, Fe, Co, Ni, Cu, Zn, As, Rb, Sr, Mo, Cd, Sn, Ba, Pb were measured in duckweed tissue by dry weight (mg/kg). Elements with low levels in duckweed Li, Cr, Sn < 1 mg/kg and Ni < 3 mg/kg were removed from further analysis as they were below the limit of detection of ICP-MS.

### Analysis of ionomics data

For water and duckweeds analyses, elements were grouped as those present in duckweed growth N-medium or trace/heavy metals (not present in N-medium) for separate analysis. Water site replicates were combined to form site averages (*n* = 3) and replicates per accession combined for accession averages (*n* = 3). Standardisation for each element by z-scores were obtained by subtracting raw data for each element from the panel mean and dividing by standard deviation (SD) to produce heat maps, radar plots and PCAs. Linear regression analysis with Pearson correlation was performed between each accession’s ionome and their originating water elemental concentration averages, to find positive and negative relationships. For comparison of species ionomes, those with fewer accessions were dropped including *L. gibba*, *L. turionifera* and *S. polyrhiza*. Differences between the remaining *Lemna* species ionome concentrations and originating water elemental profiles were compared separately with a Kruskal-Wallis test and a post-hoc Dunn’s test with Bonferroni adjustment using *P* = <0.05.

## Symbols and abbreviations

ICP-MS: Inductively coupled plasma mass spectrometry.
SPME PTFE: Solid Phase Microextraction Polytetrafluoroethylene.
HDPE: High density polyethylene.
GRIS: Global Register of Introduced and invasive Species.
LOD: Limit of detection.
NCBI: National Centre for Biotechnology Information.
SRA: Sequence read archive.
dsDNA: Double stranded deoxyribonucleic acid.
GATK: Genome analysis toolkit.
SNPs: Single nucleotide polymorphisms.
4FDS: Four-fold degenerate site.
MIS: Missingness.
MAF: Minor allele frequency.
PCA: Principal component analysis.
HAS: Hastings, UK.
COR: Cornwall, UK.
BRI: Bristol, UK.
NEW: Newport, UK.
ABE: Aberdeen, UK.
ELG: Elgin, UK.
GLA: Glasgow, UK.
LAN: Lancaster, UK.
BFD: Bradford, UK.
YOR: York, UK.
HUL: Hull, UK.
MID: Midlands, UK.
NCA: Newcastle, UK.

## Author contributions

Conceptualisation, LY, KES. Methodology, LY, KES. Software, MH. Validation, KES. Formal analyses, KES, MH. Investigation, KES, LC, PF, CM. Resources, KES, MH, CARZ, AL. Writing original draft, KES, LY. Writing, review and editing, KES, LY, AL, CARZ. Funding acquisition, LY, KES. Supervision, LY. Administration, LY, KES. All authors approved the final manuscript.

## Acknowledgements

Thanks to Tuomas Hämälä for PCA and pruning by linkage disequilibrium scripts and to Ana Da Silva for the NJ tree script. Thanks to Claire Smith and Bethany Severn for harvesting DNA from some of the accessions. Thanks to Chris Dalton for aiding with collections and to Anthony Bishopp and Laura Briers for reading and revision of the manuscript. This work was supported by a Biotechnology and Biological Sciences Research Council (BBSRC) PhD scholarship BB/M008770/1 and the University of Nottingham Future Food Beacon of Excellence for KES and LY. The authors wish to thank the European Research Council (ERC) grant no. 679056 attributed to LY.

## Data availability

Sequence data that support the findings of this study have been deposited in the Sequence Read Archive (SRA; https://www.ncbi.nlm.nih.gov/sra) with the primary accession code PRJNA1030266 (available at http://www.ncbi.nlm.nih.gov/bioproject/PRJNA1030266)

## Supplemental information

Table S1. UK duckweed collection - Sites and their descriptions.

Table S2A. UK accessions and previously characterised clones newly sequenced in this study.

Table S2B. Genomes of duckweed clones downloaded from the Short Read Archive (SRA) and included in genomic pipeline.

Table S2C. Genomes of new UK duckweed accessions and newly sequenced clones included in the genomic pipeline.

Table S3. UK duckweed species show differences between tissue concentrations of elements grown in replete N-medium.

Table S4. Duckweed species show differences between typical concentrations of elements found in water habitats.

Table S5. Range of internal duckweed concentrations of eight elements grown in standard replete nutrient conditions.

Table S6. Spatial variation of eight elements in UK water sites.

Table S7. Seasonal variation of eight elements in UK water sites.

Figure S1. Duckweed sites studied, spanning twelve regions in England, Scotland and Wales.

Figure S2. Radar plots for elements quantified in duckweed ionomes from 12 UK regions grown in controlled conditions as measured by ICP-MS.

Figure S3. Radar plots for heavy metal elements quantified in duckweed ionomes from 12 UK regions grown in controlled conditions as measured by ICP-MS.

Figure S4. Spatial variation for elements measured in water environments by ICP-MS.

Figure S5. Spatial variation for heavy metals measured in water environments in ten UK regions by ICP-MS.

Figure S6. Footprint of regional variation from 100 water environments showing elemental compositions measured by ICP-MS.

Figure S7. Spatial and seasonal spikes in elemental concentrations in native water environment as measured by ICP-MS.

## References

Amir, R. M., Randhawa, M. A., Sajid, M. W., Nadeem, M., Ahmad, A., and Wattoo, F. M. (2019). Evaluation of various soaking agents as a novel tool for heavy metal residues mitigation from spinach. Food Sci. Technol. 39:176–180.

Appenroth, K. J., Sree, K. S., Böhm, V., Hammann, S., Vetter, W., Leiterer, M., and Jahreis, G. (2017). Nutritional value of duckweeds (Lemnaceae) as human food. Food Chem. 217:266–273.

Barton, K. E. (2024). The ontogenetic dimension of plant functional ecology. Funct. Ecol. 38:98–113.

Bog, M., Sree, K. S., Fuchs, J., Hoang, P. T. N., Schubert, I., Kuever, J., Rabenstein, A., Paolacci, S., Jansen, M. A. K., and Appenroth, K. J. (2020). A taxonomic revision of *Lemna* sect. Uninerves (Lemnaceae). Taxon 69:56–66.

Bolger, A. M., Lohse, M., and Usadel, B. (2014). Trimmomatic: A flexible trimmer for Illumina sequence data. Bioinformatics 30:2114–2120.

Braglia, L., Breviario, D., Gianì, S., Gavazzi, F., de Gregori, J., and Morello, L. (2021). New insights into interspecific hybridization in *Lemna* l. Sect. lemna (lemnaceae martinov). Plants 10.

Braglia, L., Ceschin, S., Iannelli, M. A., Bog, M., Fabriani, M., Frugis, G., Gavazzi, F., Gianì, S., Mariani, F., Muzzi, M., et al. (2024). Characterization of the cryptic interspecific hybrid *Lemna×mediterranea* by an integrated approach provides new insights into duckweed diversity. J. Exp. Bot. 75:3092–3110.

Ceschin, S., Leacche, I., Pascucci, S., and Abati, S. (2016). Morphological study of *Lemna minuta* Kunth, an alien species often mistaken for the native *L. minor* L. (Araceae). Aquat. Bot. 131:51–56.

Ceschin, S., Abati, S., Traversetti, L., Spani, F., Del Grosso, F., and Scalici, M. (2019). Effects of the invasive duckweed *Lemna minuta* on aquatic animals: evidence from an indoor experiment. Plant Biosyst. 153:749–755.

Ceschin, S., Crescenzi, M., and Iannelli, M. A. (2020). Phytoremediation potential of the duckweeds *Lemna minuta* and *Lemna minor* to remove nutrients from treated waters. Environ. Sci. Pollut. Res. 13:15806–15814.

Chao, D., Dilkes, B., Luo, H., Douglas, A., Yakubova, E., Lahner, B., and Salt, D. E. (2013). Polyploids exhibit higher Potassium uptake and salinity tolerance in *Arabidopsis*. Science (80-. ). 341:658–659.

Che, Y., Fan, D., Wang, Z., Xu, N., Zhang, H., Sun, G., and Chow, W. S. (2022). Potassium mitigates salt-stress impacts on photosynthesis by alleviation of the proton diffusion potential in thylakoids. Environ. Exp. Bot. 194:104708.

Chen, Z., Taylor, A. A., Astor, S. R., Xin, J., and Terry, N. (2017). Removal of boron from wastewater: Evaluation of seven poplar clones for B accumulation and tolerance. Chemosphere 167:146–154.

Chen, G., Stepanenko, A., Lakhneko, O., Zhou, Y., Kishchenko, O., Peterson, A., Cui, D., Zhu, H., Xu, J., Morgun, B., et al. (2022). Biodiversity of duckweed (Lemnaceae) in water reservoirs of Ukraine and China assessed by chloroplast DNA barcoding. Plants 11:1–14.

Ekperusi, A. O., Sikoki, F. D., and Nwachukwu, E. O. (2019). Application of common duckweed (*Lemna minor*) in phytoremediation of chemicals in the environment: State and future perspective. Chemosphere 223:285–309.

Eschenfelder, J., Lipp, A. G., and Roberts, G. G. (2023). Quantifying excess heavy metal concentrations in drainage basins using conservative mixing models. J. Geochemical Explor. 248:107178.

Fedoniuk, T., Bog, M., Orlov, O., and Appenroth, K. J. (2022). *Lemna aequinoctialis* migrates further into temperate continental Europe—A new alien aquatic plant for Ukraine. Feddes Repert. 133:305–312.

Fordyce, F. M., Everett, P. A., Bearcock, J. M., and Lister, T. R. (2019). The geosciences in Europe’s urban sustainability: Lessons from Glasgow and beyond (CUSP) Soil metal/metalloid concentrations in the Clyde Basin, Scotland, UK: implications for land quality. Earth Environ. Sci. Trans. R. Soc. Edinburgh 108:191–216.

Friedjung Yosef, A., Ghazaryan, L., Klamann, L., Kaufman, K. S., Baubin, C., Poodiack, B., Ran, N., Gabay, T., Didi-Cohen, S., Bog, M., et al. (2022). Diversity and differentiation of duckweed species from Israel. Plants 11:3326.

GBIF.org (2022). *Lemna* L. in GBIF Secretariat (2022). GBIF Backbone Taxonomy.

Hämälä, T., Moore, C., Cowan, L., Carlile, M., Gopaulchan, D., Brandrud, M. K., Birkeland, S., Loose, M., Kolář, F., Koch, M. A., et al. (2024). Impact of whole-genome duplications on structural variant evolution in the plant genus *Cochlearia*. Nat. C 15.

Janes, R. A., Eaton, J. W., and Hardwick, K. (1996). The effects of floating mats of *Azolla filiculoides* Lam. and *Lemna minuta* Kunth on the growth of submerged macrophytes. Hydrobiologia 340:23–26.

Kadono, Y., and Iida, S. (2022). Identification of a small, spring water-associated duckweed with special reference to the taxonomy of sect. uninerves of the genus *Lemna* (Lemnaceae) in Japan. Acta Phytotaxon. Geobot. 73:57–65.

Kirjakov, I. K., and Velichkova, K. N. (2016). Invasive species *Lemna* L. (Lemnaceae) in the flora of Bulgaria. Period. Biol. 118:131–138.

Kurihara, D., Mizuta, Y., Sato, Y., and Higashiyama, T. (2015). ClearSee: A rapid optical clearing reagent for whole-plant fluorescence imaging. Dev. 142:4168–4179.

Laird, R. A., and Barks, P. M. (2018). Skimming the surface: duckweed as a model system in ecology and evolution. Am. J. Bot. 105:1962–1966.

Lam, E. (2018). Available at: http://www.ruduckweed.org/ Accessed May 13, 2020.

Lam, E., and Michael, T. P. (2022). *Wolffia*, a minimalist plant and synthetic biology chassis. Trends Plant Sci. May;27:430–439.

Landesman, L., Fedler, C., and Duan, R. (2010). Plant nutrient phytoremediation using duckweed. In Eutrophication: Causes, Consequences and Control, pp. 341–354.

Landolt, E. (1986). Biosystematic investigations in the family of duckweeds (Lemnaceae). Vol. 1 & 2. Zürich: Geobotanisches Institut der Eidgenossiesche Technische Hochschule

Lansdown, R. V (2008). Red duckweed (*Lemna turionifera* Landolt) new to Britain. Watsonia 27:127–130.

Linton, S., and Goulder, R. (1998). The duckweed *Lemna minor* compared with the alga *Selenastrum capricornutum* for bioassay of pond-water richness. Aquat. Bot. 60:27–36.

McKenna, A., Hanna, M., Banks, E., Sivachenko, A., Cibulskis, K., Kernytsky, A., Garimella, K., Altshuler, D., Gabriel, S., Daly, M., et al. (2010). The genome analysis toolkit: A MapReduce framework for analyzing next-generation DNA sequencing data. Genome Res. 20:1297–1303.

Njambuya, J., Stiers, I., and Triest, L. (2011). Competition between *Lemna minuta* and *Lemna minor* at different nutrient concentrations. Aquat. Bot. 94:158–164.

O’Mahoney, R., Coughlan, N. E., Walsh, É., and Jansen, M. A. K. (2022). Cultivation of *Lemna minor* on industry-derived, anaerobically digested, dairy processing wastewater. Plants 11:1–13.

Ortega-Albero, N., González-Orenga, S., Vicente, O., Rodríguez-Burruezo, A., and Fita, A. (2023). Responses to salt stress of the interspecific hybrid *Solanum insanum* × *Solanum melongena* and its parental species. Plants 12:295.

Paolacci, S., Harrison, S., and Jansen, M. A. K. (2016). A comparative study of the nutrient responses of the invasive duckweed *Lemna minuta*, and the native, co-generic species *Lemna minor*. Aquat. Bot. 134:47–53.

Paolacci, S., Harrison, S., and Jansen, M. A. K. (2018a). The invasive duckweed *Lemna minuta* Kunth displays a different light utilisation strategy than native *Lemna minor* Linnaeus. Aquat. Bot. 146:8–14.

Paolacci, S., Jansen, M. A. K., and Harrison, S. (2018b). Competition between *Lemna minuta*, *Lemna minor*, and *Azolla filiculoides*. Growing fast or being steadfast? Front. Chem. 6:1–15.

Reynolds, M. P., and Braun, H. J. (2022). Wheat Improvement: Food Security in a Changing Climate. In: Reynolds, M.P., Braun, HJ. (eds). Springer, Cham. 10.1007/978-3-030-90673-3_1

Salt, D. E., Baxter, I., and Lahner, B. (2008). Ionomics and the study of the plant ionome. Annu. Rev. Plant Biol. 59:709–733.

Sattar, M. U., Anjum, F. M., and Sameen, A. (2015). Mitigation of heavy metals in different vegetables through biological washing techniques. Int. J. Food Allied Sci. 1:40.

Schindelin, J., Arganda-Carreras, I., Frise, E., Kaynig, V., Longair, M., Pietzsch, T., Preibisch, S., Rueden, C., Saalfeld, S., Schmid, B., et al. (2012). Fiji: an open-source platform for biological-image analysis. Nat. Methods 9:676–682.

Shah, A. A., Ahmed, S., Malik, A., Naheed, K., Hussain, S., Yasin, N. A., Javad, S., Siddiqui, M. H., Ali, H. M., and Ali, A. (2022). Potassium silicate and zinc oxide nanoparticles modulate antioxidant system, membranous H+-ATPase and nitric oxide content in faba bean (*Vicia faba*) seedlings exposed to arsenic toxicity. Funct. Plant Biol. 2023 Feb;50(2):146–159

Smith, K. E., Cowan, L., Taylor, B., McAusland, L., Heatley, M., and Murchie, E. H. (2024a). Physiological adaptation to high irradiance in duckweeds depends on light habitat niche and is ecotype and species-specific. J. Exp. Bot. 75:2046–2063.

Smith, K. E., Zhou, M., Flis, P., Jones, D., Bishopp, A., and Yant, L. (2024b). The evolution of the duckweed ionome mirrors losses in structural complexity. Ann. Bot. 133:997–1006.

Sree, K. S., and Appenroth, K.-J. (2020). Worldwide genetic resources of duckweed: Stock collections. In: Cao X.H., Fourounjian P., Wang W., editors. The Duckweed Genomes. Springer International Publishing; Cham, Switzerland: 2020. pp. 39–46.

Sree, K. S., Sudakaran, S., and Appenroth, K. J. (2015). How fast can angiosperms grow? Species and clonal diversity of growth rates in the genus *Wolffia* (Lemnaceae). Acta Physiol. Plant. 37.

Stout, L. M., Dodova, E. N., Tyson, J. F., and Nüsslein, K. (2010). Phytoprotective influence of bacteria on growth and cadmium accumulation in the aquatic plant *Lemna minor*. Water Res. 44:4970–4979.

Taghipour, E., Bog, M., Frootan, F., Shojaei, S., Rad, N., Arezoumandi, M., Jafari, M., and Salmanian, A. H. (2022). DNA barcoding and biomass accumulation rates of native Iranian duckweed species for biotechnological applications. Front. Plant Sci. 13:1–15.

Tran, N. B. T., Tran, T. N., and Hoang, T. N. P. (2022). Morphological variation, chromosome number, and DNA barcoding of giant duckweed (*Spirodela polyrhiza*) in Vietnam. Can Tho Univ. J. Sci. 14:61–67.

Van Echelpoel, W., Boets, P., and Goethals, P. L. M. (2016). Functional response (FR) and relative growth rate (RGR) do not show the known invasiveness of *Lemna minuta* (Kunth). PLoS One 11:1–18.

Volkova, P. A., Nachatoi, V. A., and Bobrov, A. A. (2023). Hybrid between *Lemna minor* and *L. turionifera* (*L. × japonica*, Lemnaceae) in East Europe is more frequent than parental species and poorly distinguishable from them. Aquat. Bot. 184:103593.

Walsh, É., Paolacci, S., Burnell, G., and Jansen, M. A. K. (2020). The importance of the calcium-to-magnesium ratio for phytoremediation of dairy industry wastewater using the aquatic plant *Lemna minor* L. Int. J. Phytoremediation 0:1–9.

Wang, Z., Jiang, Y., Bi, H., Lu, Z., Ma, Y., Yang, X., Chen, N., Tian, B., Liu, B., Mao, X., et al. (2021). Hybrid speciation via inheritance of alternate alleles of parental isolating genes. Mol. Plant 14:208–222.

Ware, A., Jones, D. H., Flis, P., Chrysanthou, E., Smith, K. E., Kümpers, B. M., Yant, L., Atkinson, J. A., Wells, D. M., Bhosale, R., et al. (2023). Loss of ancestral function in duckweed roots is accompanied by progressive anatomical simplification and a re-distribution of nutrient transporters. Curr. Biol. 33:1795–1802.

Washburn, J. D., and Birchler, J. A. (2014). Polyploids as a “model system” for the study of heterosis. Plant Reprod. 27:1–5.

Xu, Y., Ma, S., Huang, M., Peng, M., Bog, M., Sree, K. S., Appenroth, K. J., and Zhang, J. (2015). Species distribution, genetic diversity and barcoding in the duckweed family (Lemnaceae). Hydrobiologia 743:75–87.

Xu, J., Shen, Y., Zheng, Y., Smith, G., Sun, X. S., Wang, D., Zhao, Y., Zhang, W., and Li, Y. (2023). Duckweed (Lemnaceae) for potentially nutritious human food: A review. Food Rev. Int. 39:3620–3634.

Xue, C., Gao, Y., Qu, B., Tai, P., Guo, C., Chang, W., and Zhao, G. (2021). Hybridization with an invasive plant of *Xanthium strumarium* improves the tolerance of its native congener *X. sibiricum* to Cadmium. Front. Plant Sci. 12:1–14.

Zayed, A., Gowthaman, S., and Terry, N. (1998). Phytoaccumulation of trace elements by wetland plants: I. Duckweed. J. Environ. Qual. 27:715–721.

Zhang, X., Zhao, F. J., Huang, Q., Williams, P. N., Sun, G. X., and Zhu, Y. G. (2009). Arsenic uptake and speciation in the rootless duckweed *Wolffia globosa*. New Phytol. 182:421–428.

Zhang, H., Mittal, N., Leamy, L. J., Barazani, O., and Song, B. H. (2017). Back into the wild—Apply untapped genetic diversity of wild relatives for crop improvement. Evol. Appl. 10:5–24.

Ziegler, P., Adelmann, K., Zimmer, S., Schmidt, C., and Appenroth, K. J. (2015). Relative *in vitro* growth rates of duckweeds (Lemnaceae) - the most rapidly growing higher plants. Plant Biol. 17:33–41.

