## Supplemental figures 1-7 for "An ecological, phenotypic and genomic survey of duckweeds with their associated aquatic environments in the United Kingdom"

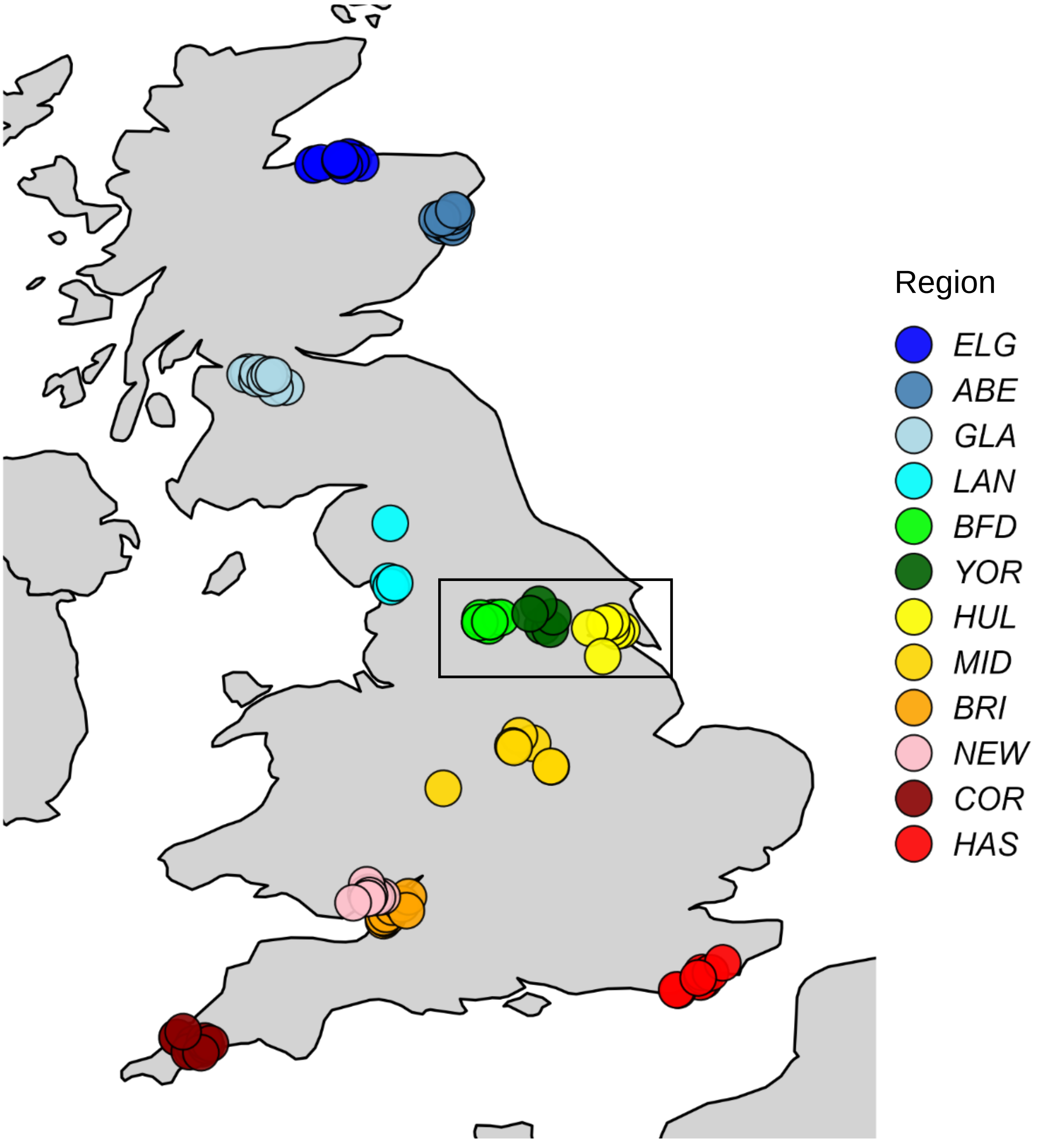


**Figure S1. Duckweed sites studied, spanning twelve regions in England, Scotland and Wales on a map of the UK.** Sites are coloured by regions and include from North to South: Elgin (ELG), Aberdeen (ABE), Glasgow (GLA), Lancashire (LAN), Bradford (BFD), York (YOR), Hull, (HUL) Midlands (MID), Bristol (BRI), Newport (NEW), Cornwall (COR), Hastings (HAS). The box marks a seasonal transect where duckweed plants were monitored and re-collected seasonally with longer term water assessments across 19 sites around BFD, YOR and HUL.

### Scotland & north west England

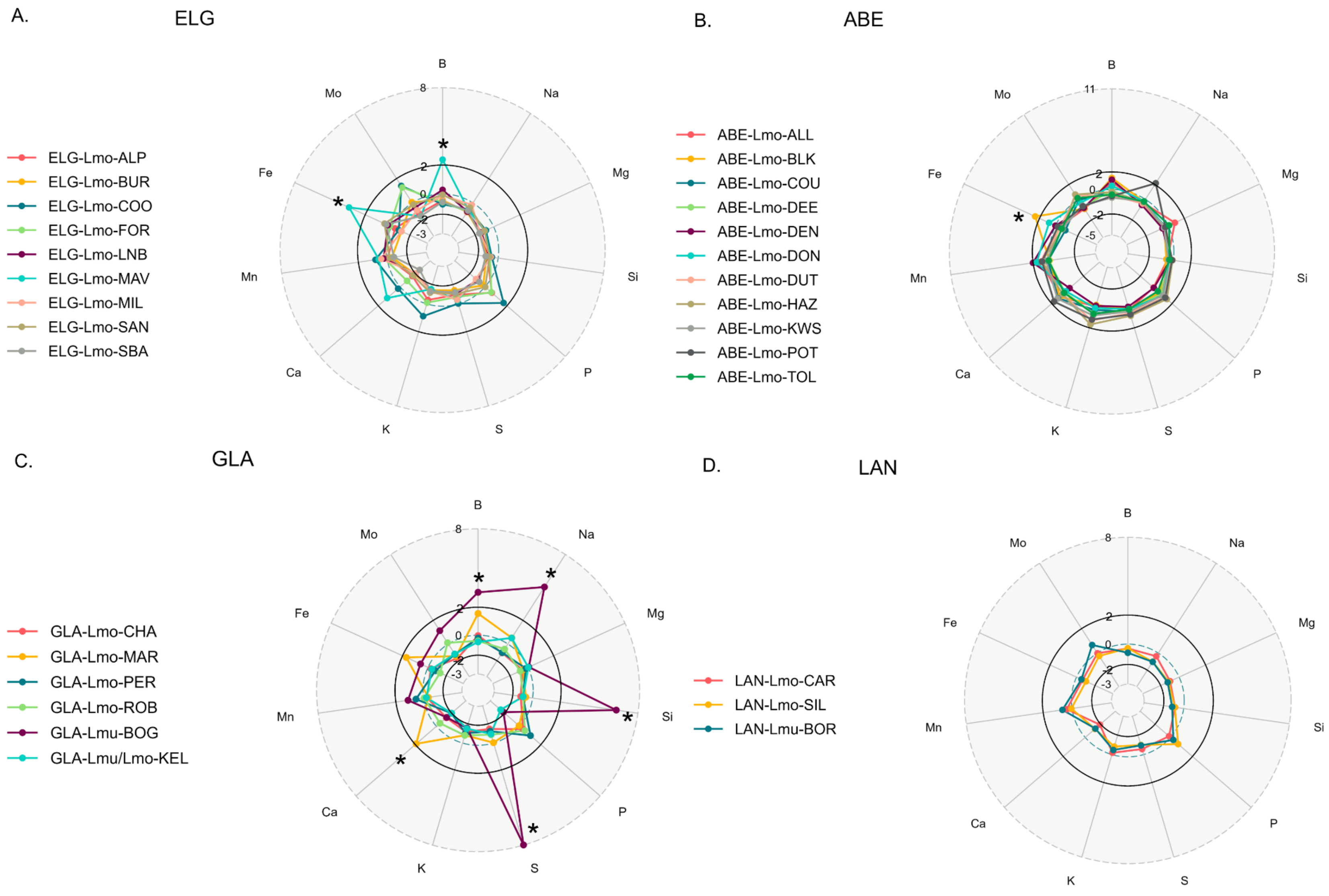

### North east – central England

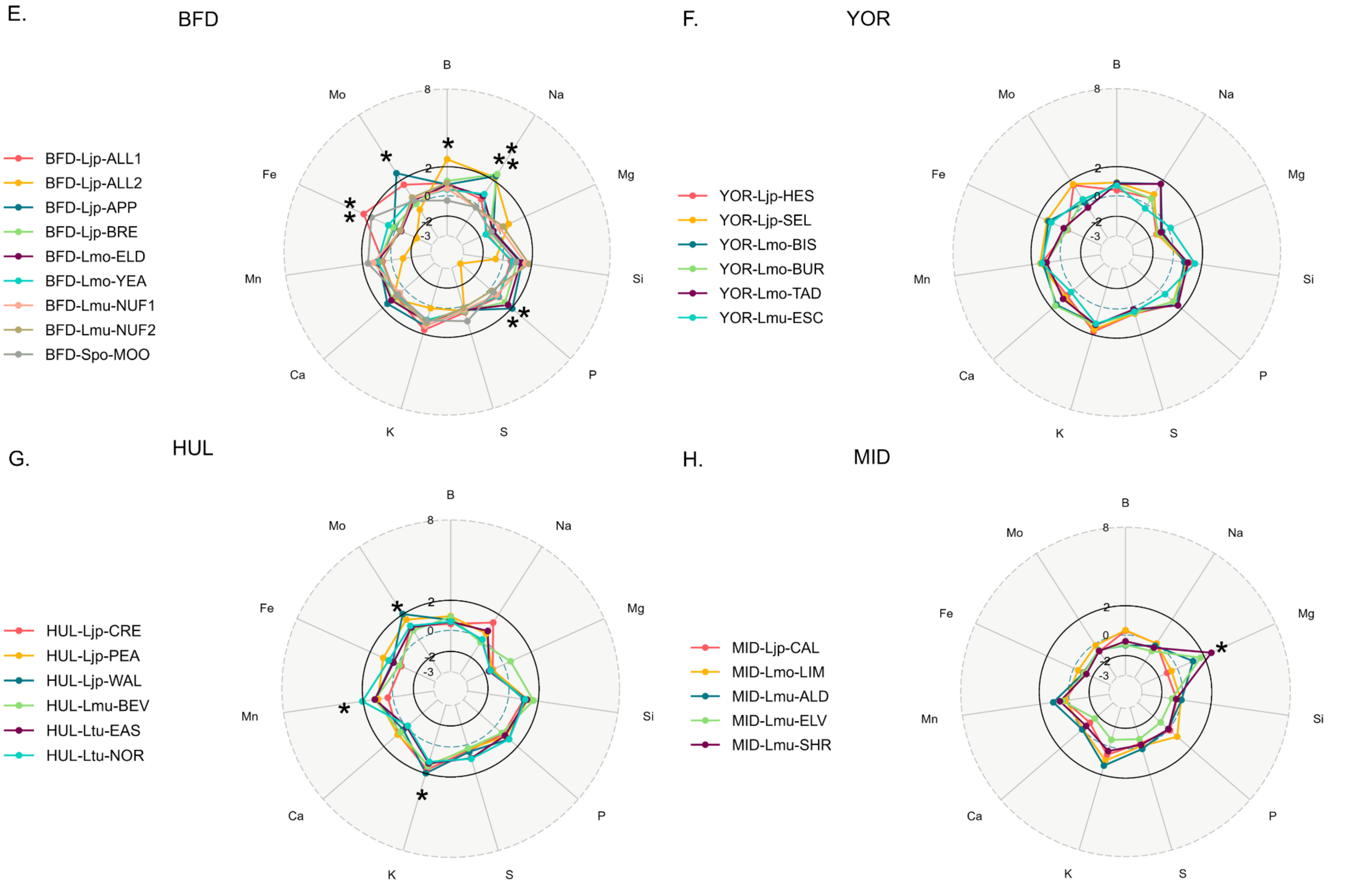

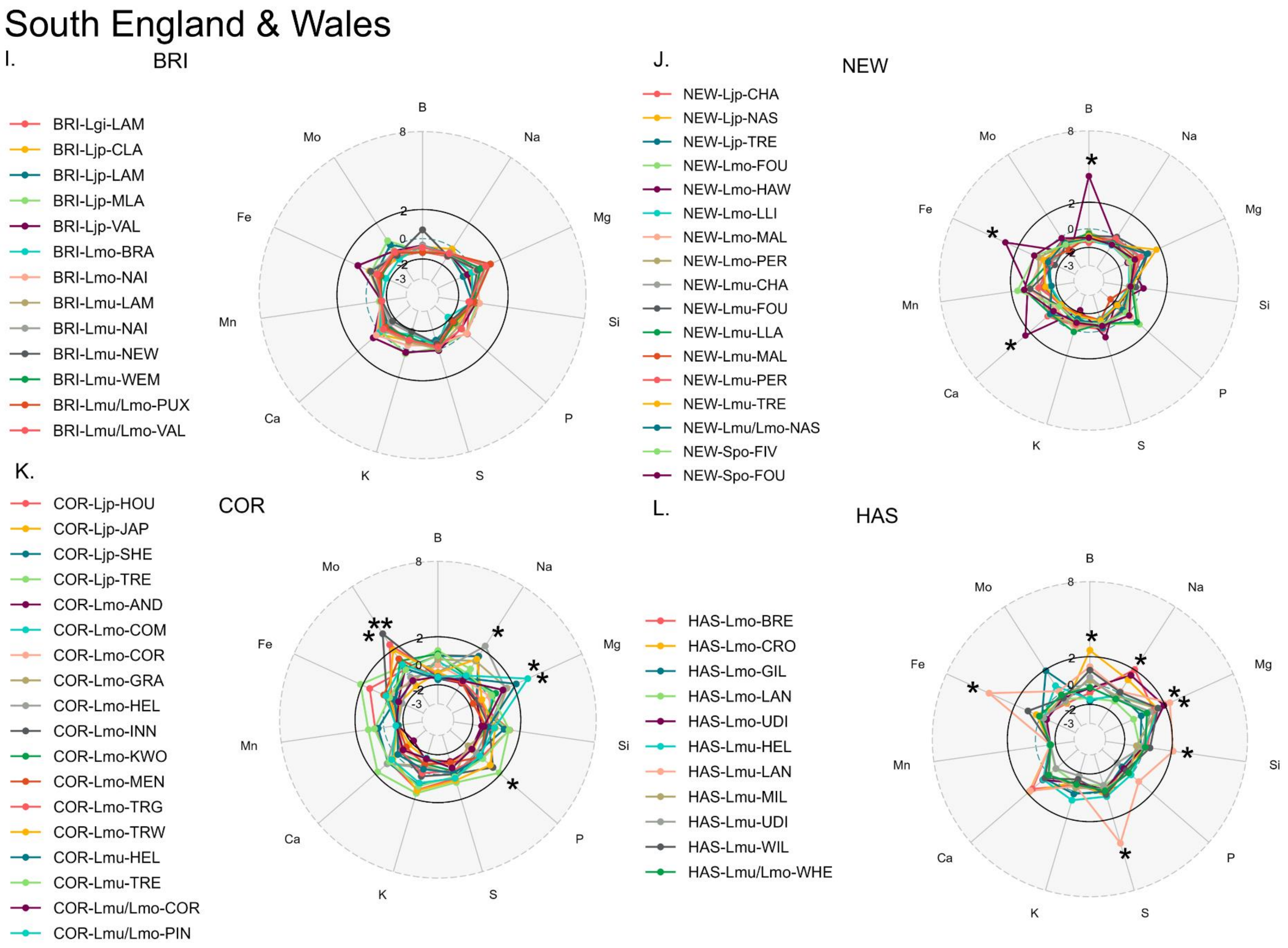
South England & Wales
I.
BRI
BRI-Lgi-LAM
BRI-Ljp-CLA
BRI-Ljp-LAM
BRI-Ljp-MLA
BRI-Ljp-VAL
BRI-Lmo-BRA
BRI-Lmo-NAI
BRI-Lmu-LAM
BRI-Lmu-NAI
BRI-Lmu-NEW
BRI-Lmu-WEM
BRI-Lmu/Lmo-PUX
BRI-Lmu/Lmo-VAL
B
Na
Mg
Si
P
S
K
Ca
Mn
Fe
Mo
8
2
0
-2
-3
J.
NEW
NEW-Ljp-CHA
NEW-Ljp-NAS
NEW-Ljp-TRE
NEW-Lmo-FOU
NEW-Lmo-HAW
NEW-Lmo-LLI
NEW-Lmo-MAL
NEW-Lmo-PER
NEW-Lmu-CHA
NEW-Lmu-FOU
NEW-Lmu-LLA
NEW-Lmu-MAL
NEW-Lmu-PER
NEW-Lmu-TRE
NEW-Lmu/Lmo-NAS
NEW-Spo-FIV
NEW-Spo-FOU
K.
COR
COR-Ljp-HOU
COR-Ljp-JAP
COR-Ljp-SHE
COR-Ljp-TRE
COR-Lmo-AND
COR-Lmo-COM
COR-Lmo-COR
COR-Lmo-GRA
COR-Lmo-HEL
COR-Lmo-INN
COR-Lmo-KWO
COR-Lmo-MEN
COR-Lmo-TRG
COR-Lmo-TRW
COR-Lmu-HEL
COR-Lmu-TRE
COR-Lmu/Lmo-COR
COR-Lmu/Lmo-PIN
L.
HAS
HAS-Lmo-BRE
HAS-Lmo-CRO
HAS-Lmo-GIL
HAS-Lmo-LAN
HAS-Lmo-UDI
HAS-Lmu-HEL
HAS-Lmu-LAN
HAS-Lmu-MIL
HAS-Lmu-UDI
HAS-Lmu-WIL
HAS-Lmu/Lmo-WHE

**Figure S2. Radar plots for elements quantified in duckweed ionomes from 12 UK regions grown in controlled conditions as measured by ICP-MS. A-D. Scotland and north west England. E-H. North east and central England. I-L. South Wales and England.** z-scores derived from normalised data for the whole duckweed panel and SD +/- 2 are considered significant. Elements are plotted at each point of the radar and include those provided in N-medium. The key corresponds to the independent site codes for each ecotype which is differentiated by its region, species and the site it was located, as indicated in Table S2A. At least *n*=3 ecotypes are included for each region.

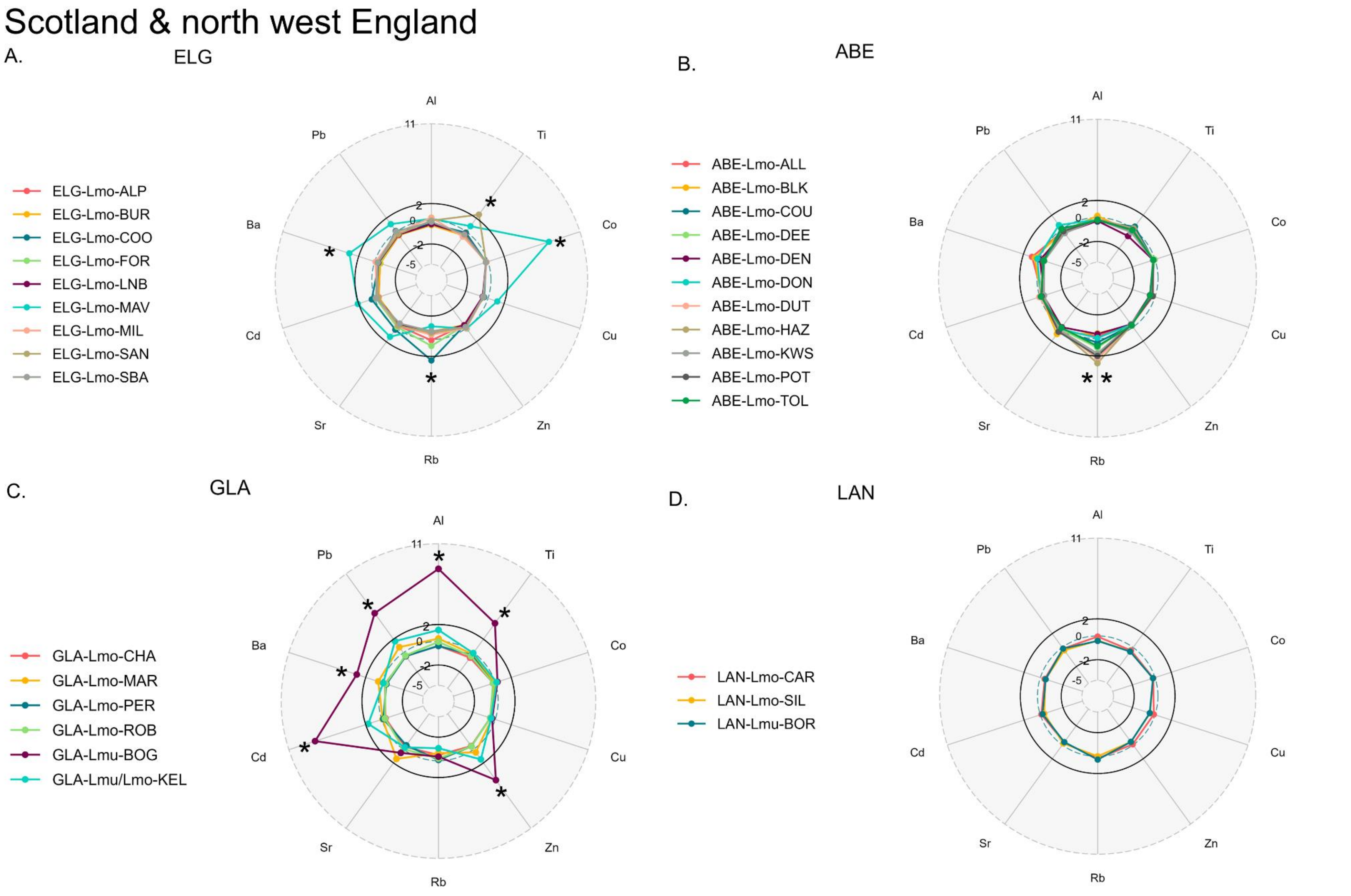

Scotland & north west England
A.
ELG
ELG-Lmo-ALP
ELG-Lmo-BUR
ELG-Lmo-COO
ELG-Lmo-FOR
ELG-Lmo-LNB
ELG-Lmo-MAV
ELG-Lmo-MIL
ELG-Lmo-SAN
ELG-Lmo-SBA
Al
Ti
Co
Cu
Zn
Rb
Sr
Cd
Ba
Pb
11
2
0
-2
-5
B.
ABE
ABE-Lmo-ALL
ABE-Lmo-BLK
ABE-Lmo-COU
ABE-Lmo-DEE
ABE-Lmo-DEN
ABE-Lmo-DON
ABE-Lmo-DUT
ABE-Lmo-HAZ
ABE-Lmo-KWS
ABE-Lmo-POT
ABE-Lmo-TOL
C.
GLA
GLA-Lmo-CHA
GLA-Lmo-MAR
GLA-Lmo-PER
GLA-Lmo-ROB
GLA-Lmu-BOG
GLA-Lmu/Lmo-KEL
D.
LAN
LAN-Lmo-CAR
LAN-Lmo-SIL
LAN-Lmu-BOR

### North east – central England

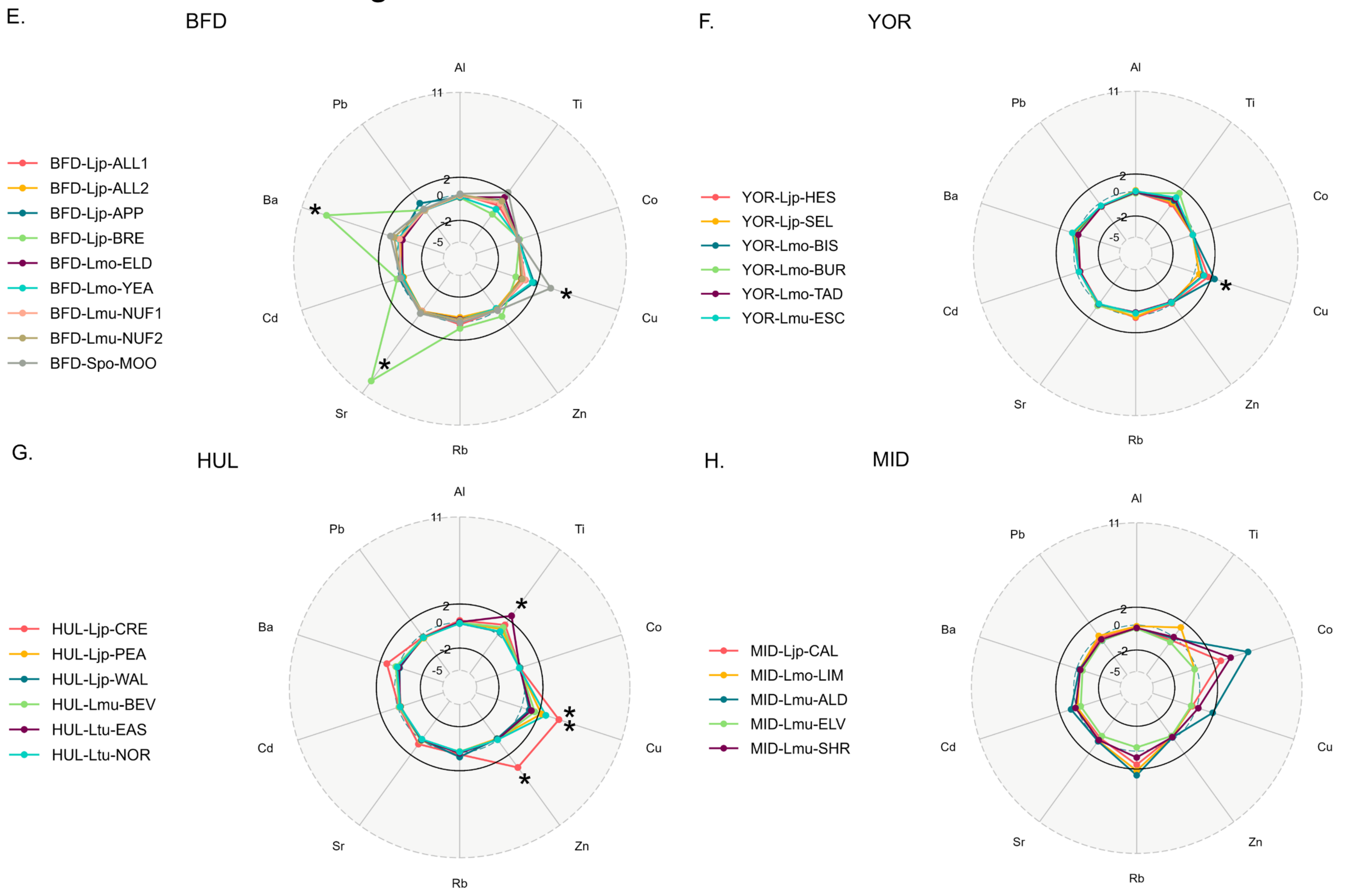

### South England & Wales

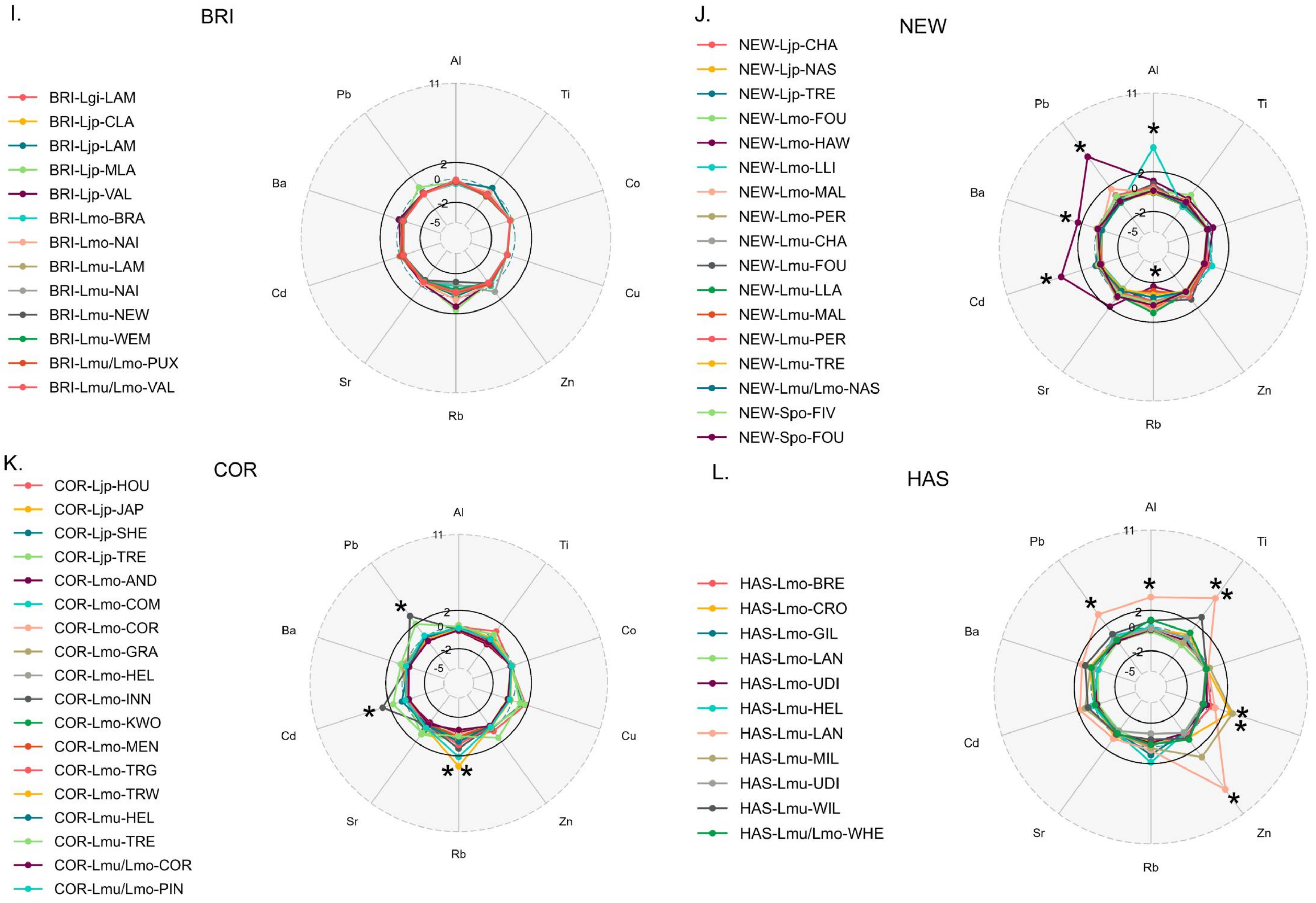

**Figure S3. Radar plots for heavy metal elements quantified in duckweed ionomes from 12 UK regions grown in controlled conditions as measured by ICP-MS. A-D. Scotland and north west England. E-H. North east and central England. I-L. South Wales and England.** z-scores are derived from normalised data for whole panel and SD +/- 2 are considered significant  The key corresponds to the independent site codes for each ecotype which is differentiated by its region, species and the site it was located, as indicated in Table S2A.

#### Scotland

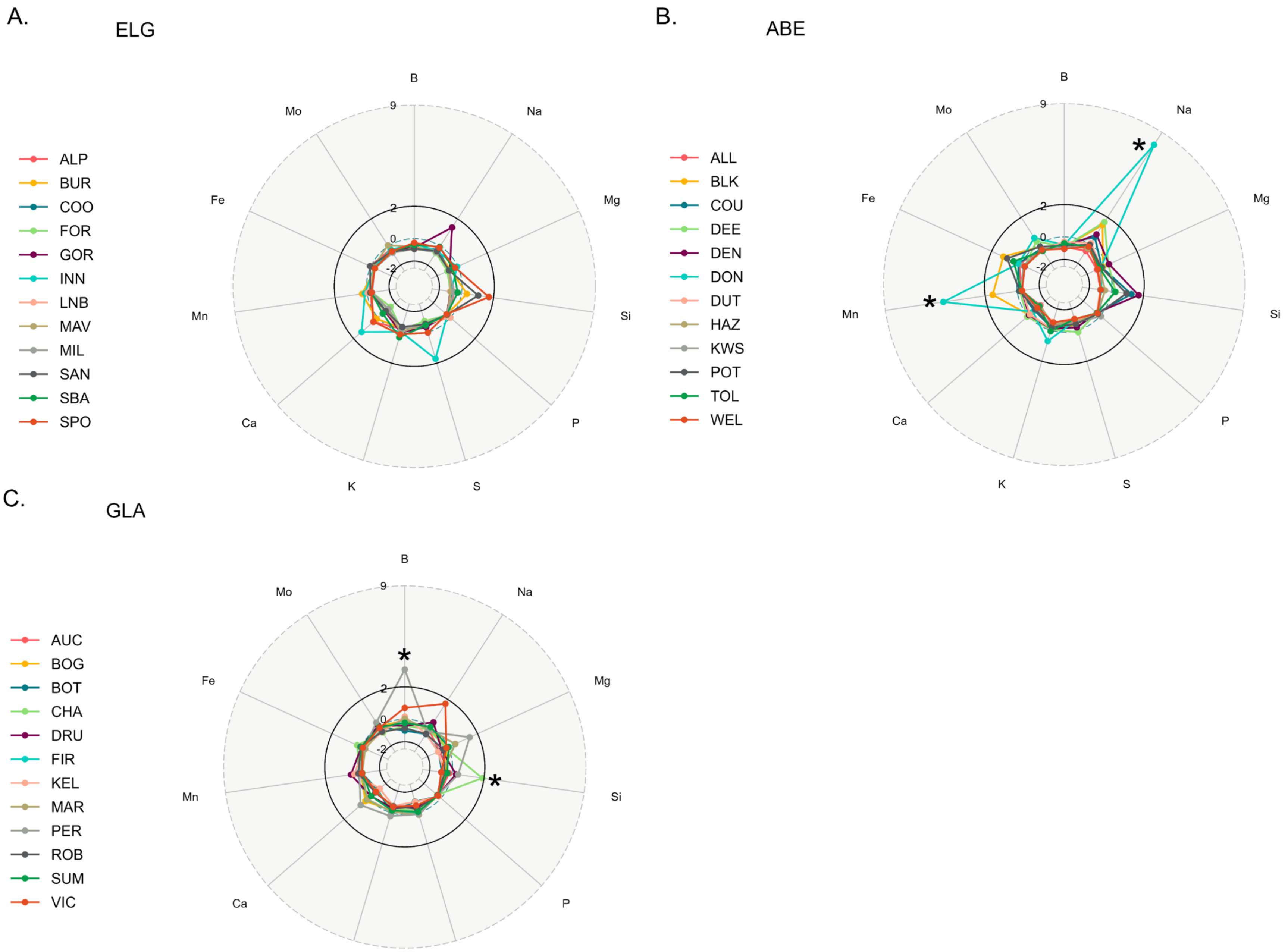
A.
ELG
B
Na
Mg
Si
P
S
K
Ca
Mn
Fe
Mo
9
2
0
-2
ALP
BUR
COO
FOR
GOR
INN
LNB
MAV
MIL
SAN
SBA
SPO
B.
ABE
*
*
ALL
BLK
COU
DEE
DEN
DON
DUT
HAZ
KWS
POT
TOL
WEL
C.
GLA
*
*
AUC
BOG
BOT
CHA
DRU
FIR
KEL
MAR
PER
ROB
SUM
VIC

### North east England

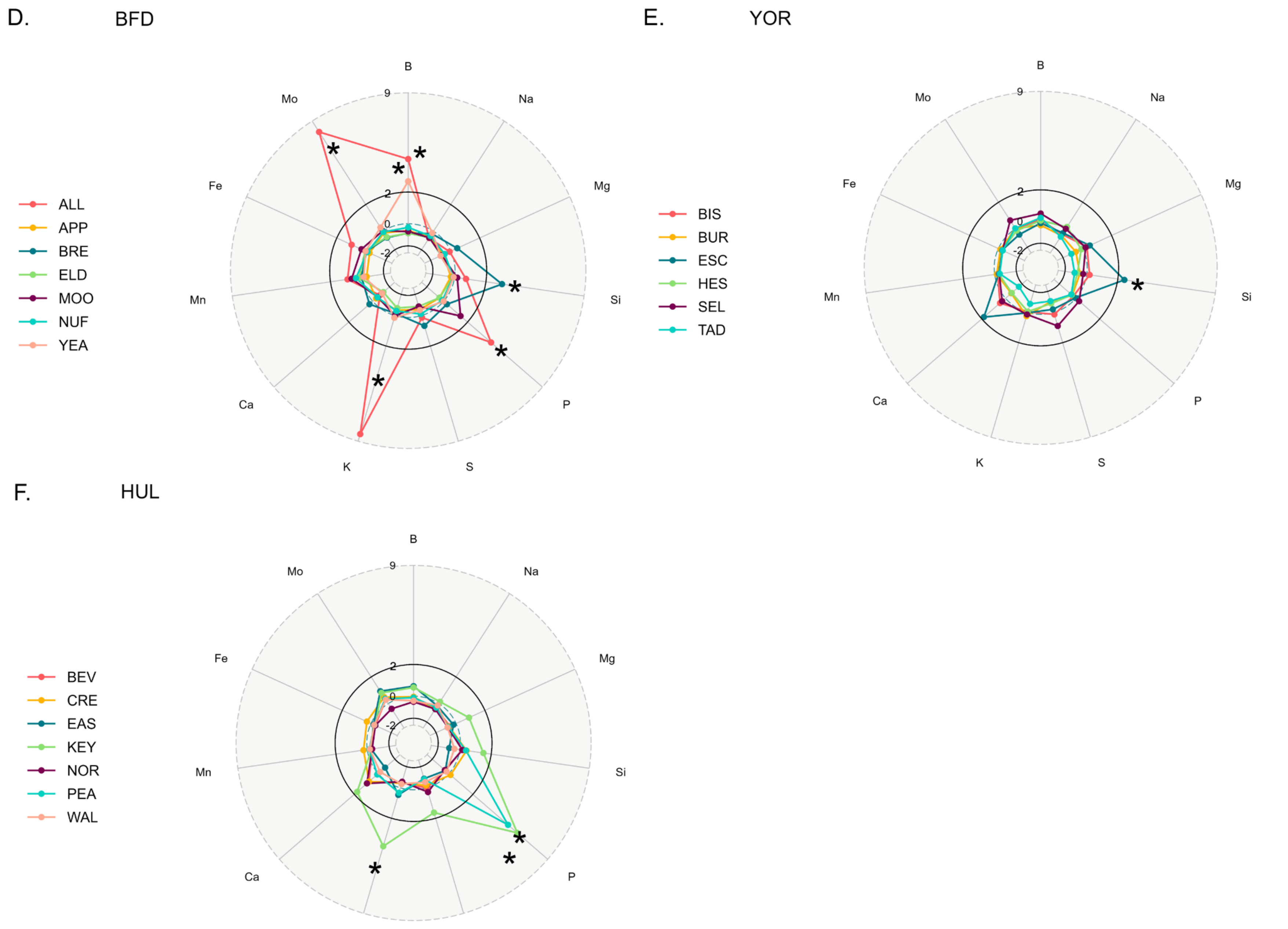

### South England & Wales

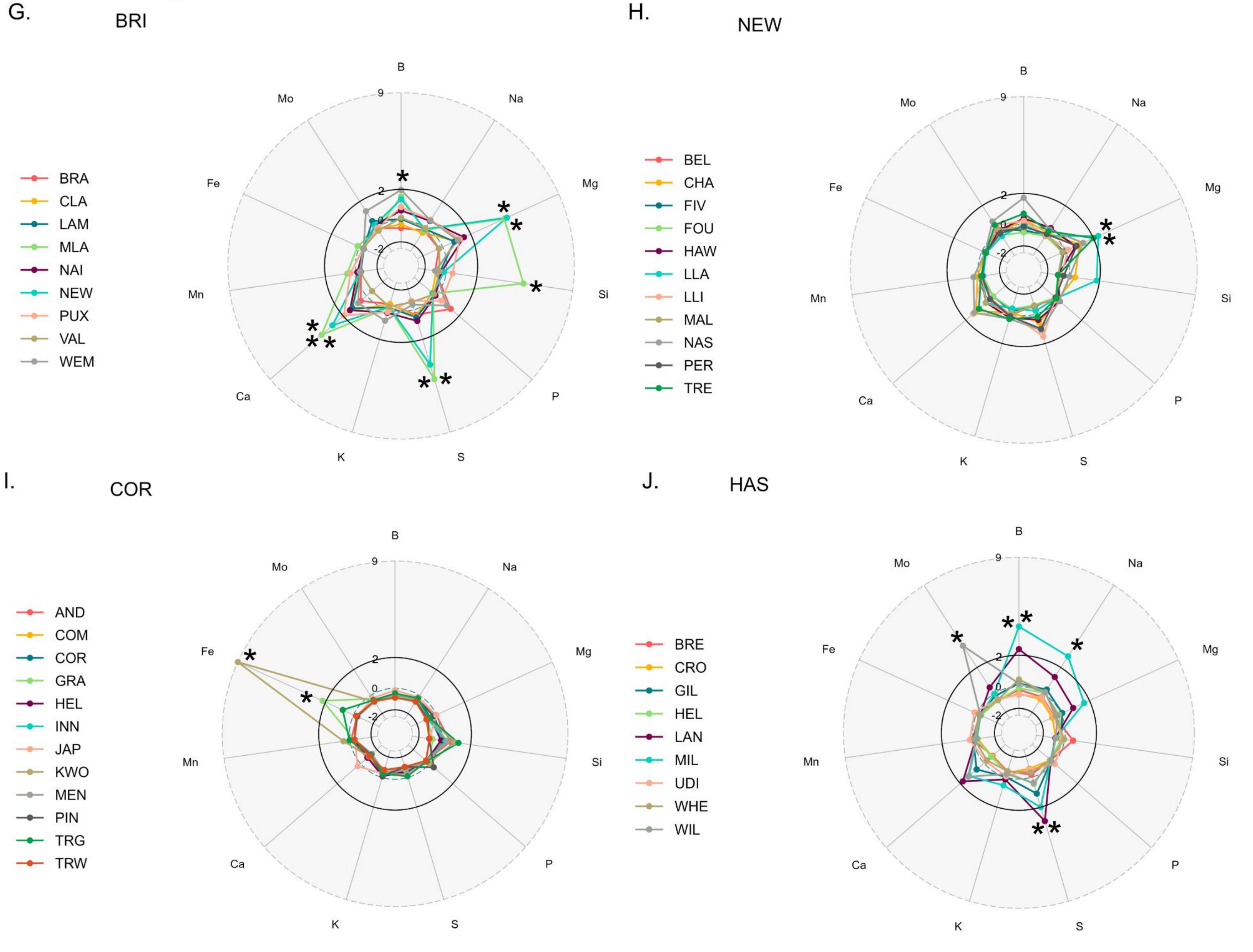

**Figure S4. Spatial variation for elements measured in water environments by ICP-MS. A-C. Scotland. D-F. North east England. G-J. South Wales and England.** Radar plots for macronutrients within individual water sites within ten UK regions. Macronutrients depicted at each point in the radar plot with relative concentration (z-scores) for each site coloured and named in the legend. The key corresponds to the site names in each region, as indicated in Table S1. All elements were present in duckweed lab-growth medium. Circles depict +2 or -2 of the SD for each population average and individual waters above or below this are considered extremes for that element.

### Scotland & north west England

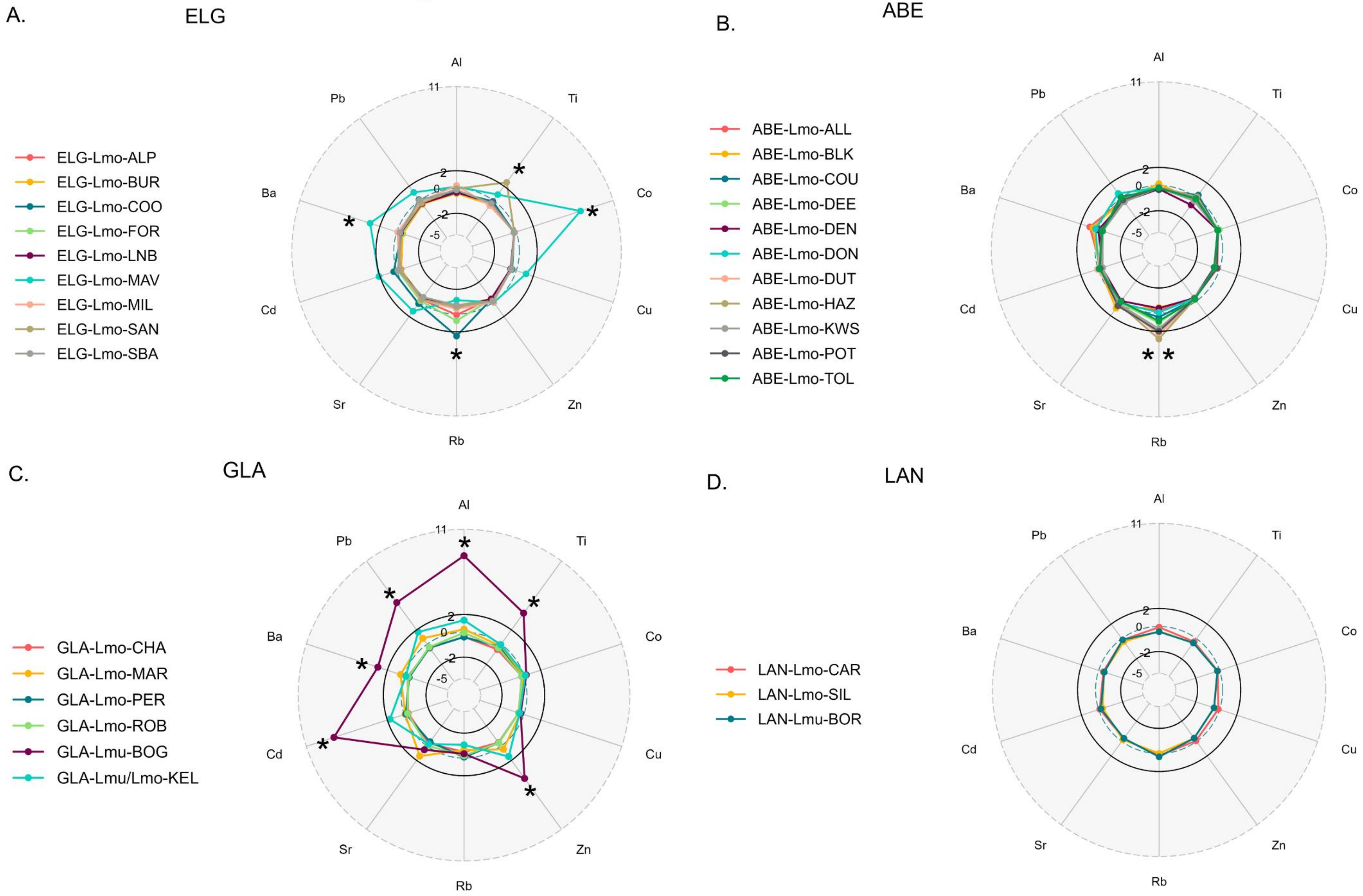

### North east England

D. BFD

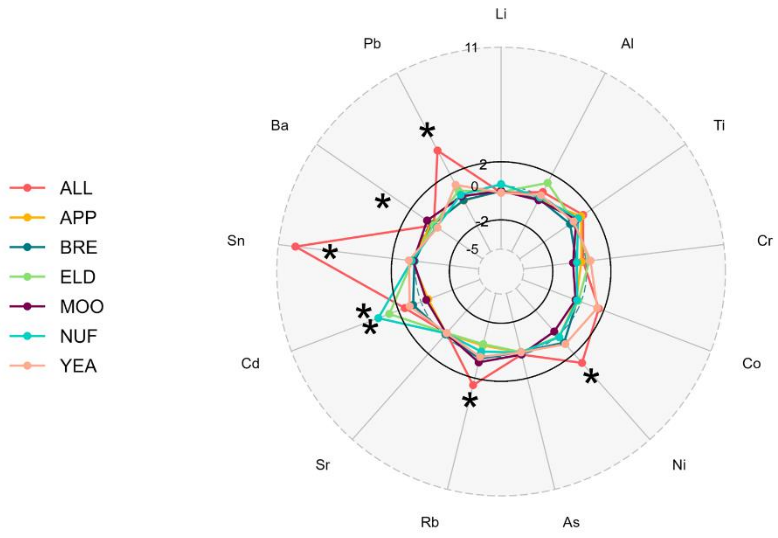


E. YOR

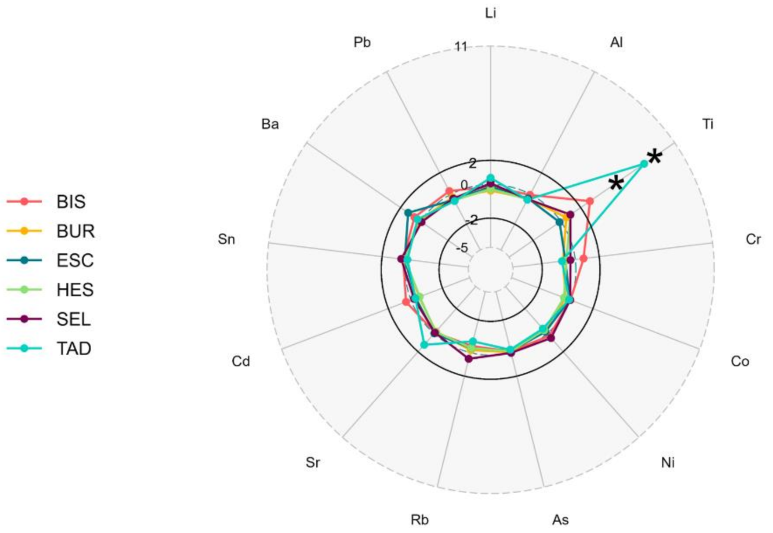


F. HUL

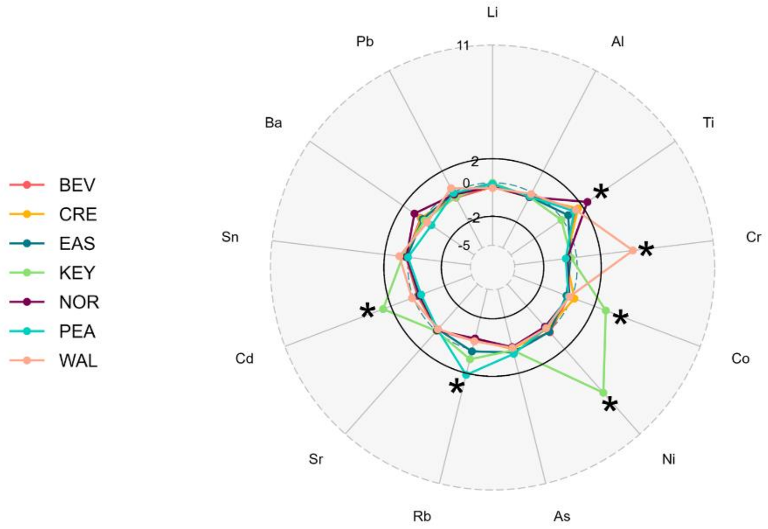

### South England & Wales

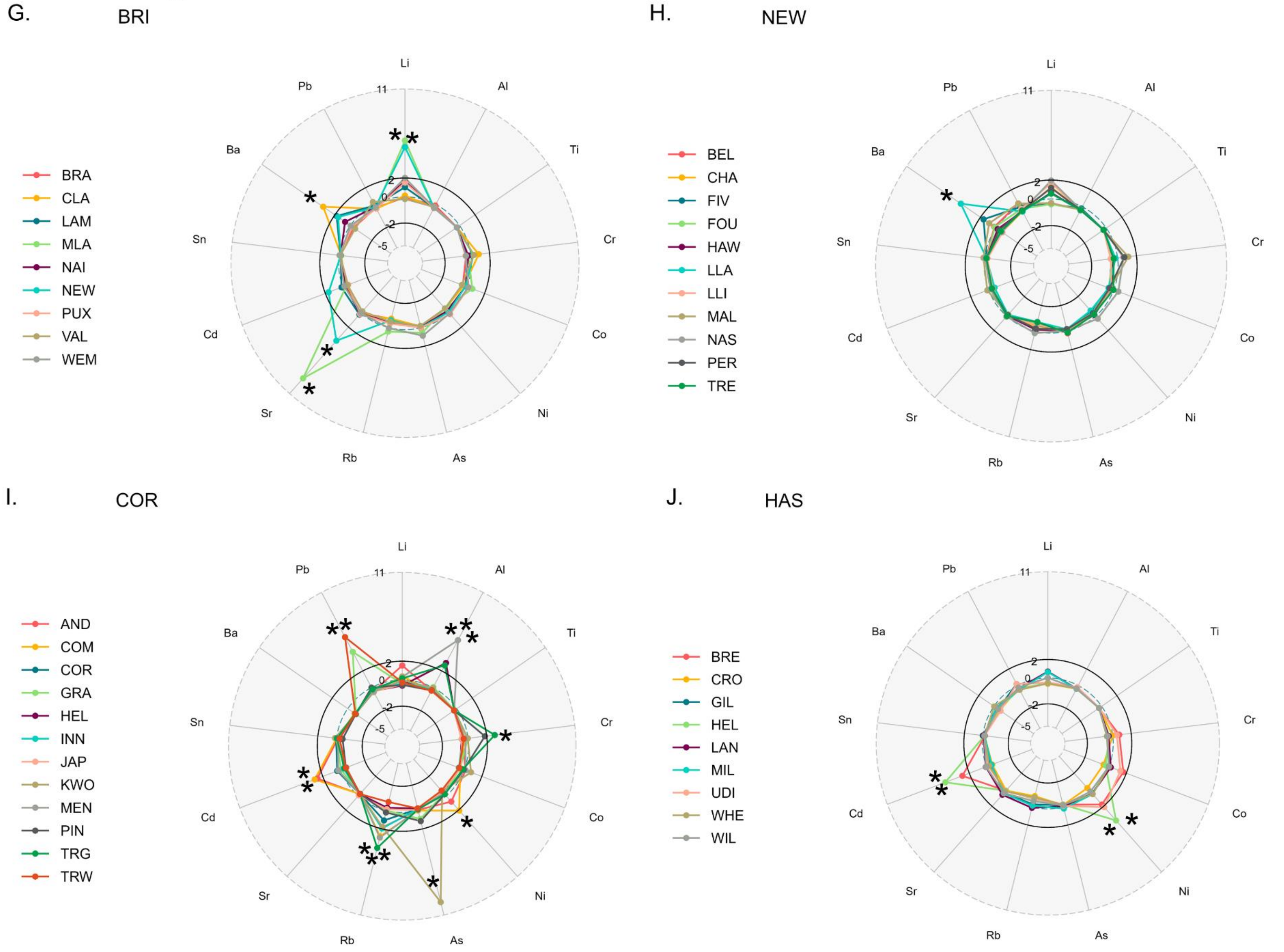

**Figure S5. Spatial variation for heavy metals measured in water environments in ten UK regions by ICP-MS. A-C. Scotland. D-F. North east England. G-J. South Wales and England.** The key corresponds to the site names in each region, as indicated in Table S1. Heavy metals at each axis of the radar plots and relative concentrations at each site are coloured and represented as lines from z-scores. Circles depict +2 or -2 of the SD for each population average and individual waters above or below this are considered extremes for that element.

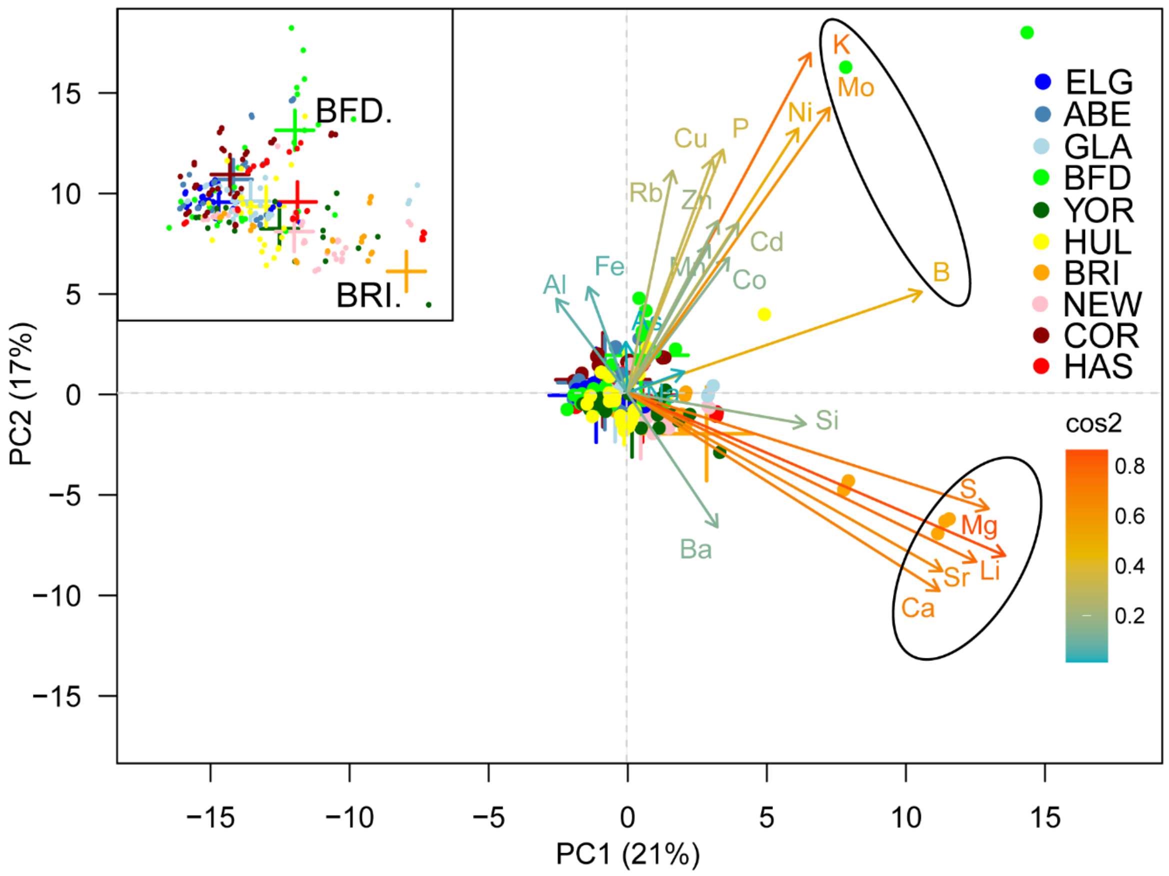


**Figure S6. Footprint of regional variation from 100 water environments showing elemental compositions measured by ICP-MS.** Individual sampling sites with water chemistry data including 21 elements plotted on PC1 and PC2. Ten collection regions with minimum of $n$ = 6 sites sampled per region are included. The arrows on the biplot are coloured by cosine (cos2) which shows elemental relationships and degree of contribution to data set variation. Circled regions correspond to elements associated with regions BFD and BRI. *Inset:* BFD and BRI show the most variation in water chemistry. Crosses depict the averages for ten regions, outliers from BRI and BFD have been removed to produce the close up inset figure.

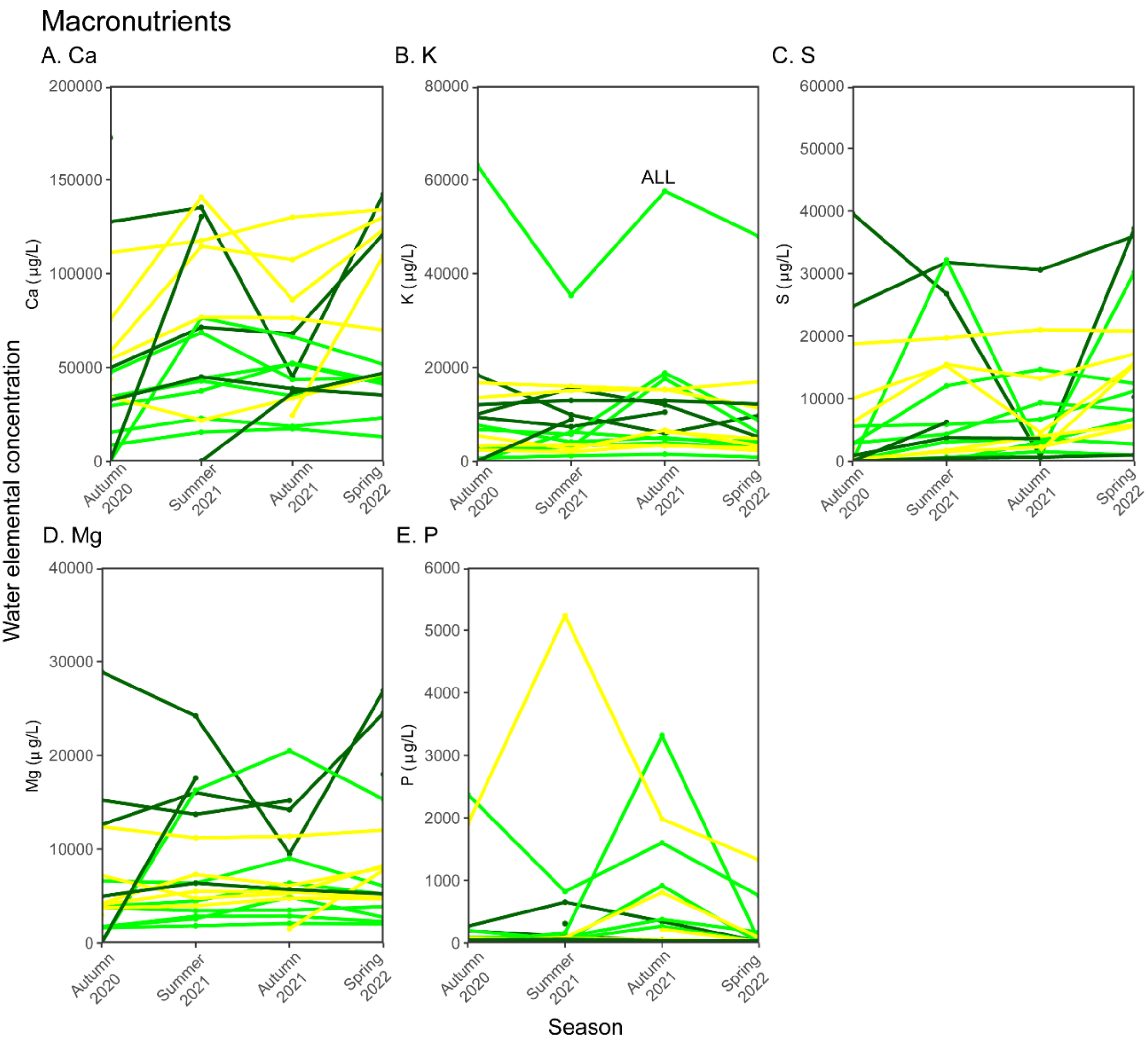

Macronutrients
A. Ca
B. K
C. S
D. Mg
E. P
Ca (µg/L)
K (µg/L)
S (µg/L)
Mg (µ g/L)
P (µg/L)
ALL
Water elemental concentration
Season
Autumn 2020
Summer 2021
Autumn 2021
Spring 2022

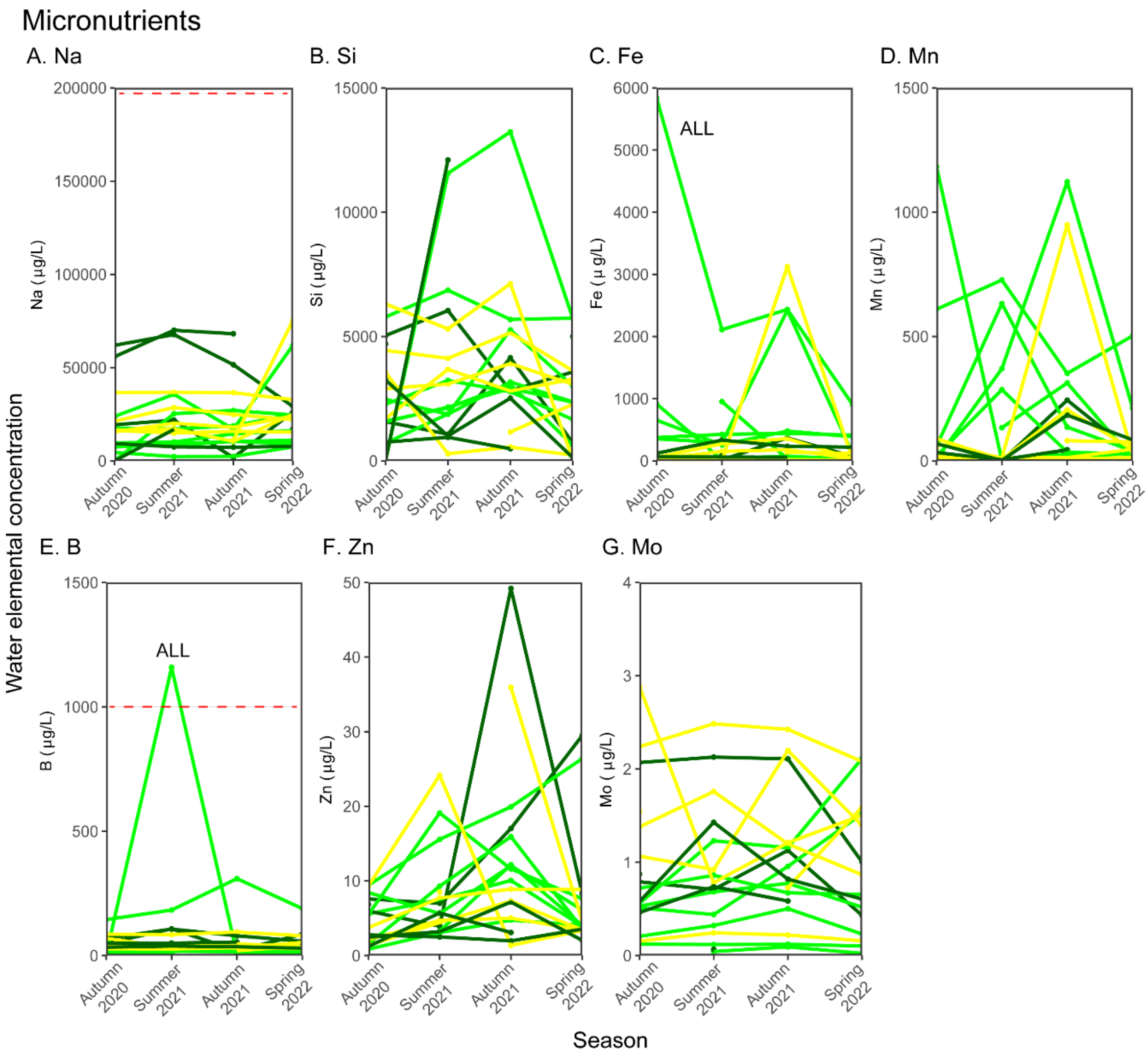
Micronutrients
A. Na
B. Si
C. Fe
D. Mn
E. B
F. Zn
G. Mo
ALL
ALL
Water elemental concentration
Na (µg/L)
Si (µg/L)
Fe (µg/L)
Mn (µg/L)
B (µg/L)
Zn (µg/L)
Mo (µg/L)
Autumn 2020
Summer 2021
Autumn 2021
Spring 2022
Season

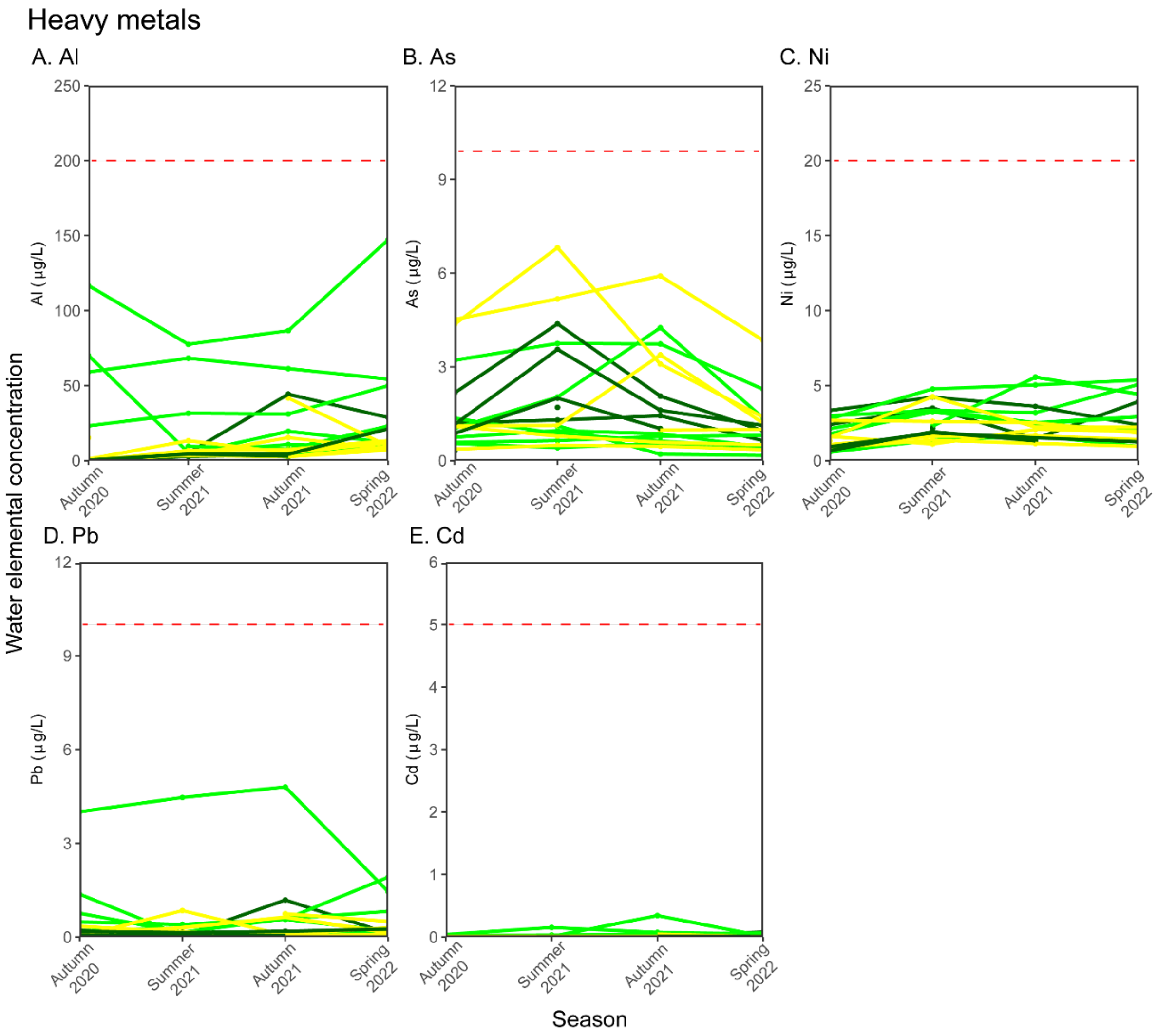


**Figure S7. Spatial and seasonal spikes in elemental concentrations in native water environments as measured by ICP-MS. Panel A. Macronutrients. A. Ca, B. K, C. S, D. Mg, E. P. Panel B. Micronutrients. A. Na, B. Si, C. Fe, D. Mn, E. B, F. Zn, G. Mn. Panel C. Heavy metals. A. Al, B. As, C. Ni, D. Pb, E. Cd.** Raw water elemental concentrations (in µg/L) are plotted as line plots overtime (autumn 2020, summer 2021, autumn 2021 and spring 2022). Three regions are plotted and lines are coloured by region: BFD (green), YOR (dark green), HUL (gold). Site ALL is labelled showing high and varied concentrations of K, Fe and B. The red dashed lines indicate the upper quantity limit allowed in drinking water for micronutrients Na and B and heavy metals Al, As, Ni, Pb and Cd using the Water quality regulations 2018.
