## Supplemental tables 1-7 for "An ecological, phenotypic and genomic survey of duckweeds with their associated aquatic environments in the United Kingdom"

**Table S1.** UK duckweed collection with native water sites details and descriptions.

| Site | Name | Region | Water source | Type | Location | Latitude | Longitude |
|---|---|---|---|---|---|---|---|
| KS02 | BIS | YOR | Ditch | Ditch under bridge with high banks | Cawood common, Bishop wood, North of Scalm lane, Selby | 53.79959 | -1.14745 |
| KS03 | ALL | BFD | Pond | Small 2 yr old shallow pond, shaded under 1 tree | Bradford valley | 53.81035 | -1.755023 |
| KS04 | SEL | YOR | Canal | Selby canal with parked boats | Selby canal | 53.77428 | -1.066171 |
| KS06 | NUF | BFD | Pond | Isolated pond, shallow depth | Nuffield health centre, Cottingley / manor pond at end of car park | 53.8307 | -1.819743 |
| KS09 | YEA | BFD | Small ponds | Tarnfield family ponds. Shallow | Yeadon ponds off tarn | 53.86976 | -1.670763 |
| KS12 | MOO | BFD | Pond | Stagnant pond | Bradford moor park | 53.80194 | -1.723388 |
| KS13 | ELD | BFD | Overflow stream off dam | Shallow overflow lake, some flow from dam to stream | Eldwick hall | 53.86671 | -1.815469 |
| KS14 | APP | BFD | Marina | Shallow marina with moored boats | Apperley bridge marina, Bradford | 53.83562 | -1.711316 |
| KS15 | BRE | BFD | Bog | Smallish bog/pond in public garden | Breary marsh, Golden acre park, Leeds | 53.87101 | -1.594412 |
| KS16 | EAS | HUL | Pond | Large pond, stagnant area after bridge | East park, East Hull | 53.76686 | -0.298782 |
| KS17 | PEA | HUL | Pond | Small pond, manmade fountains | Pearson park, Hull | 53.75825 | -0.355821 |
| KS18 | CRE | HUL | Beck | Shallow, non-flowing beck | Creyke beck, Cottingham | 53.78317 | -0.408387 |
| KS20 | BEV | HUL | Canal | Wide section of canal with fishing pegs | Beverley beck, Beverley | 53.83914 | -0.405169 |
| KS21 | WAL | HUL | Pond | Small village pond with walkway, high banking | Walkington, Beverley | 53.82131 | -0.482115 |
| KS22 | NOR | HUL | Beck | Downhill flowing beck | North cave beck, Brough | 53.78351 | -0.642291 |
| KS25 | ESC | YOR | Bog | Shallow bog in woodlands | Escwick woods bog, near York | 53.87598 | -1.034457 |
| KS27 | BUR | YOR | Pond | Medium-sized village pond | Burtree avenue pond, Skelton, York | 53.99645 | -1.132479 |
| KS28 | HES | YOR | Swamp | Large lake/swamp in woodland | Willow's fishery, Hessay, York | 53.97913 | -1.187243 |

| | | | | | | | |
|---|---|---|---|---|---|---|---|
| KS29 | TAD | YOR | River offshoot | Offshoot from river, stagnant one side, flowing at other | Offshoot river wharfe, access from Wighill lane, Tadcaster | 53.90209 | -1.281152 |
| KS33 | SIL | LAN | Ditch | Shallow bank side between reeds and wooden walkway | Leighton moss RSPB, Silverdale | 54.16884 | -2.798876 |
| KS34 | CAR | LAN | Canal | Lancaster canal near Carnforth | Carnforth, before Canal turn | 54.12518 | -2.770829 |
| KS35 | BOR | LAN | Canal | Lancaster canal between Tewitfield and Borwick | Between Tewitfield and Borwick | 54.15421 | -2.731501 |
| KS36 | PEN | LAN | Small circular pond | Small circular pond with bridge | Rheged service station, off A66, Penrith | 54.64717 | -2.779185 |
| KS37 | UON | MID | Dried up lake, arboretum | Shaded section of dried up lake | UON, Sutton bonington campus arboretum | 52.83363 | -1.253883 |
| KS38 | ELV | MID | Ditch | Lake off-shoot | Elvaston castle, Derbyshire | 52.89538 | -1.395662 |
| KS39 | LIM | MID | Bog | Small shallow muddy bog | Calke lime pits, Derbyshire | 52.81077 | -1.465801 |
| KS40 | CAL | MID | Cave pond | Sunken pond bricked area above | Grotto, Calke abbey, Derbyshire | 52.80043 | -1.45258 |
| AL01 | ALP | MID | Pond/bog in garden | Leaky 25 yr old garden pond fed by rainwater | Leicester LE5 2HU | 52.64 | -1.055 |
| AL02 | ALD | MID | Small steel container by overflow from water butt | Rainwater run off | Leicester | 52.64314 | -1.058691 |
| AL03 | SHR | MID | Ditch | Run off from roadside | Newport Shropshire | 52.46235 | -2.211213 |
| SS01 | SS01 | NCA | Garden pond | Garden pond | 10 Wolsey Court South Shields NE34 0QU | 54.97439 | -1.43347 |
| KS42 | HEL | HAS | Narrow stream | Shallow wooded flowing downhill | Helen's wood, Hastings | 50.88058 | 0.581676 |
| KS43 | GIL | HAS | Medium pond | Stagnant pond in woods with sloped landscape | Gilman's hill pond woods, St Leonards on Sea | 50.86504 | 0.543269 |
| KS44 | LAN | HAS | Sewer | Sewer fairly steeped banks with bridge | Langney sewer, Eastbourne | 50.79795 | 0.304928 |
| KS45 | WIL | HAS | Shallow ditch | Banked ditch with bridge, narrow beck minimal flow | Willingdon upper, shinewater park, Eastbourne | 50.8006 | 0.289478 |
| KS46 | WHE | HAS | Medium pond | Muddy bog land | Wheel lane, Westfield, Hastings | 50.91675 | 0.561538 |
| KS47 | BRE | HAS | Overflow dam | Reservoir overflow banked stagnant levels | Brede high woods, Battle | 50.94325 | 0.561655 |
| KS48 | UDI | HAS | Medium pond | Medium village stagnant pond | Udimore pond, Rye | 50.9405 | 0.651719 |
| KS49 | MIL | HAS | River | River with bridge/gate | Rother fishery, Rother river, Military road, Tenterden | 51.0232 | 0.782752 |
| KS50 | CRO | HAS | Small pond | Shallow pond with low flow, mud sides | Crowhurst park pond, Telham lane, Battle | 50.89911 | 0.520065 |

| | | | | | | | |
|---|---|---|---|---|---|---|---|
| KS51 | COR | COR | Shallow ditch | Shallow open ditch near river, old china clay harbour | Cornwall hotel, St Austell | 50.32616 | -4.795716 |
| KS52 | KWO | COR | Small sunken ditch | Shallow water and pipe opening into sunken tree area | Kingswood, St Austell | 50.31427 | -4.79936 |
| KS53 | GRA | COR | Shallow pond | Shallow grassy field pond, stagnant water, shaded by shrubbery | Grampound field, Truro | 50.29865 | -4.904628 |
| KS54 | MEN | COR | Medium pond | Spring-fed, clay-bottomed, stagnant no flow with island surrounded by plants | Menacuddle well pond, St Austell | 50.34607 | -4.795569 |
| KS55 | COM | COR | Medium lake | Farmyard lake on edge of farm next to streams, island in middle | Combe valley, Combe | 50.33426 | -4.879119 |
| KS56 | TRG | COR | Small sunken ditch | Small shallow sunken ditch walled on one side, no flow | Tregargus woods north, old china clay site, St Austell | 50.35485 | -4.883733 |
| KS57 | TRW | COR | Square garden pond | Square feature pond in walled garden, partial cover with net | Trewithen gardens, Truro | 50.2906 | -4.931766 |
| KS58 | PIN | COR | Medium pond | Medium pond, shallow with island and low flow | Pinetum gardens, Holmbush, St Austell | 50.34154 | -4.751426 |
| KS59 | INN | COR | Medium fishing pond | Medium fly-fishing pond, shallow at edges | Innis fly fishing, St Austell | 50.37814 | -4.766337 |
| KS60 | AND | COR | Large pond | Large park pond with flow | St Andrews Pond, St Andrew's Rd, Tywardreath, Par | 50.35764 | -4.706538 |
| KS61 | HEL | COR | Medium 'jungle' pond | ~100 year old pond in jungle garden area, spring-fed, shallow 3 ft | The Lost Gardens Heligan, B3273, Pentewan, Saint Austell PL26 6EN | 50.28258 | -4.808394 |
| KS62 | JAP | COR | Medium pond | Medium pond with bridge, waterfall | The Japanese garden, St Mawgan, Newquay, TR8 4ET | 50.45574 | -4.99803 |
| KS63 | PUX | BRI | Roadside ditch | Shallow ditch banked between road/cow farm, muddy/clay sides, near river | Puxton lane, Puxton, Hewish, Weston-Super-Mare, BS246TA | 51.372 | -2.851253 |
| KS64 | WEM | BRI | Roadside ditch | Some flow and depth, grass banks next to horse farm | Wemberham lane, Yatton | 51.38846 | -2.835666 |
| KS65 | CLA | BRI | Roadside ditch | Shallow ditch with bridge, grass banks next to sheep farm | Claverham drove rhyne, Claverham drove | 51.40839 | -2.804857 |
| KS66 | LAM | BRI | Roadside ditch | Ditch with some depth, no flow, embankment | Lampley rhyne, Clevedon | 51.39765 | -2.852591 |
| KS67 | NAI | BRI | Roadside ditch | Ditch with embankment | Blind yeo, Nailsea wall | 51.42396 | -2.821461 |
| KS68 | VAL | BRI | Medium pond | Stagnant medium sized park pond | Vale pond, Portishead | 51.48274 | -2.754449 |
| KS69 | NEW | BRI | Roadside ditch | Stagnant shallow ditch. Near marsh and motorways | Newlands rhyne | 51.50863 | -2.656918 |
| KS70 | MLA | BRI | Roadside ditch | Muddy ditch with little water, embankment, farm nearby, connects to river via sluice | Moor lane, Bristol | 51.56832 | -2.58513 |
| KS71 | BRA | BRI | Park ponds | Park ponds flowing downhill | Brandon hill, Bristol | 51.45379 | -2.606958 |

| | | | | | | | |
|---|---|---|---|---|---|---|---|
| KS72 | TRE | NEW | Roadside ditch | Ditch on side of road, bridge, no flowing water with embankment in industrial estate | Tesco depot, Magor, Newport | 51.56992 | -2.865184 |
| KS73 | LLI | NEW | Large pond | Large pond, open with fishing activity, some flow on one side. | Lliswerry pond, Newport | 51.58391 | -2.952568 |
| KS74 | NAS | NEW | Roadside ditch | Ditch on side of road/reen, half shaded, embanked | Lakes reen, Nash road, Newport | 51.56355 | -2.948225 |
| KS75 | CHA | NEW | Roadside ditch with drain pipe | Reen/ditch with opening, pipe, bridge, old steel works area, near Severn river | Chapel reen, Broad St Common, Whitson, Newport | 51.56428 | -2.916177 |
| KS76 | MAL | NEW | Shallow brook | Minimal flow, section trapped behind permanent lock, higher level, connects to canal network either side | Malpas brook, Bettws, Newport | 51.60444 | -3.011118 |
| KS77 | FOU | NEW | Canal locks | Gentle flow, shallow, section between open and closed locks | Fourteen locks, Cwm lane, Rogerstone | 51.59157 | -3.040473 |
| KS78 | FIV | NEW | Canal locks | Gentle flow, shallow, mud edge sectioned off from main water body, water containing high sediment | Monmouth canal, five locks, Cwmbran | 51.66664 | -3.031435 |
| KS79 | BEL | NEW | Small park pond | Pond, different levels downstream connected by sections and bridge | Belle vue park, Newport | 51.57849 | -3.000884 |
| KS80 | PER | NEW | Ditch system | Reen, Ditch, muti-directions, shallow, gentle flow near edges, some stagnant | Percoed reen, Duffryn way, Duffryn, Newport | 51.55498 | -3.01977 |
| KS81 | HAW | NEW | Roadside ditch | Reen, ditch. Not embanked. Some depth in water, flow, lock/gated mechanism | Percoed reen, Hawse lane, Cardiff | 51.55498 | -3.01977 |
| KS82 | LLA | NEW | Shallow pond | Wooded shallow pond middle of circular road, stagnant, mud sediment. | Llandenis oval, Cardiff | 51.51842 | -3.177074 |
| KS83 | PER | GLA | Shallow pond | Shallow, shaded by reeds/long grass | Smaller perchy pond, Wishaw, Glasgow | 55.77158 | -3.899133 |
| KS84 | CHA | GLA | Park pond | Country park, medium pond, shallow edges, some depth | Chatelherault country park, Hamilton, Glasgow, ML3 7UE | 55.76391 | -4.014075 |
| KS85 | KEL | GLA | Park pond | Medium sized pond, shallow muddy water, island in middle, some flow | Kelvingrove park, Glasgow, G3 7SD | 55.8692 | -4.284518 |
| KS86 | BOT | GLA | Small man-made pond | Small shallow pond, some flow, mostly open | Glasgow botanical garden, Glasgow | 55.87847 | -4.288534 |
| KS87 | FIR | GLA | Canal overflow area | Canal link area with fishing pegs around, some flow, open, near clay pits | Firhill court, Glasgow | 55.88263 | -4.270283 |
| KS88 | MAR | GLA | Canal locks | Lock network, with flow, catchment pool area | Maryhill locks, Glasgow, G4 9SP | 55.89368 | -4.297603 |
| KS89 | VIC | GLA | Large boating lake | Park boating lake with bridge | Victoria Park, Glasgow, G14 9NW | 55.87526 | -4.332853 |
| KS90 | AUC | GLA | Ditch/drain system? | Sunken with bridge, in middle of housing | Auckland Wynd, Glasgow G40 4RN | 55.84076 | -4.20635 |

| | | | | | | | |
|---|---|---|---|---|---|---|---|
| KS91 | ROB | GLA | Small pond | Wetlands sunken shallow small pond | Robroyston Park, 220 Robroyston Rd, Glasgow G33 1JQ | 55.88944 | -4.19682 |
| KS92 | BOG | GLA | Marsh | Marsh near housing with embankment | Boghall road, Uddingston, Glasgow, G71 | 55.84264 | -4.11906 |
| KS93 | DRU | GLA | Large loch | Large loch in country park | Loch lochend, Drumpellier country park, Coatbridge, ML5 2EH | 55.87271 | -4.075 |
| KS94 | SUM | GLA | Canal system | Canal with some flow, some depth, old steel/iron site | Summerlee industrial museum, Heritage Way, Coatbridge ML5 1QD | 55.86568 | -4.03001 |
| KS95 | SBA | ELG | Woodland ditch | Shaded ditch lined with trees, shallow water | Spey Bay, Fochabers IV32 7PJ | 57.66333 | -3.06098 |
| KS96 | SPO | ELG | Burn (stream) | Stream connecting up and flowing to sea, some flow, some depth, bridge and embankment | Dry Burn, Spey portgorden | 57.66286 | -3.04523 |
| KS97 | ALP | ELG | Large farm pond | Large pond with some depth, inlet/outlet from Mosstowie canal, trees with rill drain system, 20 yr old | Mossend Farm, Mosstowie, Elgin IV30 8TU | 57.62975 | -3.41362 |
| KS98 | FOR | ELG | Medium-sized pond | Shallow sunken bog with embankment | Forres golf course, Forres, IV36 2RD | 57.60876 | -3.59307 |
| KS99 | SAN | ELG | Large lake | Large lake with network of feeders/waterfall. Has island and bridge/jetty | Sanquhar Loch, Forres IV36 | 57.60191 | -3.60725 |
| KS100 | BUR | ELG | Large pond | Large open pond, some depth in woodland arboretum garden | Burgie Arboretum Woodland Garden, Burgie Estate, Forres IV36 2QU | 57.62021 | -3.52381 |
| KS101 | MAV | ELG | Small pond | Small pond (apparently fed by other small pond via inlet) | Maverston golf course, Garmouth Road, Elgin IV30 8LR | 57.65437 | -3.17686 |
| KS102 | INN | ELG | Medium pond | Open blue pond, flowing, some depth with an island | Innes pond, Innes house, Lochhill, Elgin IV30 8NG | 57.66735 | -3.22723 |
| KS103 | GOR | ELG | Large pond | Large pond in estate with flow and depth | Gordon Castle Lake, Gordon Castle Estate, Fochabers IV32 7PQ | 57.61726 | -3.09553 |
| KS104 | LNB | ELG | Large loch | Large loch in estate with flow and depth, trees two sides, open | Loch na Bo, Elgin IV30 8QY | 57.62752 | -3.20042 |
| KS105 | MIL | ELG | Large loch | Large lake (with shallow stream attached), some flow, two sides tree-lined other open | Milbuies loch, Elgin | 57.5954 | -3.270719 |
| KS106 | EDG | ELG | Ditch in wetlands | Narrow stream some flow with embankment. Old peat bog now wetlands | Ward's wildlife site, Edgar road, Elgin, IV30 | 57.64044 | -3.319449 |

| | | | | | | | |
|---|---|---|---|---|---|---|---|
| KS107 | COO | ELG | Medium pond | Medium sized pond, with flow, open pond in park with island | Cooper's park, Elgin, IV30 1HS | 57.65186 | -3.3146 |
| KS108 | TOL | ABE | Medium pond | Medium stagnant pond in woods. Some trees, treefall | Tollohill wood, Aberdeen, AB12 5XN | 57.11141 | -2.12911 |
| KS109 | DEE | ABE | Overflow pond | Pipe inlet/outlet with some flow | Deeside pond, Off station road, Bucksburn, Aberdeen | 57.10089 | -2.235427 |
| KS110 | ALL | ABE | Large pond | Large pond in park with trees around, open one side jetty with low water level, high sediment, | Allan park pond, Park Brae, Cults, Aberdeen, AB15 9HS | 57.11361 | -2.17897 |
| KS111 | HAZ | ABE | Medium pond | Stagnant medium pond covered in plant matter, pipe inlet/outlet | Hazeldene road pond, Hazlehead, Aberdeen, AB15 | 57.13494 | -2.174375 |
| KS112 | COU | ABE | Medium pond | Shallow pond for fishing, with inlet, some flow | Couper's pond, 3 Macaulay Gardens, Hazlehead, Aberdeen, AB15 8FN | 57.13441 | -2.15543 |
| KS113 | WEL | ABE | Large pond | Large pond with flow and depth | Wellington road pond, AB12 | 57.0871 | -2.111919 |
| KS114 | DUT | ABE | Pond network | Pond network in public park. Linked lakes, stagnant, bridges and islands | The Linked Lakes, Duthie park, Aberdeen, AB11 7BH | 57.12937 | -2.1056 |
| KS115 | DON | ABE | Pollution control dam | Medium-sized dam, near river. Mud/shallow water,middle - grass islands and flow, with sediment | Donside pond, Gordon Brae, Danestone, Aberdeen, AB228BN | 57.17694 | -2.118509 |
| KS116 | BLK | ABE | Medium pond | Medium with flow. Some depth, with sediment, near burn | Blackdog burn pond, Blackdog, Aberdeen | 57.21679 | -2.072611 |
| KS117 | DEN | ABE | Medium pond | Medium pond, stagnant in public park, oil in water | Denman park pond, Westhill AB32 | 57.15189 | -2.27732 |
| KS118 | KWS | ABE | Medium pond | Medium pond, stagnant, jetty with low flow | Kingswells pond, Fairley, Aberdeen | 57.16412 | -2.21807 |
| KS119 | POT | ABE | Narrow stream | Narrow, shallow stream/burn, embankment, shallow, low flow. | Potterton park burn, Aberdeen AB23 8UG | 57.22992 | -2.09853 |
| KS120 | KEY | HUL | Ditch | Ditch under bridge. High banks | Keyingham drain, Keyingham | 53.71046 | -0.152052 |
| MP01 | ABB | NCA | Pond | Small garden pond | Wolsey Court South Shields NE34 0QU | 54.97439 | -1.43347 |
| LY01 | TRE | COR | Pond | Medium-sized pond | Trencreek holiday park | 50.40796 | -5.06188 |
| LY02 | SHE | COR | Stream | Slow flowing, fresh steam | Sherford stream, off Sherford road | 51.00341 | -3.10177 |
| LY03 | HOU | COR | Stream | Small stream no flow | Housel bay | 49.96474 | -5.19621 |
| SS01 | SS01 | SHA | Garden pond | Garden pond | 10 Wolsey Court South Shields NE34 0QU | 54.97439 | -1.43347 |

**Table S2A.** UK accessions and previously characterised clones newly sequenced in this study.

| Accession | Site code | Species | Latitude | Longitude | Registered clone | SRA project | SRA sample | SRA clone |
|---|---|---|---|---|---|---|---|---|
| KS02 | YOR-Lmo-BIS | *Lemna minor* | 53.799 | 1.147 | *Lemna minor* 5882 | PRJNA1026139 | SAMN37735463 | SRR26858637 |
| KS03 | BFD-Ljp-ALL1 | *Lemna japonica* | 53.81 | 1.755 | *Lemna japonica* 5883 | PRJNA1026139 | SAMN37735464 | SRR26858636 |
| KS04 | YOR-Ljp-SEL | *Lemna japonica* | 53.774 | 1.066 | *Lemna japonica* 5884 | PRJNA1026139 | SAMN37735465 | SRR26858625 |
| KS06A | BFD-Lmu-NUF1 | *Lemna minuta* | 53.83 | 1.819 | *Lemna minuta* 5885 | PRJNA1026139 | SAMN37735466 | SRR26858620 |
| KS06B | BFD-Lmu-NUF2 | *Lemna minuta* | 53.83 | 1.819 | *Lemna minuta* 5886 | PRJNA1026139 | SAMN37735467 | SRR26858619 |
| KS09 | BFD-Lmo-YEA | *Lemna minor* | 53.869 | 1.67 | *Lemna minor* 5887 | PRJNA1026139 | SAMN37735468 | SRR26858618 |
| KS12 | BFD-Spo-MOO1 | *Spirodela polyrhiza* | 53.801 | 1.723 | *Spirodela polyrhiza* 5888 | PRJNA1026139 | SAMN37735469 | SRR26858617 |
| KS13 | BFD-Lmo-ELD | *Lemna minor* | 53.866 | 1.815 | *Lemna minor* 5889 | PRJNA1026139 | SAMN37735470 | SRR26858616 |
| KS14 | BFD-Ljp-APP | *Lemna japonica* | 53.835 | 1.711 | *Lemna japonica* 5890 | PRJNA1026139 | SAMN37735471 | SRR26858615 |
| KS15 | BFD-Ljp-BRE | *Lemna japonica* | 53.871 | 1.597 | *Lemna japonica* 5891 | PRJNA1026139 | SAMN37735472 | SRR26858614 |
| KS16 | HUL-Ltu-EAS | *Lemna turionifera* | 53.766 | 0.298 | *Lemna turionifera* 5892 | PRJNA1026139 | SAMN37735473 | SRR26858635 |
| KS17 | HUL-Ljp-PEA | *Lemna japonica* | 53.758 | 0.355 | *Lemna japonica* 5893 | PRJNA1026139 | SAMN37735474 | SRR26858634 |
| KS18 | HUL-Ljp-CRE | *Lemna japonica* | 53.783 | 0.408 | *Lemna japonica* 5894 | PRJNA1026139 | SAMN37735475 | SRR26858633 |
| KS20 | HUL-Lmu-BEV | *Lemna minuta* | 53.839 | 0.405 | *Lemna minuta* 5895 | PRJNA1026139 | SAMN37735476 | SRR26858632 |
| KS21 | HUL-Ljp-WAL | *Lemna japonica* | 53.821 | 0.482 | *Lemna japonica* 5896 | PRJNA1026139 | SAMN37735477 | SRR26858631 |
| KS22 | HUL-Ltu-NOR | *Lemna turionifera* | 53.783 | 0.642 | *Lemna turionifera* 5897 | PRJNA1026139 | SAMN37735478 | SRR26858630 |
| KS25 | YOR-Lmu-ESC | *Lemna minuta* | 53.875 | 1.034 | *Lemna minuta* 5898 | PRJNA1026139 | SAMN37735479 | SRR26858629 |
| KS27 | YOR-Lmo-BUR | *Lemna minor* | 53.996 | 1.132 | *Lemna minor* 5899 | PRJNA1026139 | SAMN37735480 | SRR26858628 |
| KS28 | YOR-Ljp-HES | *Lemna japonica* | 53.979 | 1.187 | *Lemna japonica* 5900 | PRJNA1026139 | SAMN37735481 | SRR26858627 |
| KS29 | YOR-Lmo-TAD | *Lemna minor* | 53.902 | 1.281 | *Lemna minor* 5901 | PRJNA1026139 | SAMN37735482 | SRR26858626 |
| LY01A | COR-Ljp-TRE | *Lemna japonica* | 50.407 | -5.061 | *Lemna japonica* 5902 | PRJNA1026139 | SAMN37735483 | SRR26858624 |
| LY01B | COR-Lmu-TRE | *Lemna minuta* | 50.407 | -5.061 | *Lemna minuta* 5903 | PRJNA1026139 | SAMN37735484 | SRR26858623 |
| LY02 | COR-Ljp-SHE | *Lemna japonica* | 51.003 | -3.101 | *Lemna japonica* 5904 | PRJNA1026139 | SAMN37735485 | SRR26858622 |
| LY03 | COR-Ljp-HOU | *Lemna japonica* | 49.964 | -5.196 | *Lemna japonica* 5905 | PRJNA1026139 | SAMN37735486 | SRR26858621 |
| AL01 | MID-Lmu-ALP | *Lemna minuta* | 52.64 | -1.055 | NR | PRJNA1030266 | NR | SRR27840097 |
| AL02 | MID-Lmu-ALD | *Lemna minuta* | 52.643 | -1.059 | NR | PRJNA1030266 | NR | SRR27840096 |
| AL03 | MID-Lmu-SHR | *Lemna minuta* | 52.462 | -2.211 | NR | PRJNA1030266 | NR | SRR27840083 |
| KS100 | ELG-Lmo-BUR | *Lemna minor* | 57.620 | -3.524 | NR | PRJNA1030266 | NR | SRR27840072 |

| | | | | | | | | |
|---|---|---|---|---|---|---|---|---|
| KS101 | ELG-Lmo-MAV | *Lemna minor* | 57.654 | -3.177 | NR | PRJNA1030266 | NR | SRR27840031 |
| KS104 | ELG-Lmo-LNB | *Lemna minor* | 57.628 | -3.200 | NR | PRJNA1030266 | NR | SRR27840020 |
| KS107 | ELG-Lmo-COO | *Lemna minor* | 57.652 | -3.315 | NR | PRJNA1030266 | NR | SRR27840009 |
| KS108 | ABE-Lmo-TOL | *Lemna minor* | 57.111 | -2.129 | NR | PRJNA1030266 | NR | SRR27839998 |
| KS109 | ABE-Lmo-DEE | *Lemna minor* | 57.101 | -2.235 | NR | PRJNA1030266 | NR | SRR27840053 |
| KS110 | ABE-Lmo-ALL | *Lemna minor* | 57.114 | -2.179 | NR | PRJNA1030266 | NR | SRR27840042 |
| KS111 | ABE-Lmo-HAZ | *Lemna minor* | 57.135 | -2.174 | NR | PRJNA1030266 | NR | SRR27840095 |
| KS112 | ABE-Lmo-COU | *Lemna minor* | 57.134 | -2.155 | NR | PRJNA1030266 | NR | SRR27840092 |
| KS114 | ABE-Lmo-DUT | *Lemna minor* | 57.129 | -2.106 | NR | PRJNA1030266 | NR | SRR27840091 |
| KS115 | ABE-Lmo-DON | *Lemna minor* | 57.177 | -2.119 | NR | PRJNA1030266 | NR | SRR27840090 |
| KS116 | ABE-Lmo-BLK | *Lemna minor* | 57.217 | -2.073 | NR | PRJNA1030266 | NR | SRR27840089 |
| KS117 | ABE-Lmo-DEN | *Lemna minor* | 57.152 | -2.277 | NR | PRJNA1030266 | NR | SRR27840088 |
| KS118 | ABE-Lmo-KWS | *Lemna minor* | 57.164 | -2.218 | NR | PRJNA1030266 | NR | SRR27840087 |
| KS119 | ABE-Lmo-POT | *Lemna minor* | 57.230 | -2.099 | NR | PRJNA1030266 | NR | SRR27840086 |
| KS33 | LAN-Lmo-SIL | *Lemna minor* | 54.169 | -2.799 | NR | PRJNA1030266 | NR | SRR27840085 |
| KS34 | LAN-Lmo-CAR | *Lemna minor* | 54.125 | -2.771 | NR | PRJNA1030266 | NR | SRR27840084 |
| KS38 | MID-Lmu-ELV | *Lemna minuta* | 52.895 | -1.396 | NR | PRJNA1030266 | NR | SRR27840082 |
| KS39 | MID-Lmo-LIM | *Lemna minor* | 52.811 | -1.466 | NR | PRJNA1030266 | NR | SRR27840081 |
| KS40A | MID-Ljp-CAL | *Lemna japonica* | 52.800 | -1.453 | *Lemna japonica* 5942 | PRJNA1030266 | NR | SRR27840080 |
| KS42 | HAS-Lmu-HEL | *Lemna minuta* | 50.881 | 0.582 | NR | PRJNA1030266 | NR | SRR27840079 |
| KS43 | HAS-Lmo-GIL | *Lemna minor* | 50.865 | 0.543 | NR | PRJNA1030266 | NR | SRR27840078 |
| KS44A | HAS-Lmo-LAN | *Lemna minor* | 50.798 | 0.305 | NR | PRJNA1030266 | NR | SRR27840077 |
| KS44B | HAS-Lmu-LAN | *Lemna minuta* | 50.798 | 0.305 | NR | PRJNA1030266 | NR | SRR27840076 |
| KS45 | HAS-Lmu-WIL | *Lemna minuta* | 50.801 | 0.289 | NR | PRJNA1030266 | NR | SRR27840075 |
| KS46B | HAS-Lmu/Lmo-WHE | *Lmu/Lmo* | 50.917 | 0.562 | *Lemna minor/minuta* hybrid 5943 | PRJNA1030266 | NR | SRR27840074 |
| KS47 | HAS-Lmo-BRE | *Lemna minor* | 50.943 | 0.562 | NR | PRJNA1030266 | NR | SRR27840073 |
| KS48A | HAS-Lmo-UDI | *Lemna minor* | 50.941 | 0.652 | NR | PRJNA1030266 | NR | SRR27840071 |
| KS48B | HAS-Lmu-UDI | *Lemna minuta* | 50.941 | 0.652 | NR | PRJNA1030266 | NR | SRR27840070 |
| KS49 | HAS-Lmu-MIL | *Lemna minuta* | 51.023 | 0.783 | NR | PRJNA1030266 | NR | SRR27840069 |
| KS50 | HAS-Lmo-CRO | *Lemna minor* | 50.899 | 0.520 | NR | PRJNA1030266 | NR | SRR27840068 |
| KS51A | COR-Lmo-COR | *Lemna minor* | 50.941 | 0.652 | NR | PRJNA1030266 | NR | SRR27840067 |

| | | | | | | | | |
|---|---|---|---|---|---|---|---|---|
| KS51B | COR-Lmu/Lmo-COR | *Lmu/Lmo* | 50.941 | 0.652 | *Lemna minor/minuta* hybrid 5944 | PRJNA1030266 | NR | SRR27840066 |
| KS52 | COR-Lmo-KWO | *Lemna minor* | 50.314 | -4.799 | NR | PRJNA1030266 | NR | SRR27840065 |
| KS53 | COR-Lmo-GRA | *Lemna minor* | 50.299 | -4.905 | NR | PRJNA1030266 | NR | SRR27840064 |
| KS54 | COR-Lmo-MEN | *Lemna minor* | 50.346 | -4.796 | NR | PRJNA1030266 | NR | SRR27840063 |
| KS55 | COR-Lmo-COM | *Lemna minor* | 50.334 | -4.879 | NR | PRJNA1030266 | NR | SRR27840062 |
| KS56 | COR-Lmo-TRG | *Lemna minor* | 50.355 | -4.884 | NR | PRJNA1030266 | NR | SRR27840030 |
| KS57 | COR-Lmo-TRW | *Lemna minor* | 50.291 | -4.932 | NR | PRJNA1030266 | NR | SRR27840029 |
| KS58B | COR-Lmu/Lmo-PIN | *Lmu/Lmo* | 50.342 | -4.751 | *Lemna minor/minuta* hybrid 5945 | PRJNA1030266 | NR | SRR27840028 |
| KS59 | COR-Lmo-INN | *Lemna minor* | 50.378 | -4.766 | NR | PRJNA1030266 | NR | SRR27840027 |
| KS60 | COR-Lmo-AND | *Lemna minor* | 50.358 | -4.707 | NR | PRJNA1030266 | NR | SRR27840026 |
| KS61A | COR-Lmo-HEL | *Lemna minor* | 50.283 | -4.808 | NR | PRJNA1030266 | NR | SRR27840025 |
| KS61B | COR-Lmu-HEL | *Lemna minuta* | 50.283 | -4.808 | NR | PRJNA1030266 | NR | SRR27840024 |
| KS62 | COR-Ljp-JAP | *Lemna japonica* | 50.456 | -4.998 | *Lemna japonica* 5946 | PRJNA1030266 | NR | SRR27840023 |
| KS63 | BRI-Lmu/Lmo-PUX | *Lmu/Lmo* | 51.372 | -2.851 | *Lemna minor/minuta* hybrid 5947 | PRJNA1030266 | NR | SRR27840022 |
| KS64B | BRI-Lmu-WEM | *Lemna minuta* | 51.388 | -2.836 | NR | PRJNA1030266 | NR | SRR27840021 |
| KS65A | BRI-Ljp-CLA1 | *Lemna japonica* | 51.408 | -2.805 | *Lemna japonica* 5948 | PRJNA1030266 | NR | SRR27840019 |
| KS65B | BRI-Ljp-CLA2 | *Lemna japonica* | 51.408 | -2.805 | *Lemna japonica* 5949 | PRJNA1030266 | NR | SRR27840018 |
| KS66A | BRI-Ljp-LAM | *Lemna japonica* | 51.398 | -2.853 | *Lemna japonica* 5906 | PRJNA1074359 | SAMN39856635 | SRR27937329 |
| KS66B | BRI-Lgi-LAM | *Lemna gibba* | 51.398 | -2.853 | *Lemna gibba* 5932 | PRJNA1030266 | NR | SRR27840017 |
| KS66C | BRI-Lmu-LAM | *Lemna minuta* | 51.398 | -2.853 | NR | PRJNA1030266 | NR | SRR27840016 |
| KS67A | BRI-Lmo-NAI | *Lemna minor* | 51.424 | -2.821 | NR | PRJNA1030266 | NR | SRR27840015 |
| KS67B | BRI-Lmu-NAI | *Lemna minuta* | 51.424 | -2.821 | NR | PRJNA1030266 | NR | SRR27840014 |
| KS68A | BRI-Ljp-VAL | *Lemna japonica* | 51.483 | -2.754 | *Lemna japonica* 5950 | PRJNA1030266 | NR | SRR27840013 |
| KS68B | BRI-Lmu/Lmo-VAL | *Lmu/Lmo* | 51.483 | -2.754 | *Lemna minor/minuta* hybrid 5951 | PRJNA1030266 | NR | SRR27840012 |
| KS69 | BRI-Lmu-NEW | *Lemna minuta* | 51.509 | -2.657 | NR | PRJNA1030266 | NR | SRR27840011 |
| KS70 | BRI-Ljp-MLA | *Lemna japonica* | 51.568 | -2.585 | *Lemna japonica* 5952 | PRJNA1030266 | NR | SRR27840010 |
| KS72A | NEW-Ljp-TRE | *Lemna japonica* | 51.454 | -2.607 | *Lemna japonica* 5953 | PRJNA1030266 | NR | SRR27840008 |
| KS72B | NEW-Lmu-TRE | *Lemna minuta* | 51.454 | -2.607 | NR | PRJNA1030266 | NR | SRR27840007 |
| KS73 | NEW-Lmo-LLI | *Lemna minor* | 51.570 | -2.865 | NR | PRJNA1030266 | NR | SRR27840006 |

| | | | | | | | | |
|---|---|---|---|---|---|---|---|---|
| KS74A | NEW-Ljp-NAS | *Lemna japonica* | 51.584 | -2.953 | *Lemna japonica* 5954 | PRJNA1030266 | NR | SRR27840005 |
| KS74B | NEW-Lmu/Lmo-NAS | *Lmu/Lmo* | 51.584 | -2.953 | *Lemna minor/minuta* hybrid 5955 | PRJNA1030266 | NR | SRR27840004 |
| KS75A | NEW-Ljp-CHA | *Lemna japonica* | 51.564 | -2.948 | *Lemna japonica* 5956 | PRJNA1030266 | NR | SRR27840003 |
| KS75B | NEW-Lmu-CHA | *Lemna minuta (unk)* | 51.564 | -2.948 | NR | PRJNA1030266 | NR | SRR27840002 |
| KS76A | NEW-Lgi-MAL | *Lemna gibba* | 51.564 | -2.916 | *Lemna gibba* 5933 | PRJNA1030266 | NR | SRR27840001 |
| KS76B | NEW-Lmu-MAL | *Lemna minuta* | 51.564 | -2.916 | NR | PRJNA1030266 | NR | SRR27840000 |
| KS77A | NEW-Spo-FOU | *Spirodela polyrhiza* | 51.604 | -3.011 | *Spirodela polyrhiza* 5907 | PRJNA1074359 | SAMN39856636 | SRR27937328 |
| KS77B | NEW-Lmo-FOU | *Lemna minor* | 51.604 | -3.011 | NR | PRJNA1030266 | NR | SRR27839999 |
| KS77C | NEW-Lmu-FOU | *Lemna minuta* | 51.604 | -3.011 | NR | PRJNA1030266 | NR | SRR27839997 |
| KS78A | NEW-Spo-FIV | *Spirodela polyrhiza* | 51.592 | -3.040 | *Spirodela polyrhiza* 5908 | PRJNA1074359 | SAMN39856637 | SRR27937327 |
| KS80A | NEW-Lmo-PER | *Lemna minor* | 51.667 | -3.031 | NR | PRJNA1030266 | NR | SRR27839996 |
| KS80B | NEW-Lmu-PER | *Lemna minuta* | 51.667 | -3.031 | NR | PRJNA1030266 | NR | SRR27840061 |
| KS81 | NEW-Lmo-HAW | *Lemna minor* | 51.578 | -3.001 | NR | PRJNA1030266 | NR | SRR27840060 |
| KS82A | NEW-Lgi-LLA1 | *Lemna gibba* | 51.555 | -3.020 | *Lemna gibba* 5934 | PRJNA1030266 | NR | SRR27840059 |
| KS82B | NEW-Lmo-LLA2 | *Lemna minor* | 51.555 | -3.020 | NR | PRJNA1030266 | NR | SRR27840058 |
| KS83 | GLA-Lmo-PER | *Lemna minor* | 51.555 | -3.020 | NR | PRJNA1030266 | NR | SRR27840057 |
| KS85 | GLA-Lmu/Lmo-KEL | *Lmu/Lmo* | 51.518 | -3.177 | *Lemna minor/minuta* hybrid 5957 | PRJNA1030266 | NR | SRR27840056 |
| KS88 | GLA-Lmo-MAR | *Lemna minor* | 55.772 | -3.899 | NR | PRJNA1030266 | NR | SRR27840055 |
| KS91 | GLA-Lmo-ROB | *Lemna minor* | 55.764 | -4.014 | NR | PRJNA1030266 | NR | SRR27840054 |
| KS92B | GLA-Lmu-BOG | *Lemna minuta* | 55.869 | -4.285 | NR | PRJNA1030266 | NR | SRR27840052 |
| KS95 | ELG-Lmo-SBA | *Lemna minor* | 55.878 | -4.289 | NR | PRJNA1030266 | NR | SRR27840051 |
| KS97A | ELG-Ltr-ALP | *Lemna trisulca* | 55.883 | -4.270 | *Lemna trisulca* 5935 | PRJNA1030266 | NR | SRR27840050 |
| KS97B | ELG-Lmo-ALP | *Lemna minor* | 55.894 | -4.298 | NR | PRJNA1030266 | NR | SRR27840049 |
| KS98 | ELG-Lmo-FOR | *Lemna minor* | 55.875 | -4.333 | NR | PRJNA1030266 | NR | SRR27840048 |
| KS99 | ELG-Lmo-SAN | *Lemna minor* | 55.841 | -4.206 | NR | PRJNA1030266 | NR | SRR27840047 |
| KSALL2 | BFD-Ljp-ALL2 | *Lemna japonica* | 55.889 | -4.197 | *Lemna japonica* 5958 | PRJNA1030266 | NR | SRR27840046 |
| KSAPP1 | BFD-Ltr-APP | *Lemna trisulca* | 55.843 | -4.119 | *Lemna trisulca* 5936 | PRJNA1030266 | NR | SRR27840045 |
| KSAPP2 | BFD-Spo-APP | *Spirodela polyrhiza* | 55.873 | -4.075 | *Spirodela polyrhiza* 5937 | PRJNA1030266 | NR | SRR27840044 |
| KSKEY | HUL-Ljp-KEY | *Lemna japonica* | 55.866 | -4.030 | *Lemna japonica* 5959 | PRJNA1030266 | NR | SRR27840043 |
| KSMOOR1 | BFD-Spo-MOO2 | *Spirodela polyrhiza* | 57.663 | -3.061 | *Spirodela polyrhiza* 5938 | PRJNA1030266 | NR | SRR27840041 |

| | | | | | | | | |
|---|---|---|---|---|---|---|---|---|
| KSMOOR2 | BFD-Ljp-MOO | *Lemna japonica* | 57.663 | -3.045 | *Lemna japonica* 5960 | PRJNA1030266 | NR | SRR27840040 |
| KSNOR1 | HUL-Ltr-NOR | *Lemna trisulca* | 57.630 | -3.414 | *Lemna trisulca* 5939 | PRJNA1030266 | NR | SRR27840039 |
| KSNUF3 | BFD-Lmo-NUF3 | *Lemna minor* | 57.609 | -3.593 | *Lemna minor* 5909 | PRJNA1074359 | SAMN39856638 | SRR27937326 |
| KSSEL1 | YOR-Lgi-SEL | *Lemna gibba* | 57.602 | -3.607 | *Lemna gibba* 5940 | PRJNA1030266 | NR | SRR27840038 |
| KSYEAD1 | BFD-Ltr-YEA | *Lemna trisulca* | 53.869 | 1.670 | *Lemna trisulca* 5941 | PRJNA1030266 | NR | SRR27840037 |
| MP01 | NCA-Ljp-MPO | *Lemna japonica* | 51.015 | -1.473 | *Lemna japonica* 5961 | PRJNA1030266 | NR | SRR27840094 |

| **Clone** | **Site code** | **Species** | **Latitude** | **Longitude** | **Registered clone** | **SRA project** | **SRA sample** | **SRA clone** |
|---|---|---|---|---|---|---|---|---|
| *L. japonica* 9250 | WW-Lja9250 | *Lemna japonica* | | | *Lemna japonica* 9250 | PRJNA1030266 | NR | SRR27840036 |
| *L. minuta* 9260 | WW-Lmu9260 | *Lemna minuta* | | | *Lemna minuta* 9260 | PRJNA1030266 | NR | SRR27840035 |
| *L. punctata* 0049 | WW-Lpu0049 | *Lemna punctata* | | | *Lemna punctata* 0049 | PRJNA1030266 | NR | SRR27840034 |
| *L. trisulca* 7192 | WW-Ltr7192 | *Lemna trisulca* | | | *Lemna trisulca* 7192 | PRJNA1030266 | NR | SRR27840033 |
| *L. yungensis* 9208 | WW-Lyu9208 | *Lemna yungensis* | | | *Lemna yungensis* 9208 | PRJNA1030266 | NR | SRR27840032 |
| *S. intermedia* 9394 | WW-Sin9394 | *Spirodela intermedia* | | | *Spirodela intermedia* 9394 | PRJNA1030266 | NR | SRR27840093 |
| *L. japonica* 7123 | WW-Lja7123 | *Lemna japonica* | | | *Lemna japonica* 7123 | | | |
| *L. minor* 7295 | WW-Lmo7295 | *Lemna minor* | | | *Lemna minor* 7295 | | | |
| *L. minor* 8389 | WW-Lmo8389 | *Lemna minor* | | | *Lemna minor* 8389 | | | |
| *L.minuta* 6600 | WW-Lmu6600 | *Lemna minuta* | | | *Lemna minuta* 6600 | | | |

Registered clones refers to new accession submission on ruduckweeds.org. SRA refers to submission of genomes on Short read archive (SRA) with project, sample and individual codes for new accessions. Blue colouration refers to clones already characterised and registered. *NR - not released.

**Table S2B.** Genomes of duckweed clones downloaded from the Short Read Archive (SRA) and included in the genomic pipeline. Clones with < 5% coverage were removed from further analysis indicated in red.

| Clone | SRA code | Number of reads | Percentage covered | Mean coverage |
|---|---|---|---|---|
| *L. minor* 7016 | SRR10958777 | 8088768 | 86.04 | 55.7 |
| *L. minor* 5500 | SRR10958800 | 5970720 | 86.81 | 40.1 |
| *L. turionifera* 6002 | SRR23943402 | 1335194 | 46.41 | 7.27 |
| *L. gibba* 131 | SRR074103 | 991234 | 14.71 | 3.02 |
| *L. turionifera* 9434 | SRR8291590 | 22769 | 1.495 | 0.117 |
| *L. minuta* 9484 | SRR8291594 | 8920 | 0.1448 | 0.0371 |
| *L. minuta* 6717 | SRR8291596 | 8571 | 0.3825 | 0.034 |
| *L. minuta* 7612 | SRR8291595 | 3745 | 0.1921 | 0.0151 |
| *L. minuta* 9581 | SRR8291593 | 848 | 0.08327 | 0.00342 |
| *S. polyrhiza* 9504 | SRR11472010 | 45 | 0.01389 | 0.000146 |
| *S. intermedia* 8410 | ERR3957957 | 0 | 0 | 0 |

**Table S2C**. Genomes of new UK duckweed accessions and newly sequenced clones included in the genomic pipeline.

| Accession | Site code | Species | Number of reads | Percentage covered | Mean coverage |
|---|---|---|---|---|---|
| KS02 | YOR-Lmo-BIS | *Lemna minor* | 792908 | 77.39 | 5.58 |
| KS03 | BFD-Ljp-ALL1 | *Lemna japonica* | 1642080 | 83.25 | 11.1 |
| KS04 | YOR-Ljp-SEL | *Lemna japonica* | 2931134 | 85.41 | 19.6 |
| KS06A | BFD-Lmu-NUF1 | *Lemna minuta* | 523797 | 10.35 | 1.53 |
| KS06B | BFD-Lmu-NUF2 | *Lemna minuta* | 310907 | 8.41 | 0.98 |
| KS09 | BFD-Lmo-YEA | *Lemna minor* | 1156648 | 79.24 | 8.52 |
| KS12 | BFD-Spo-MOO1 | *Spirodela polyrhiza* | 757884 | 9.833 | 1.98 |
| KS13 | BFD-Lmo-ELD | *Lemna minor* | 1376941 | 81.05 | 10.2 |
| KS14 | BFD-Ljp-APP | *Lemna japonica* | 3021601 | 85.79 | 21.2 |
| KS15 | BFD-Ljp-BRE | *Lemna japonica* | 1479101 | 78.65 | 10 |
| KS16 | HUL-Ltu-EAS | *Lemna turionifera* | 1081370 | 44.87 | 5.85 |

| | | | | | |
|---|---|---|---|---|---|
| KS17 | HUL-Ljp-PEA | *Lemna japonica* | 1218293 | 81.99 | 8.21 |
| KS18 | HUL-Ljp-CRE | *Lemna japonica* | 1097315 | 81.09 | 7.35 |
| KS20 | HUL-Lmu-BEV | *Lemna minuta* | 347620 | 8.397 | 1.1 |
| KS21 | HUL-Ljp-WAL | *Lemna japonica* | 1256535 | 79.44 | 8.49 |
| KS22 | HUL-Ltu-NOR | *Lemna turionifera* | 3241725 | 53.42 | 17.1 |
| KS25 | YOR-Lmu-ESC | *Lemna minuta* | 307614 | 8.329 | 0.97 |
| KS27 | YOR-Lmo-BUR | *Lemna minor* | 1303766 | 79.93 | 9.68 |
| KS28 | YOR-Ljp-HES | *Lemna japonica* | 1386609 | 82.31 | 9.33 |
| KS29 | YOR-Lmo-TAD | *Lemna minor* | 2698291 | 82.75 | 19.9 |
| LY01A | COR-Ljp-TRE | *Lemna japonica* | 1520377 | 82.41 | 10.3 |
| LY01B | COR-Lmu-TRE | *Lemna minuta* | 431435 | 17.74 | 1.35 |
| LY02 | COR-Ljp-SHE | *Lemna japonica* | 1517640 | 82.82 | 10.3 |
| LY03 | COR-Ljp-HOU | *Lemna japonica* | 1259293 | 82.07 | 8.5 |
| AL01 | MID-Lmu-ALP | *Lemna minuta* | 422763 | 14.06 | 1.37 |
| AL02 | MID-Lmu-ALD | *Lemna minuta* | 822894 | 14.87 | 2.39 |
| AL03 | MID-Lmu-SHR | *Lemna minuta* | 488595 | 10.95 | 1.31 |
| KS100 | ELG-Lmo-BUR | *Lemna minor* | 4631474 | 85.03 | 32.9 |
| KS101 | ELG-Lmo-MAV | *Lemna minor* | 1277761 | 79.99 | 8.92 |
| KS104 | ELG-Lmo-LNB | *Lemna minor* | 4572537 | 85.29 | 32.6 |
| KS107 | ELG-Lmo-COO | *Lemna minor* | 1565440 | 81.24 | 11.5 |
| KS108 | ABE-Lmo-TOL | *Lemna minor* | 2318986 | 82.2 | 16.5 |
| KS109 | ABE-Lmo-DEE | *Lemna minor* | 1414233 | 80.65 | 10.4 |
| KS110 | ABE-Lmo-ALL | *Lemna minor* | 2433634 | 83.78 | 17.2 |
| KS111 | ABE-Lmo-HAZ | *Lemna minor* | 2567984 | 84.07 | 18.4 |
| KS112 | ABE-Lmo-COU | *Lemna minor* | 2991000 | 83.97 | 21.2 |
| KS114 | ABE-Lmo-DUT | *Lemna minor* | 1426163 | 80.84 | 10.4 |
| KS115 | ABE-Lmo-DON | *Lemna minor* | 2522849 | 82.2 | 14 |
| KS116 | ABE-Lmo-BLK | *Lemna minor* | 1705583 | 81.43 | 12.4 |
| KS117 | ABE-Lmo-DEN | *Lemna minor* | 476228 | 69.11 | 3.31 |
| KS118 | ABE-Lmo-KWS | *Lemna minor* | 2431844 | 82.86 | 17.4 |
| KS119 | ABE-Lmo-POT | *Lemna minor* | 2685678 | 82.97 | 19.1 |

| | | | | | |
|---|---|---|---|---|---|
| KS33 | LAN-Lmo-SIL | *Lemna minor* | 4875223 | 84.66 | 34.9 |
| KS34 | LAN-Lmo-CAR | *Lemna minor* | 2629542 | 82.95 | 18.8 |
| KS38 | MID-Lmu-ELV | *Lemna minuta* | 468179 | 10.76 | 0.981 |
| KS39 | MID-Lmo-LIM | *Lemna minor* | 3126447 | 83.73 | 22.3 |
| KS40A | MID-Ljp-CAL | *Lemna japonica* | 2437207 | 84.63 | 15.8 |
| KS42 | HAS-Lmu-HEL | *Lemna minuta* | 478863 | 10.61 | 1.26 |
| KS43 | HAS-Lmo-GIL | *Lemna minor* | 133422 | 38.44 | 0.948 |
| KS44A | HAS-Lmo-LAN | *Lemna minor* | 1539729 | 80.97 | 11.2 |
| KS44B | HAS-Lmu-LAN | *Lemna minuta* | 387567 | 9.72 | 0.997 |
| KS45 | HAS-Lmu-WIL | *Lemna minuta* | 512705 | 11.29 | 1.25 |
| KS46B | HAS-Lmu/Lmo-WHE | *Lmu/Lmo* | 638159 | 27.42 | 2.1 |
| KS47 | HAS-Lmo-BRE | *Lemna minor* | 2934396 | 83.88 | 21 |
| KS48A | HAS-Lmo-UDI | *Lemna minor* | 1415382 | 80.9 | 10.4 |
| KS48B | HAS-Lmu-UDI | *Lemna minuta* | 475833 | 10.7 | 1.26 |
| KS49 | HAS-Lmu-MIL | *Lemna minuta* | 2181308 | 16.26 | 6.04 |
| KS50 | HAS-Lmo-CRO | *Lemna minor* | 5379121 | 84.93 | 38.5 |
| KS51A | COR-Lmo-COR | *Lemna minor* | 1232844 | 80.13 | 8.95 |
| KS51B | COR-Lmu/Lmo-COR | *Lmu/Lmo* | 770044 | 31.26 | 2.53 |
| KS52 | COR-Lmo-KWO | *Lemna minor* | 1968260 | 82.06 | 14 |
| KS53 | COR-Lmo-GRA | *Lemna minor* | 1111904 | 79.85 | 8.19 |
| KS54 | COR-Lmo-MEN | *Lemna minor* | 1680320 | 81.39 | 12.2 |
| KS55 | COR-Lmo-COM | *Lemna minor* | 2743462 | 82.8 | 19.4 |
| KS56 | COR-Lmo-TRG | *Lemna minor* | 2965827 | 83.23 | 21.2 |
| KS57 | COR-Lmo-TRW | *Lemna minor* | 292621 | 61 | 2.1 |
| KS58B | COR-Lmu/Lmo-PIN | *Lmu/Lmo* | 2382832 | 48.82 | 7.17 |
| KS59 | COR-Lmo-INN | *Lemna minor* | 3084228 | 83.49 | 22 |
| KS60 | COR-Lmo-AND | *Lemna minor* | 2448660 | 82.8 | 17.5 |
| KS61A | COR-Lmo-HEL | *Lemna minor* | 1262225 | 80.47 | 9.28 |
| KS61B | COR-Lmu-HEL | *Lemna minuta* | 896527 | 12.23 | 2.62 |
| KS62 | COR-Ljp-JAP | *Lemna japonica* | 4375540 | 86.27 | 28.3 |
| KS63 | BRI-Lmu/Lmo-PUX | *Lmu/Lmo* | 517792 | 69.35 | 3.28 |

| | | | | | |
|---|---|---|---|---|---|
| KS64B | BRI-Lmu-WEM | *Lemna minuta* | 612677 | 10.88 | 1.76 |
| KS65A | BRI-Ljp-CLA1 | *Lemna japonica* | 1530601 | 82.31 | 10.1 |
| KS65B | BRI-Ljp-CLA2 | *Lemna japonica* | 1148531 | 80.93 | 7.24 |
| KS66A | BRI-Ljp-LAM | *Lemna japonica* | 1545977 | 82.11 | 10.1 |
| KS66B | BRI-Lgi-LAM | *Lemna gibba* | 1595583 | 78.26 | 8.37 |
| KS66C | BRI-Lmu-LAM | *Lemna minuta* | 450640 | 10.31 | 1.18 |
| KS67A | BRI-Lmo-NAI | *Lemna minor* | 5566145 | 85.18 | 39.5 |
| KS67B | BRI-Lmu-NAI | *Lemna minuta* | 711880 | 21.92 | 2.1 |
| KS68A | BRI-Ljp-VAL | *Lemna japonica* | 2207580 | 83.95 | 14.3 |
| KS68B | BRI-Lmu/Lmo-VAL | *Lmu/Lmo* | 874414 | 34.97 | 2.87 |
| KS69 | BRI-Lmu-NEW | *Lemna minuta* | 107502 | 4.84 | 0.249 |
| KS70 | BRI-Ljp-MLA | *Lemna japonica* | 1142971 | 81.1 | 7.54 |
| KS72A | NEW-Ljp-TRE | *Lemna japonica* | 1975400 | 83.5 | 12.9 |
| KS72B | NEW-Lmu-TRE | *Lemna minuta* | 694894 | 12.55 | 1.99 |
| KS73 | NEW-Lmo-LLI | *Lemna minor* | 2339786 | 82.41 | 16.7 |
| KS74A | NEW-Ljp-NAS | *Lemna japonica* | 1622319 | 82.74 | 10.8 |
| KS74B | NEW-Lmu/Lmo-NAS | *Lmu/Lmo* | 154863 | 14.14 | 0.5 |
| KS75A | NEW-Ljp-CHA | *Lemna japonica* | 3081153 | 85.17 | 20 |
| KS75B | NEW-Lmu-CHA | *Lemna minuta (unk)* | 0 | 0 | 0 |
| KS76A | NEW-Lgi-MAL | *Lemna gibba* | 1907267 | 81.51 | 13.8 |
| KS76B | NEW-Lmu-MAL | *Lemna minuta* | 540172 | 10.62 | 1.39 |
| KS77A | NEW-Spo-FOU | *Spirodela polyrhiza* | 1108382 | 9.252 | 3.13 |
| KS77B | NEW-Lmo-FOU | *Lemna minor* | 1377004 | 80.58 | 10 |
| KS77C | NEW-Lmu-FOU | *Lemna minuta* | 501751 | 10.68 | 1.31 |
| KS78A | NEW-Spo-FIV | *Spirodela polyrhiza* | 433303 | 7.868 | 0.981 |
| KS80A | NEW-Lmo-PER | *Lemna minor* | 1635862 | 81 | 11.7 |
| KS80B | NEW-Lmu-PER | *Lemna minuta* | 442018 | 10.4 | 1.11 |
| KS81 | NEW-Lmo-HAW | *Lemna minor* | 2124835 | 82.71 | 15 |
| KS82A | NEW-Lgi-LLA1 | *Lemna gibba* | 158496 | 18.32 | 0.539 |
| KS82B | NEW-Lmo-LLA2 | *Lemna minor* | 6588240 | 85.58 | 47 |
| KS83 | GLA-Lmo-PER | *Lemna minor* | 1231037 | 80.29 | 9.06 |

| | | | | | |
|---|---|---|---|---|---|
| KS85 | GLA-Lmu/Lmo-KEL | *Lmu/Lmo* | 1702567 | 61.17 | 5.92 |
| KS88 | GLA-Lmo-MAR | *Lemna minor* | 2008275 | 81.87 | 14.3 |
| KS91 | GLA-Lmo-ROB | *Lemna minor* | 1334701 | 80.4 | 9.76 |
| KS92B | GLA-Lmu-BOG | *Lemna minuta* | 548954 | 10.72 | 1.5 |
| KS95 | ELG-Lmo-SBA | *Lemna minor* | 3365699 | 83.7 | 24.2 |
| KS97A | ELG-Ltr-ALP | *Lemna trisulca* | 1800983 | 54.7 | 9.33 |
| KS97B | ELG-Lmo-ALP | *Lemna minor* | 2545383 | 82.56 | 18.2 |
| KS98 | ELG-Lmo-FOR | *Lemna minor* | 3199862 | 83.85 | 22.8 |
| KS99 | ELG-Lmo-SAN | *Lemna minor* | 2914512 | 83.91 | 20.6 |
| KSALL2 | BFD-Ljp-ALL2 | *Lemna japonica* | 2570836 | 84.94 | 16.8 |
| KSAPP1 | BFD-Ltr-APP | *Lemna trisulca* | 4168011 | 54.81 | 21.8 |
| KSAPP2 | BFD-Spo-APP | *Spirodela polyrhiza* | 301646 | 6.988 | 0.663 |
| KSKEY | HUL-Ljp-KEY | *Lemna japonica* | 2756002 | 85.16 | 17.9 |
| KSMOOR1 | BFD-Spo-MOO2 | *Spirodela polyrhiza* | 430447 | 7.667 | 1.05 |
| KSMOOR2 | BFD-Ljp-MOO | *Lemna japonica* | 1692380 | 82.96 | 11.1 |
| KSNOR1 | HUL-Ltr-NOR | *Lemna trisulca* | 1809715 | 72.17 | 9.87 |
| KSNUF3 | BFD-Lmo-NUF3 | *Lemna minor* | 1306180 | 80.57 | 9.42 |
| KSSEL1 | YOR-Lgi-SEL | *Lemna gibba* | 991069 | 73.64 | 5.13 |
| KSYEAD1 | BFD-Ltr-YEA | *Lemna trisulca* | 1936634 | 50.48 | 10.2 |
| MP01 | NCA-Ljp-MPO | *Lemna japonica* | 1329520 | 81.86 | 8.84 |

| **Clone** | **Site code** | **Species** | **Number of reads** | **Percentage covered** | **Mean coverage** |
|---|---|---|---|---|---|
| *L. japonica* 9250 | WW-Lja9250 | *Lemna japonica* | 3200303 | 84.32 | 21.3 |
| *L. minuta* 9260 | WW-Lmu9260 | *Lemna minuta* | 526680 | 10.8 | 1.38 |
| *L. punctata* 0049 | WW-Lpu0049 | *Lemna punctata* | 510974 | 23.93 | 1.75 |
| *L. trisulca* 7192 | WW-Ltr7192 | *Lemna trisulca* | 1374501 | 45.81 | 7.09 |
| *L. yungensis* 9208 | WW-Lyu9208 | *Lemna yungensis* | 1130919 | 14.93 | 2.94 |
| *S. intermedia* 9394 | WW-Sin9394 | *Spirodela intermedia* | 274124 | 6.735 | 0.562 |
| *L. japonica* 7123 | WW-Lja7123 | *Lemna japonica* | 2844367 | 83.47 | 17.9 |
| *L. minor* 7295 | WW-Lmo7295 | *Lemna minor* | 3996123 | 85.25 | 26.7 |
| *L. minor* 8389 | WW-Lmo8389 | *Lemna minor* | 5044832 | 86.09 | 33.8 |
| *L.minuta* 6600 | WW-Lmu6600 | *Lemna minuta* | 1164164 | 14.32 | 2.87 |

**Table S3.** UK duckweed species show differences between tissue concentrations of elements grown in replete N-medium. Four UK duckweed species are included in the table rows and columns, with the intersections marking significant differences between species. Red indicates the species in the row has higher accumulation, green indicates the species in the column has higher accumulation of that element. Kruskal-Wallis and Dunn's post-hoc tests were used with significance set at P=<0.05. All non-significant relationships have been removed for clarity.

A.

| Mg | *L. minor* | *L. japonica* | *L. minuta* |
|---|---|---|---|
| *L. japonica* | | | |
| *L. minuta* | 0.0000 | 0.0000 | |
| *Lmu/Lmo* | 0.0027 | 0.0000 | |

B.

| K | *L. minor* | *L. japonica* | *L. minuta* |
|---|---|---|---|
| *L. japonica* | 0.0065 | | |
| *L. minuta* | | 0.0066 | |
| *Lmu/Lmo* | | 0.0038 | |

C.

| Ca | *L. minor* | *L. japonica* | *L. minuta* |
|---|---|---|---|
| *L. japonica* | | | |
| *L. minuta* | | 0.0402 | |
| *Lmu/Lmo* | | | |

D.

| Si | *L. minor* | *L. japonica* | *L. minuta* |
|---|---|---|---|
| *L. japonica* | 0.0127 | | |
| *L. minuta* | | | |
| *Lmu/Lmo* | | | |

E.

| Mn | *L. minor* | *L. japonica* | *L. minuta* |
|---|---|---|---|
| *L. japonica* | | | |
| *L. minuta* | | | |
| *Lmu/Lmo* | | 0.0187 | |

F.

| Mo | *L. minor* | *L. japonica* | *L. minuta* |
|---|---|---|---|
| *L. japonica* | 0.0102 | | |
| *L. minuta* | | | |
| *Lmu/Lmo* | | | |

G.

| As | *L. minor* | *L. japonica* | *L. minuta* |
|---|---|---|---|
| *L. japonica* | 0.05 | | |
| *L. minuta* | | | |
| *Lmu/Lmo* | | | |

**Table S4.** Duckweed species show differences between typical elemental concentrations found in water habitats. Four UK duckweed species are included in the table rows and columns, with the intersections marking significant differences between species. Red indicates the species in the row has higher presence of that element in its water habitats, green indicates the species in the column has higher presence in its water habitats. Kruskal-Wallis and Dunn's post-hoc tests were used with significance set at P=<0.05. All non-significant relationships have been removed for clarity.

A.

| Mg | *L. minor* | *L. japonica* | *L. minuta* |
|---|---|---|---|
| *L. japonica* | | | |
| *L. minuta* | 0.046 | | |
| *Lmu/Lmo* | | | |

B.

| P | *L. minor* | *L. japonica* | *L. minuta* |
|---|---|---|---|
| *L. japonica* | 0.0016 | | |
| *L. minuta* | | | |
| *Lmu/Lmo* | | | |

C.

| S | *L. minor* | *L. japonica* | *L. minuta* |
|---|---|---|---|
| *L. japonica* | | | |
| *L. minuta* | 0.0194 | | |
| *Lmu/Lmo* | | | |

D

| Ca | *L. minor* | *L. japonica* | *L. minuta* |
|---|---|---|---|
| *L. japonica* | 0.0019 | | |
| *L. minuta* | 0.0003 | | |
| *Lmu/Lmo* | | | |

E

| B | *L. minor* | *L. japonica* | *L. minuta* |
|---|---|---|---|
| *L. japonica* | 0.0081 | | |
| *L. minuta* | 0.0039 | | |
| *Lmu/Lmo* | | | |

F

| Mo | *L. minor* | *L. japonica* | *L. minuta* |
|---|---|---|---|
| *L. japonica* | 0.001 | | |
| *L. minuta* | 0.0163 | | |
| *Lmu/Lmo* | | | |

G

| Sr | *L. minor* | *L. japonica* | *L. minuta* |
|---|---|---|---|
| *L. japonica* | 0.0156 | | |
| *L. minuta* | 0.0032 | | |
| *Lmu/Lmo* | | | |

H

| As | *L. minor* | *L. japonica* | *L. minuta* |
|---|---|---|---|
| *L. japonica* | 0.0007 | | |
| *L. minuta* | | | |
| *Lmu/Lmo* | | | |

I

| Al | *L. minor* | *L. japonica* | *L. minuta* |
|---|---|---|---|
| *L. japonica* | 0.0033 | | |
| *L. minuta* | 0.044 | | |
| *Lmu/Lmo* | | | |

**Table S5.** Range of internal duckweed concentrations of eight elements grown in standard replete nutrient conditions. A subset of macronutrients and toxic elemental concentrations measured in 116 duckweed accessions (in mg/kg) by ICP-MS.

| (mg/kg) | Mg | Ca | S | K | Si | Mn | Fe | Pb |
|---|---|---|---|---|---|---|---|---|
| Min conc. | 2097.29 | 2981.19 | 443.97 | 3660.63 | 0.00 | 3.54 | 70.56 | 0.00 |
| Max conc. | 9396.18 | 18703 | 21430 | 11038 | 1045.91 | 2494.01 | 1139.76 | 12.98 |
| **Fold change** | 3.5x | 5.3x | 43.7x | 29.2x | 11.0x | 703x | 15.2x | 191x |

**Table S6.** Spatial variation of eight elements found in UK water sites. A subset of macronutrients and potentially toxic elements measured in 100 UK duckweed water sampling sites. Concentrations are measured in μg/L by ICP-MS and given to 2 dp. Limits for Mn, Fe and Pb were obtained from drinking water standards (Rohlich, 1979).

| (μg/L) | Mg | Ca | S | K | Si | Mn | Fe | Pb |
|---|---|---|---|---|---|---|---|---|
| Min conc. | 1286.12 | 8.58 | 43.20 | 633.68 | 56.38 | 0.32 | 5.08 | 0.01 |
| Max conc. | 74149 | 231550 | 101887 | 305299 | 15731 | 2178 | 11610 | 6.44 |
| **Limit** | | | | | | **50** | **200** | **10** |
| **Fold change** | 56x | 2285x | 2357x | 2285x | 278x | 6807x | 2285x | 437x |

**Table S7.** Seasonal variation of eight elements in UK water sites. * A subset of macronutrients and potentially toxic elements measured in 19 UK water sites at four seasonal time points. Concentrations are measured in µg/L by ICP-MS and given to 2 dp. Limits for Mn, Fe and Pb were obtained from drinking water standards (Rohlich, 1979).

| **(µg/L)** | **Mg** | **Ca** | **S** | **K** | **Si** | **Mn** | **Fe** | **Pb** |
|---|---|---|---|---|---|---|---|---|
| Min conc. | 1546.88 | 8.58 | 43.20 | 680.39 | 56.38 | 0.07 | 0.56 | 0.03 |
| Max conc. | 43199 | 142369 | 101887 | 63130 | 13229 | 1122.27 | 3123.74 | 4.79 |
| **Limit** | | | | | | **50** | **200** | **10** |
| **Fold change** | 28x | 16586x | 2357x | 93x | 235x | 14470x | 5584x | 158x |
